# The VT GAL4, LexA, and split-GAL4 driver line collections for targeted expression in the *Drosophila* nervous system

**DOI:** 10.1101/198648

**Authors:** Laszlo Tirian, Barry J. Dickson

## Abstract

In studying the cellular interactions within complex tissues, it is extremely valuable to be able to reproducibly and flexibly target transgene expression to restricted subsets of cells. This approach is particularly valuable in studying the nervous system, with its bewildering diversity of neuronal cell types. We report here the generation of over 18,000 driver lines (the VT collection) that exploit the GAL4, LexA, and split-GAL4 systems to express transgenes in distinct and highly specific cell types in *Drosophila*. We document the expression patterns of over 14,000 of these lines in the adult male brain.

## Results and Discussion

The GAL4/UAS system (Brand and Perrimon, 1993) has proven to be a robust and versatile method for targeted transgene expression in *Drosophila*. Large collections of GAL4 lines have previously been generated (Gohl et al., 2011; Hayashi et al., 2002; Jenett et al., 2012). Complementing these resources, we generated a total of 8,166 enhancer-GAL4 lines (the “VT” or “Vienna Tiles” lines) using the strategy described by Pfeiffer et al. (2008). Each of these VT GAL4 driver lines contains a predicted ∼2kb enhancer region, or in some cases a promoter region, cloned into the pBPGUw vector or its promoterless equivalent pBPGw. These GAL4 driver constructs were inserted by site-specific recombination into the *attP2* landing site on chromosome arm 3L (Groth et al., 2004; Table 1).

**Table 1.** Numbers of VT lines generated and images.

| Driver and landing site | Lines | Imaged lines |
| --- | --- | --- |
| GAL4 in attP2 | 8166 | 8081 |
| LexAGAD in attP2 | 446 | 438 |
| LexAp65 in attP2 | 658 | 657 |
| LexAGAD in attP40 | 2240 | 1694 |
| LexAp65 in attP40 | 280 | 265 |
| ZpGDBD in attP2 | 2736 | 1488 |
| p65ADZp in attP2 | 732 | 728 |
| p65ADZp in attP40 | 2094 | 1284 |

The expression pattern of 7,705 VT GAL4 driver lines in the embryo has been determined using in situ hybridization to detect the *GAL4* mRNA (Kvon et al., 2014). In this work, we used a *UAS-mCD8-GFP* reporter and confocal imaging to reveal the expression patterns of 8,081 VT GAL4 lines in confocal images of adult male brains. Most of these lines showed reproducible expression in restricted subsets of neuronal cell types. Approximately 3000 lines that were considered most useful for neurobiological experiments were selected for recloning into LexA and split-GAL4 vectors.

The LexA/lexAop system (Lai and Lee, 2006) can be used in conjunction with the GAL4/UAS system to independently target distinct transgenes to two distinct cell subsets. In neurobiological experiments, for example, a researcher often wishes to optogenetically activate one population of neurons while silencing or imaging activity in another population. To enable such experiments, we transferred many of the more useful enhancers and promoters into LexA vectors (Pfeiffer et al., 2010), generating a total of 3,624 VT LexA lines (Table 1). We used vectors coding for LexA fusion proteins in which the LexA DNA-binding domain is coupled to either a GAL4 or a human p65 activation domain (Pfeiffer et al., 2010), thus providing both GAL80-suppressible and GAL80-insensitive LexA variants. These VT LexA constructs were inserted into either *attP2* (Groth et al., 2004), *attP40* on 2L (Markstein et al., 2008), or both sites (Table 1). Insertions in the *attP2* site more reliably recapitulate the expression pattern of the original VT GAL4 lines, but are subject to transvection when paired with VT GAL4 lines also in *attP2* (Mellert and Truman, 2012). The insertions in *attP40* avoid transvection with any enhancer-GAL4 lines in *attP2*, with which they can also be combined in doubly homozygous stocks. The expression patterns of 3,054 VT LexA lines in adult male brains were determined using a *lexAop-myr-GFP* reporter (Table 1).

The VT GAL4 and LexA driver lines, like the analogous lines generated at Janelia (Jenett et al., 2012), rarely restrict expression to a single neuronal cell type. We therefore inserted the most useful enhancers and promoters into the ZpGAL4DBD and p65ADZp split-GAL4 vectors for intersectional targeting (Luan et al., 2006; Pfeiffer et al., 2010). In this system, neither the ZpGAL4DBD nor the p65ADZp hemidriver alone is sufficient to drive expression of a UAS transgene; only when the two are coexpressed in the same cell is a functional GAL4 reconstituted to drive transgene expression. We generated 2,736 VT ZpGAL4DBD lines, all inserted in *attP2*, and 2,826 VT p65ADZp lines, approximately one third of which were inserted in *attP2* and the rest in *attP40* (Table 1). To determine the expression patterns of individual hemidriver lines, they were combined with a pan-neuronal hemidriver (either *elav-p65ADZp* or *elav-ZpGAL4DBD*) and *UAS-mCDB8-GFP* for confocal imaging of male brains. Images were acquired for 1,488 VT ZpGAL4DBD lines and 2,012 VT p65ADZp lines (Table 1).

Our collections of VT GAL4, LexA, and split-GAL4 driver lines complement similar collections generated at Janelia (Dionne et al., 2017; Jenett et al., 2012), with which they are fully compatible. In our experience, the many thousand split-GAL4 hemidriver lines available in these collections ensure that, more often than not, multiple independent intersections can be obtained for any desired neuronal cell type. The critical factor in the success of any such endeavor is the ability to efficiently identify the useful intersections from the several million potential DBD and AD combinations, a task greatly facilitated by the image databases of parental GAL4 lines and, in the case of the VT lines, also the split-GAL4 hemidriver lines.

Our data set of 41,492 confocal images of male brains from 14,635 lines can be accessed online at Virtual Fly Brain (www.virtualflybrain.org) (Milyaev et al., 2012). Many of the VT GAL4 and LexA lines have also been independently imaged in female brains and ventral nerve cords at Janelia (www.janelia.org/gal4-gen1). Approximately 1000 of the most useful VT GAL4 lines are available from the Vienna Drosophila Resource Center (www.vdrc.at). Almost all of the split-GAL4 hemidriver lines, together with a small number of additional VT hemidriver lines subsequently generated at Janelia, are available from the Bloomington Drosophila Stock Center (Table 2, www.flystocks.bio.indiana.edu).

**Table 2.**
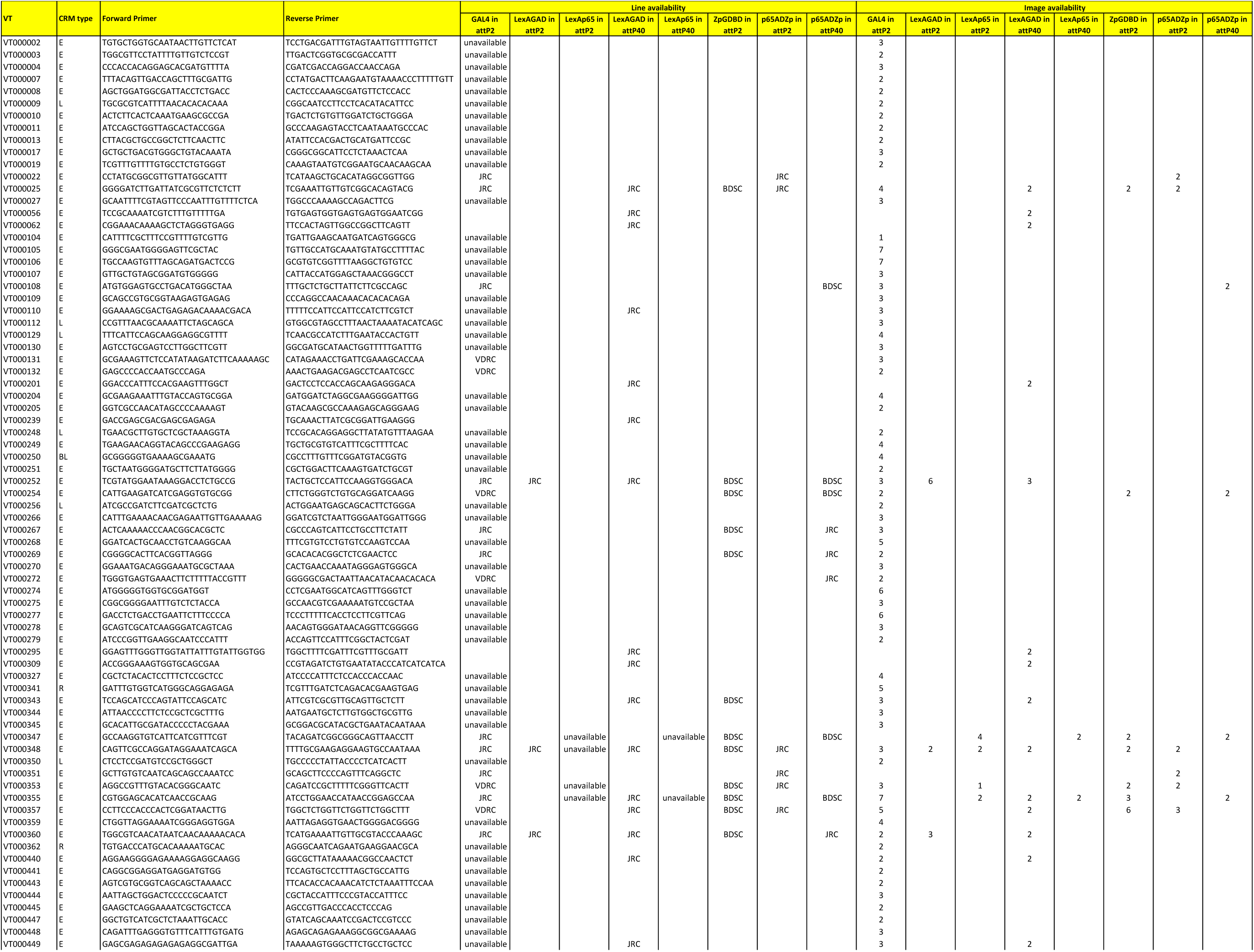
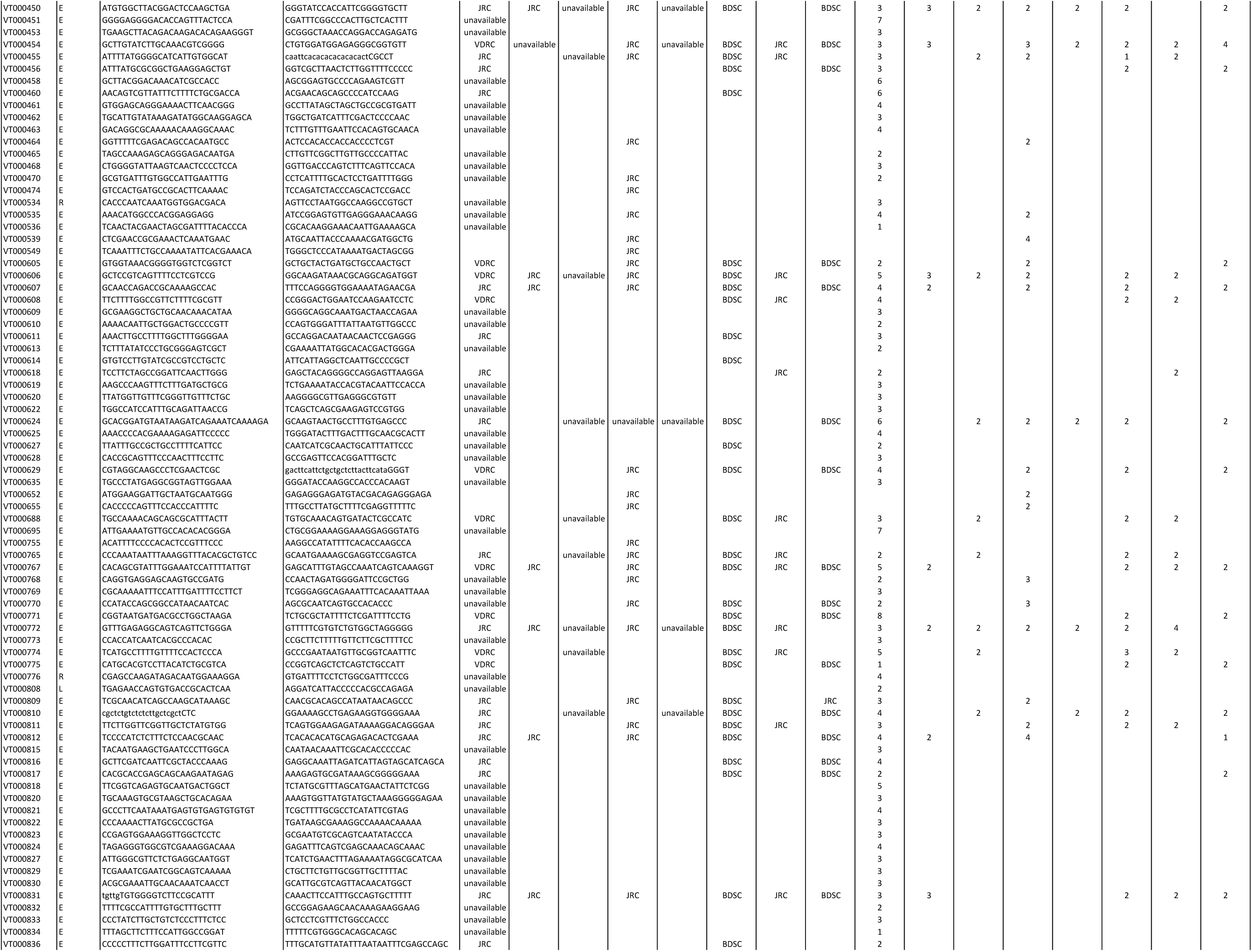
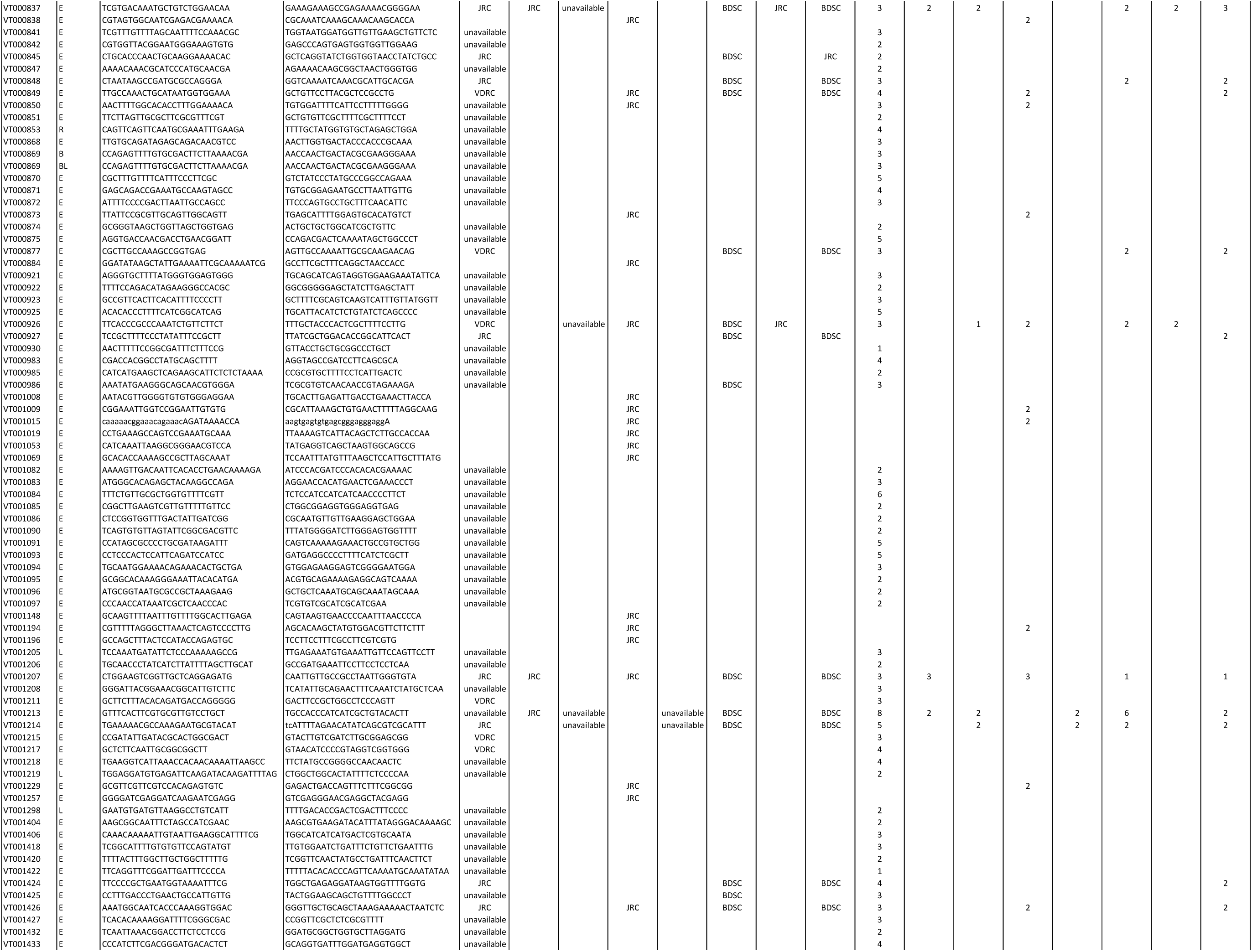
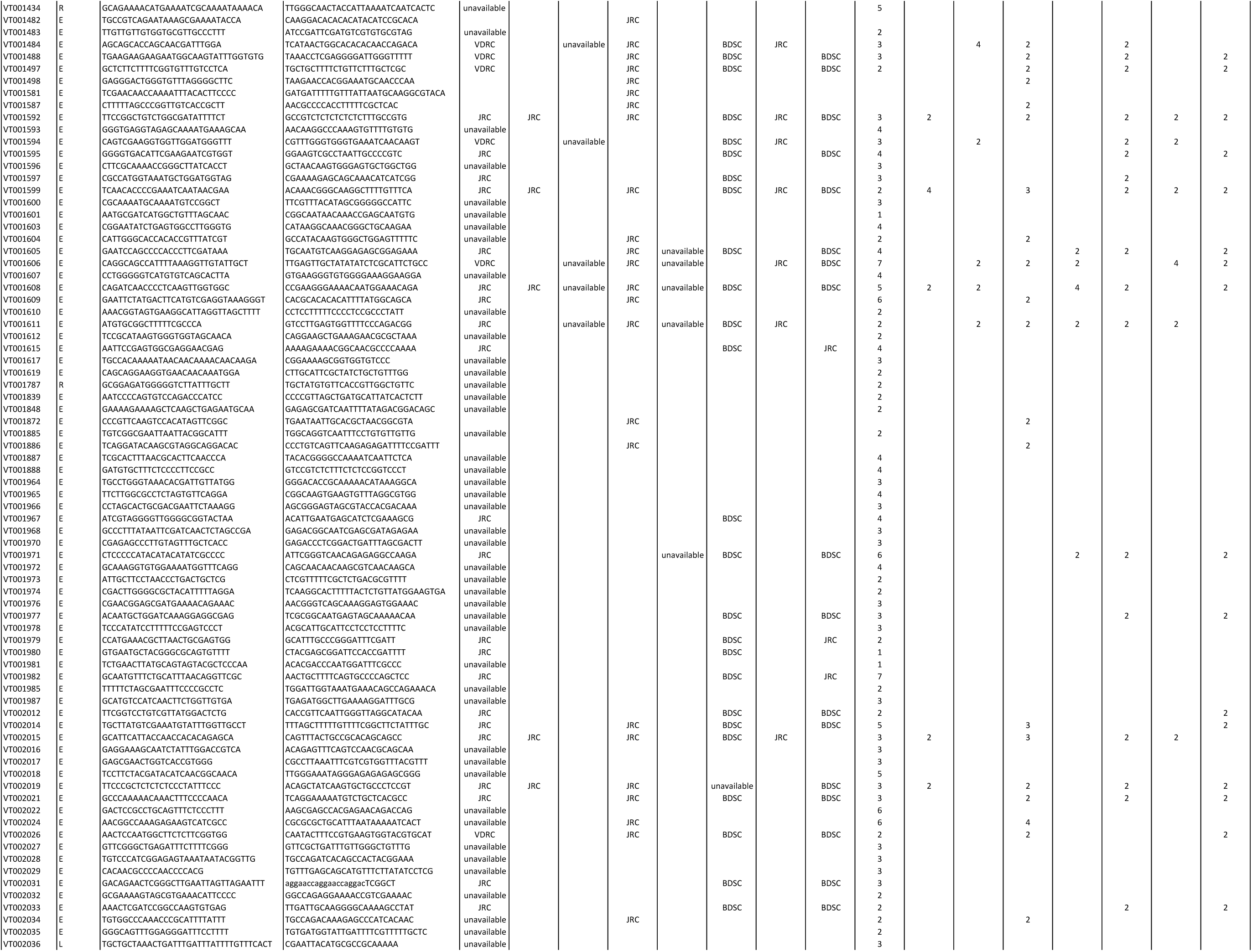
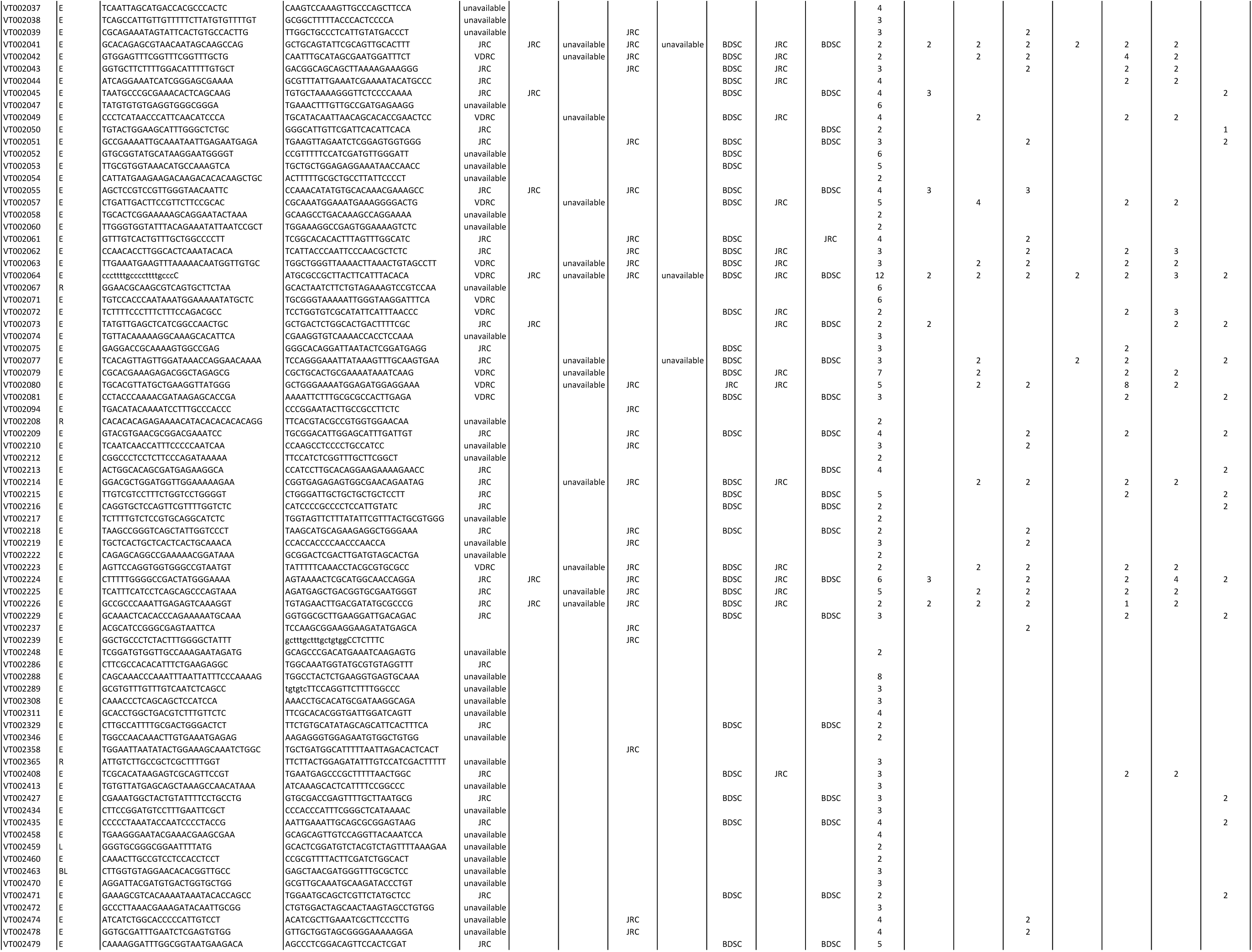
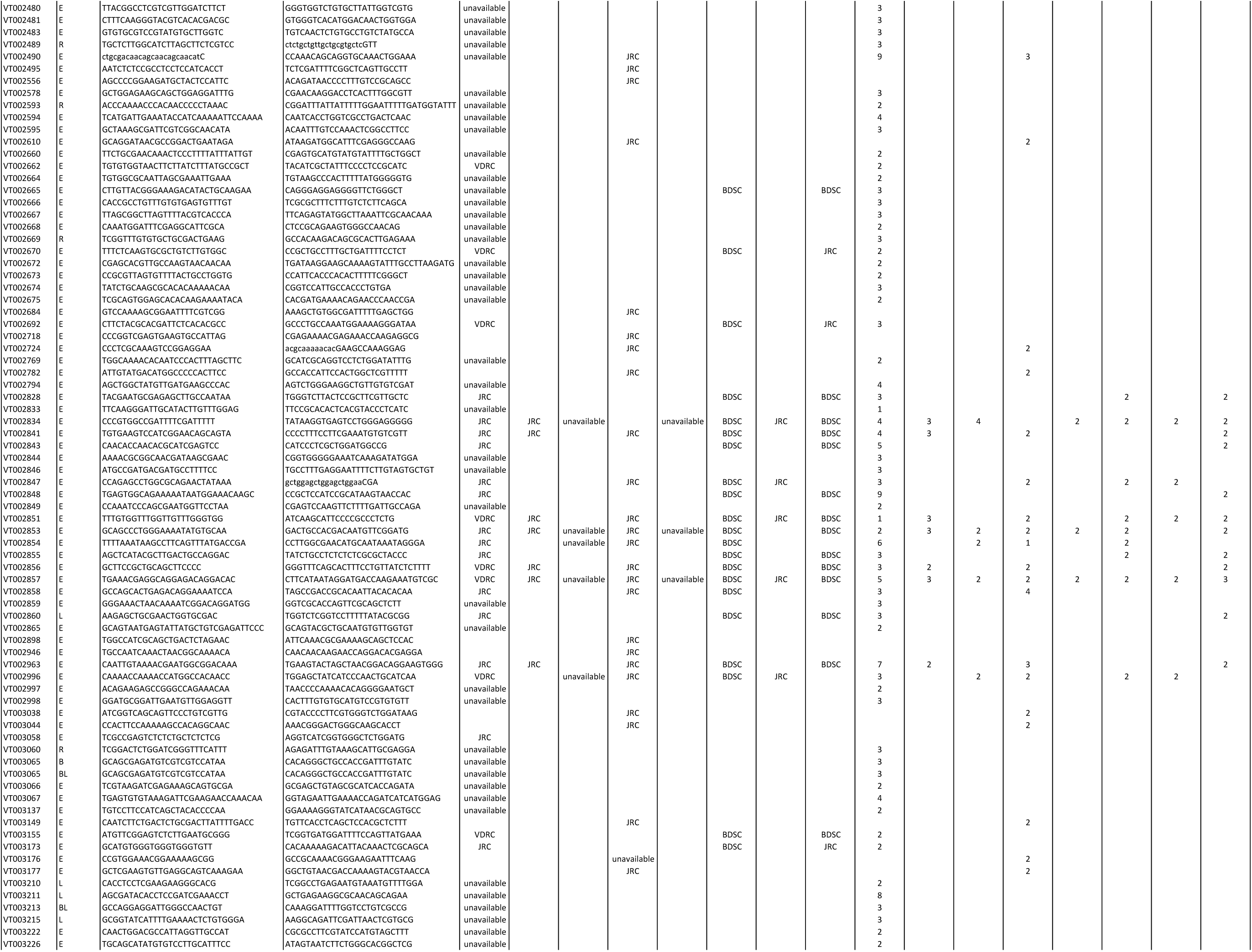
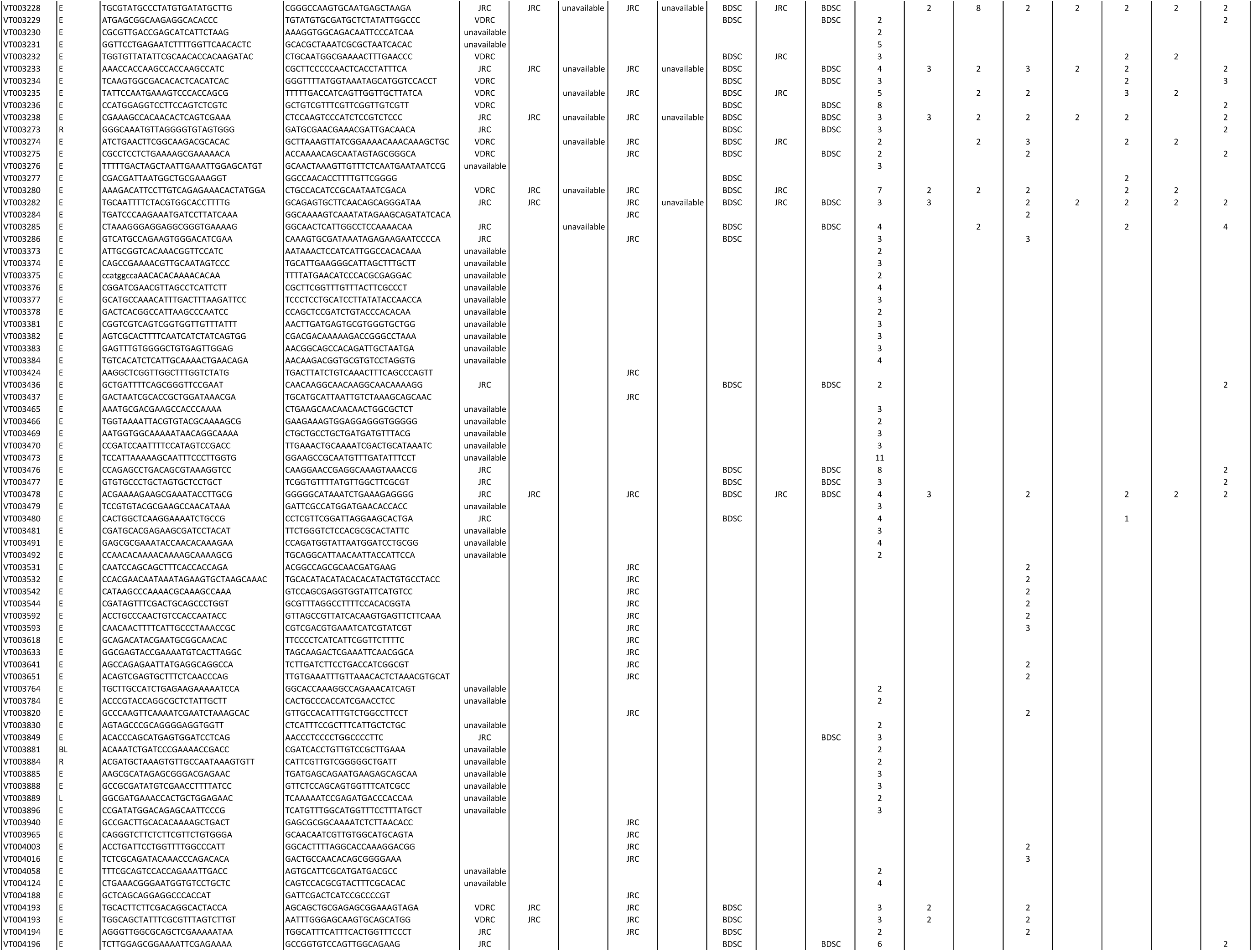
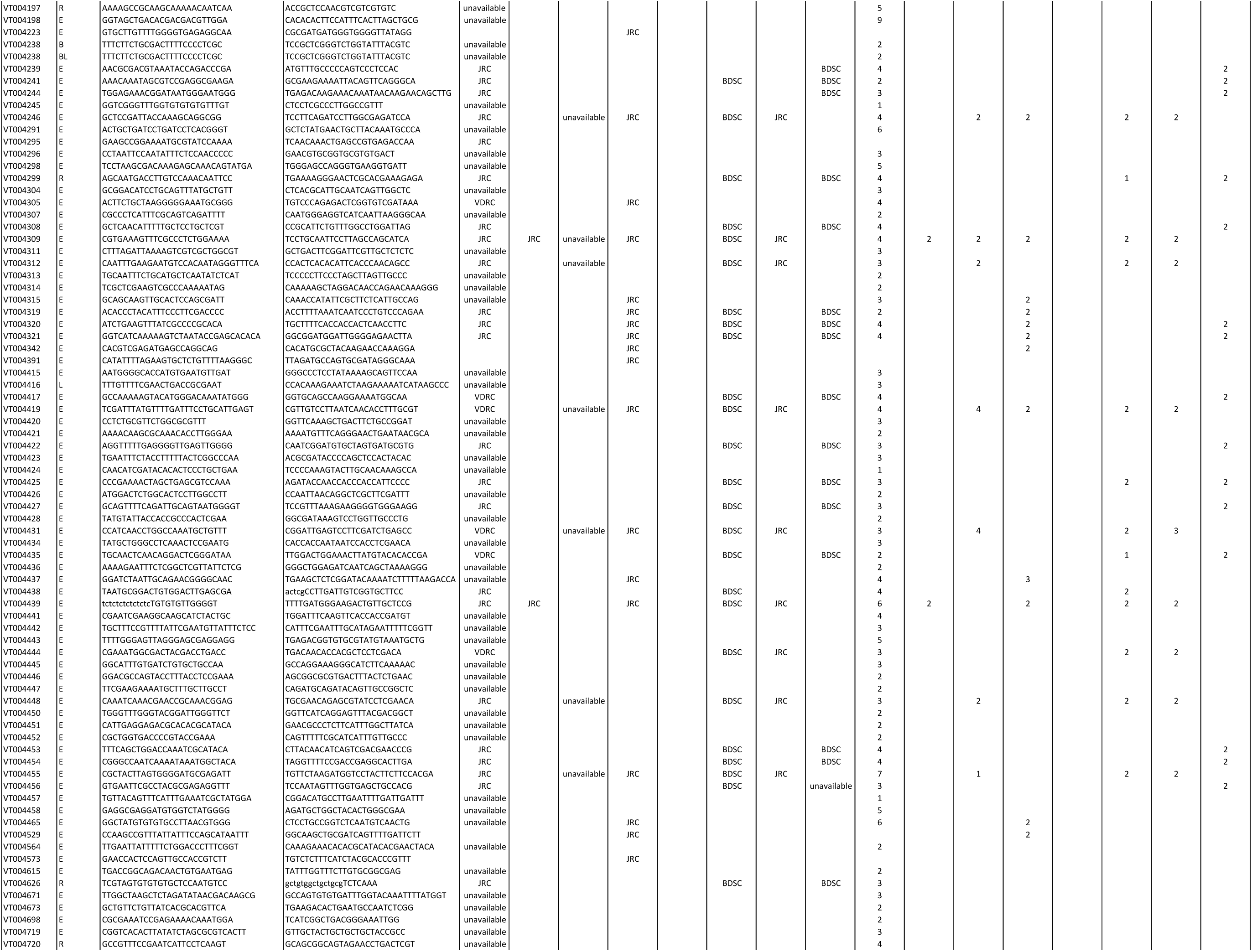
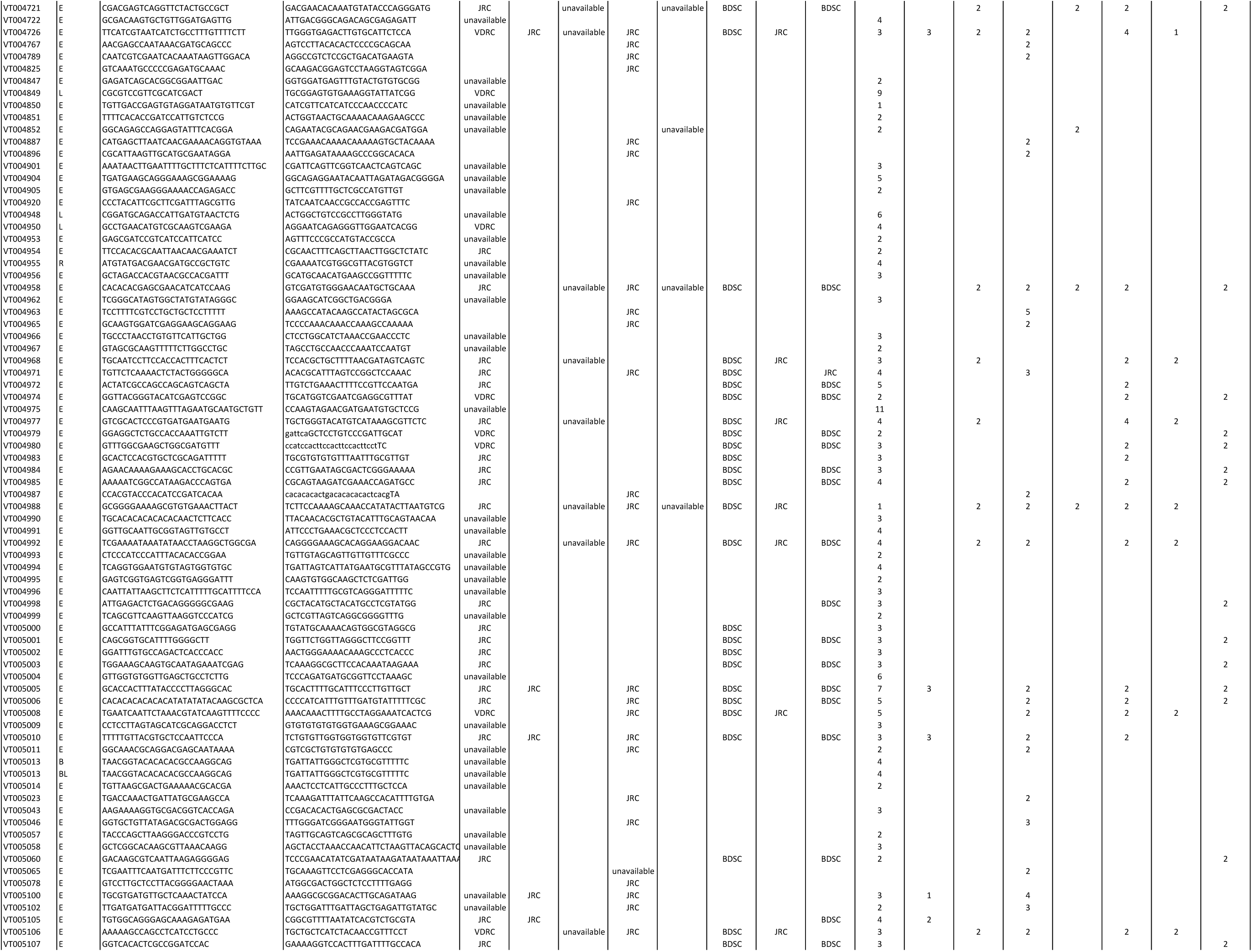
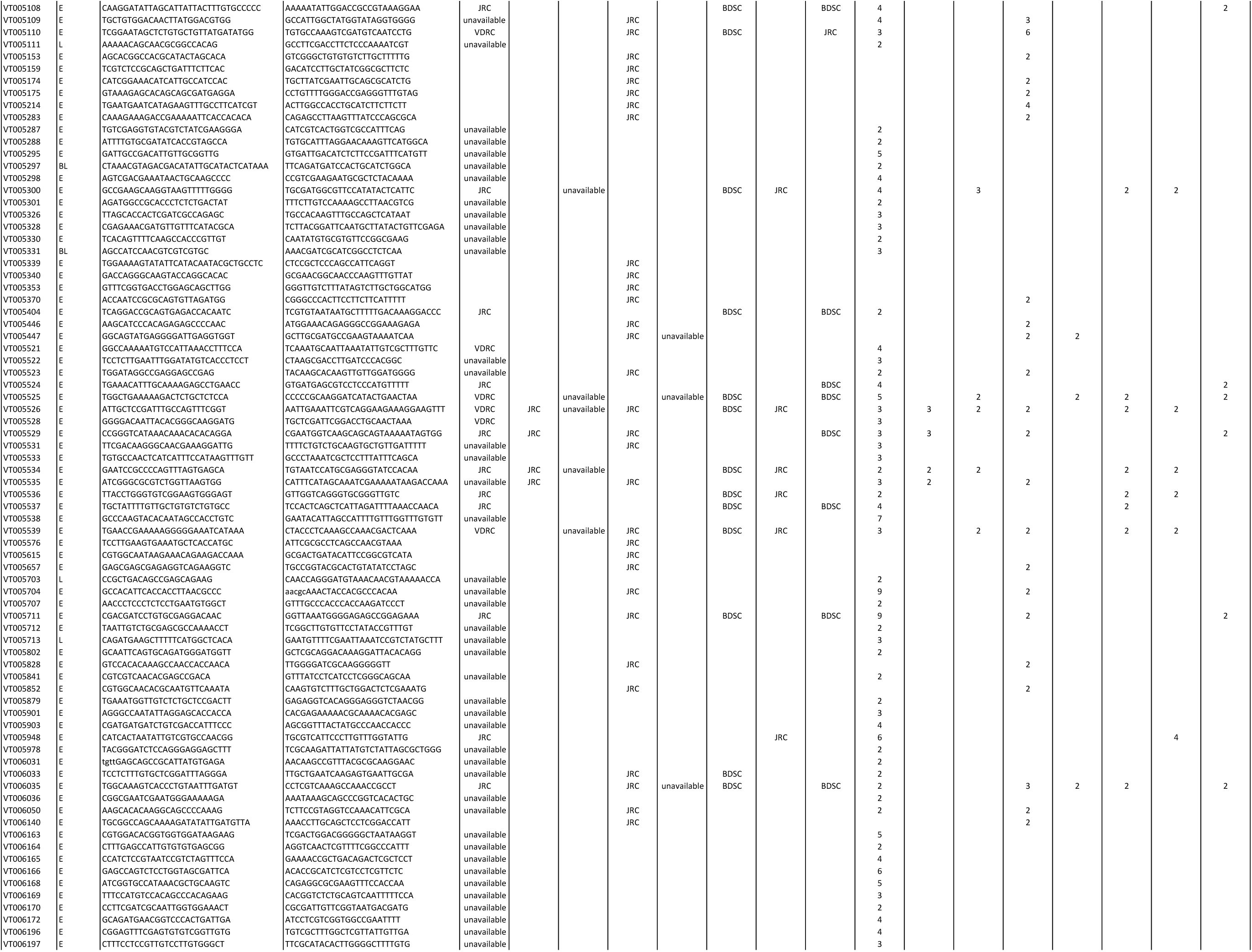
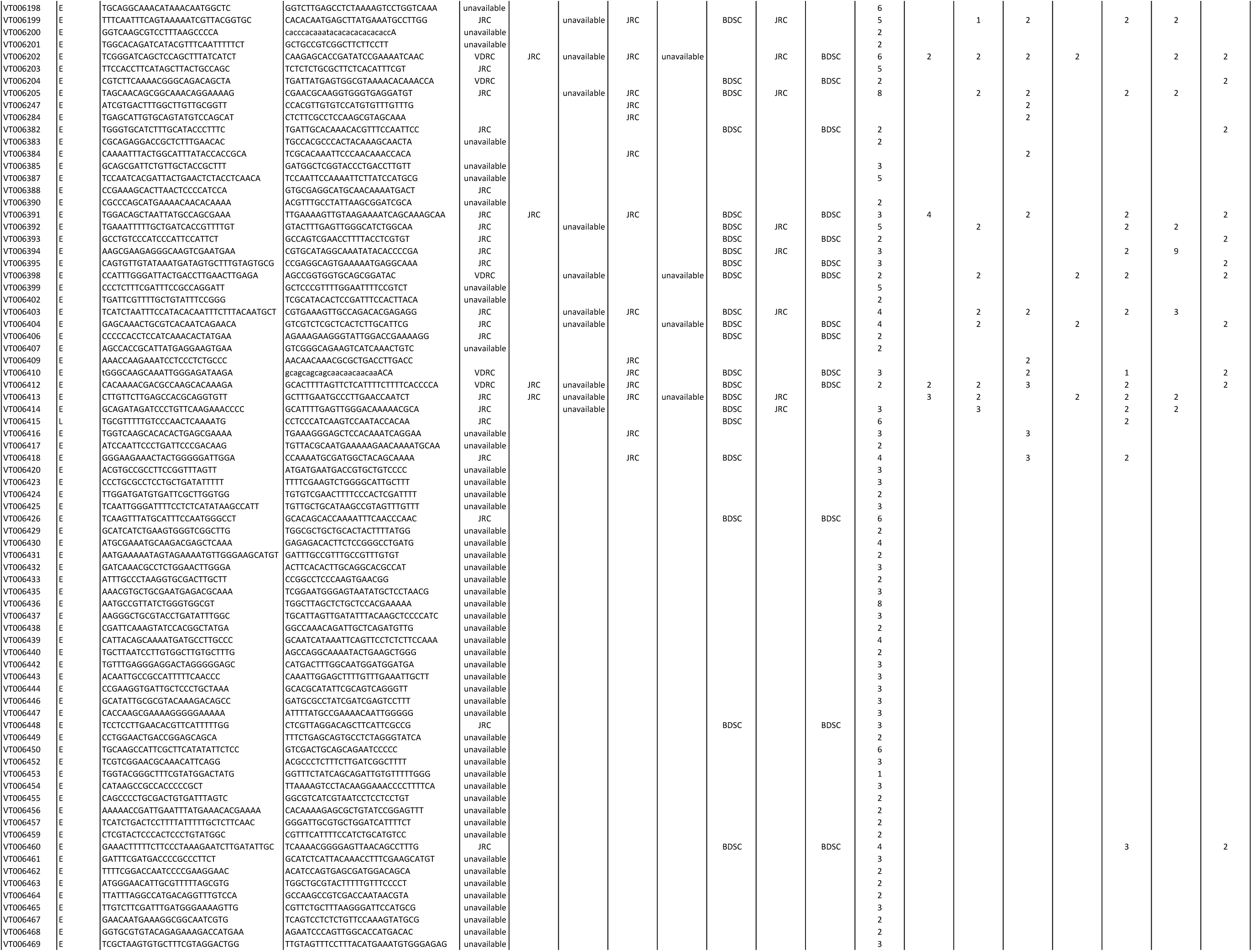
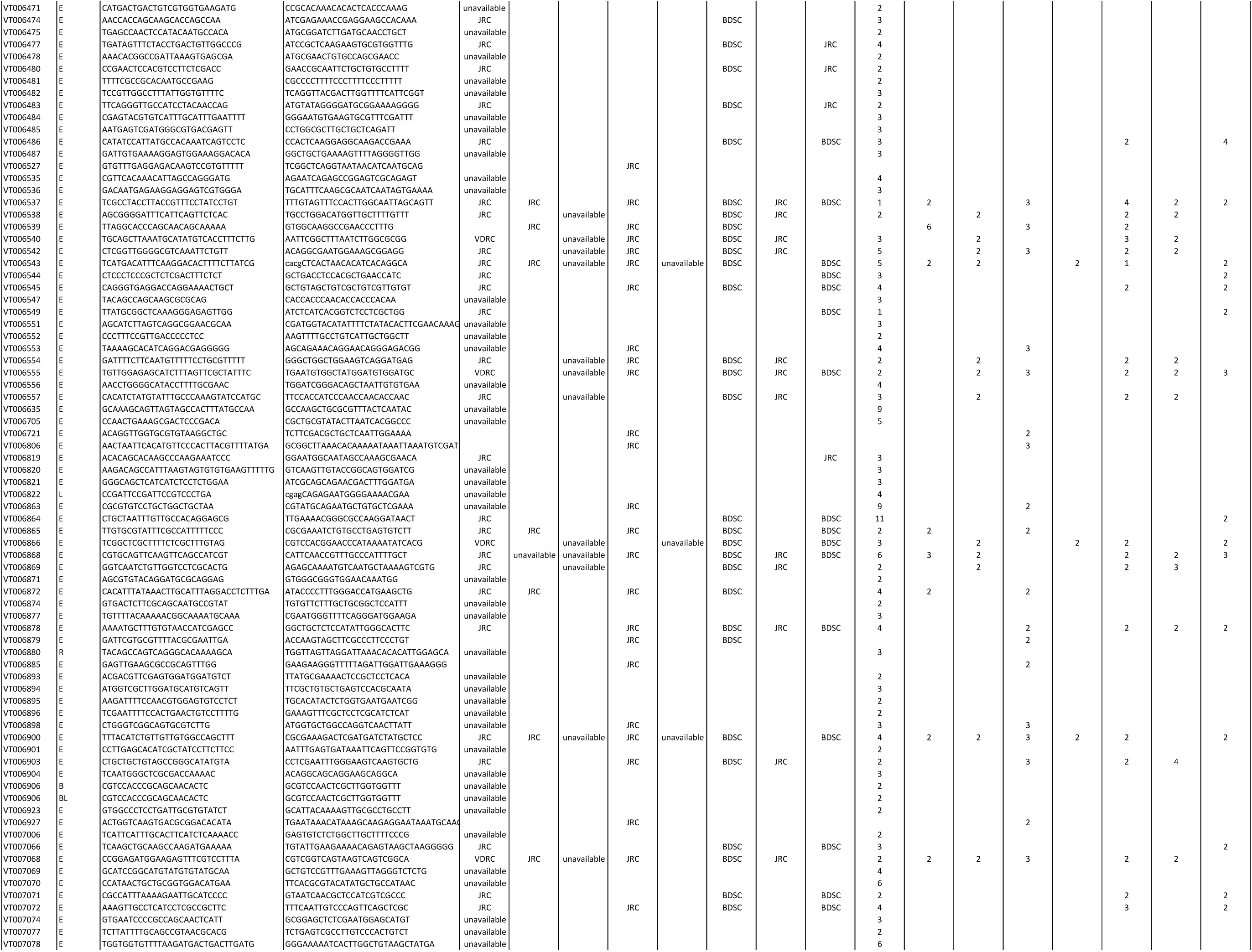
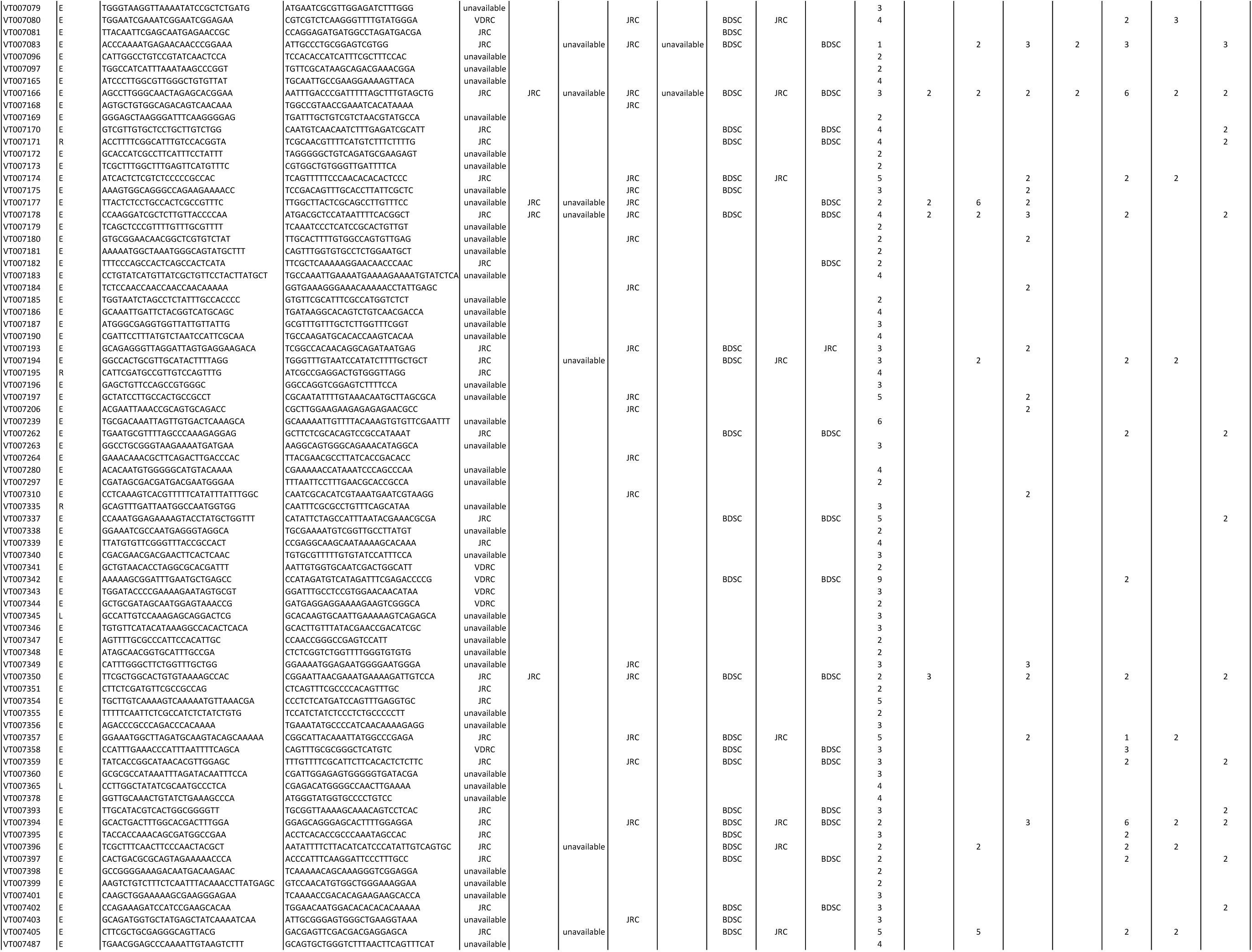
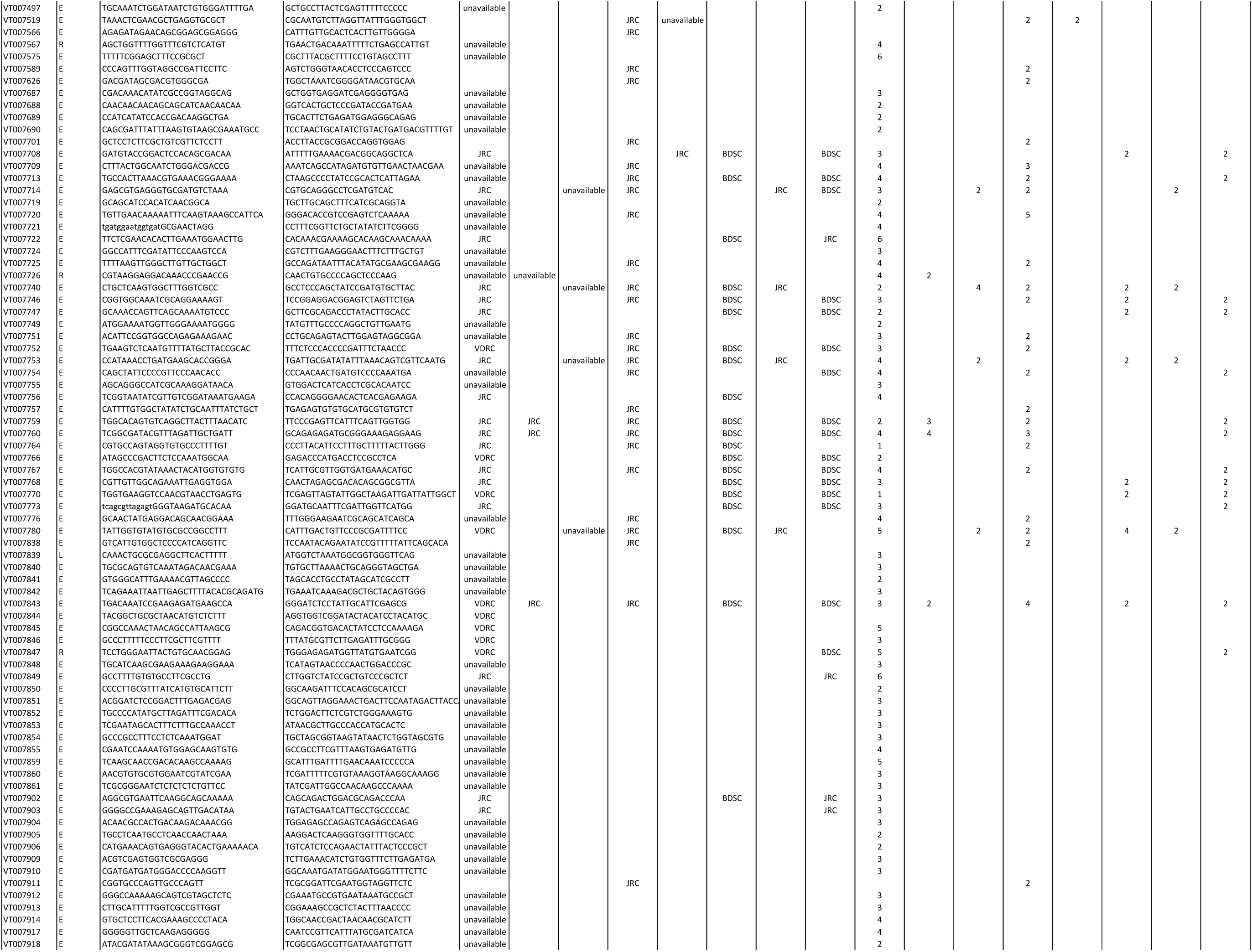
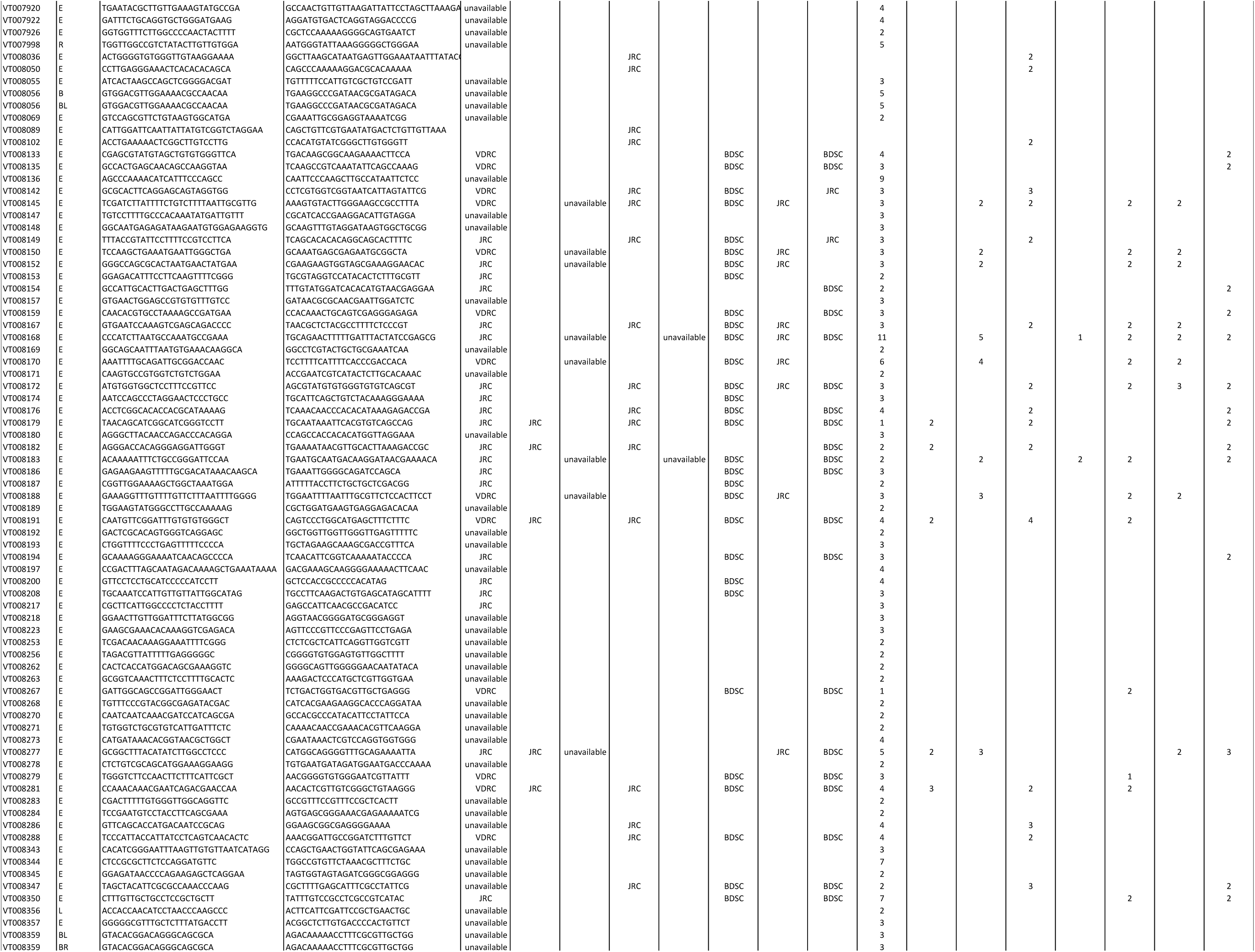
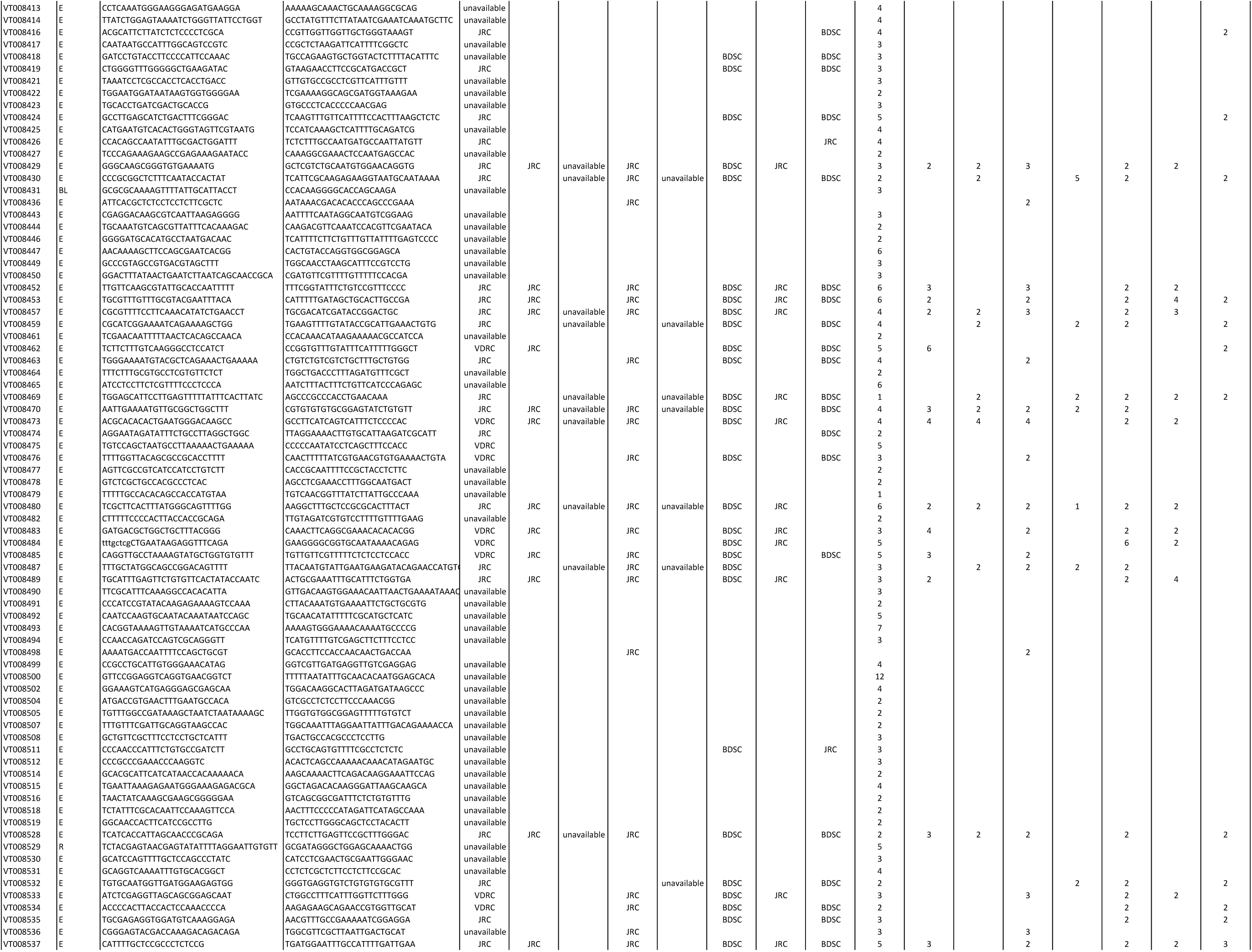
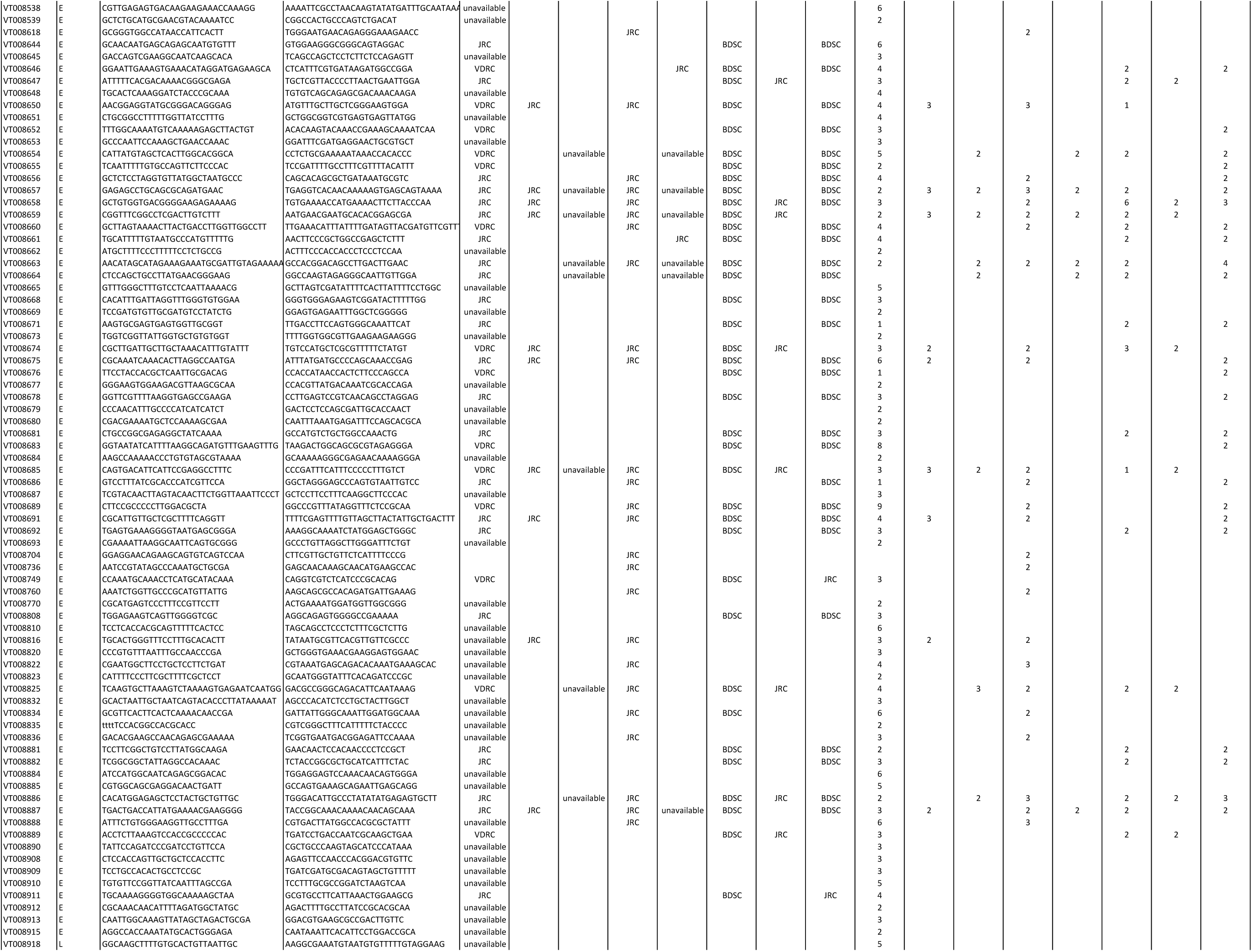
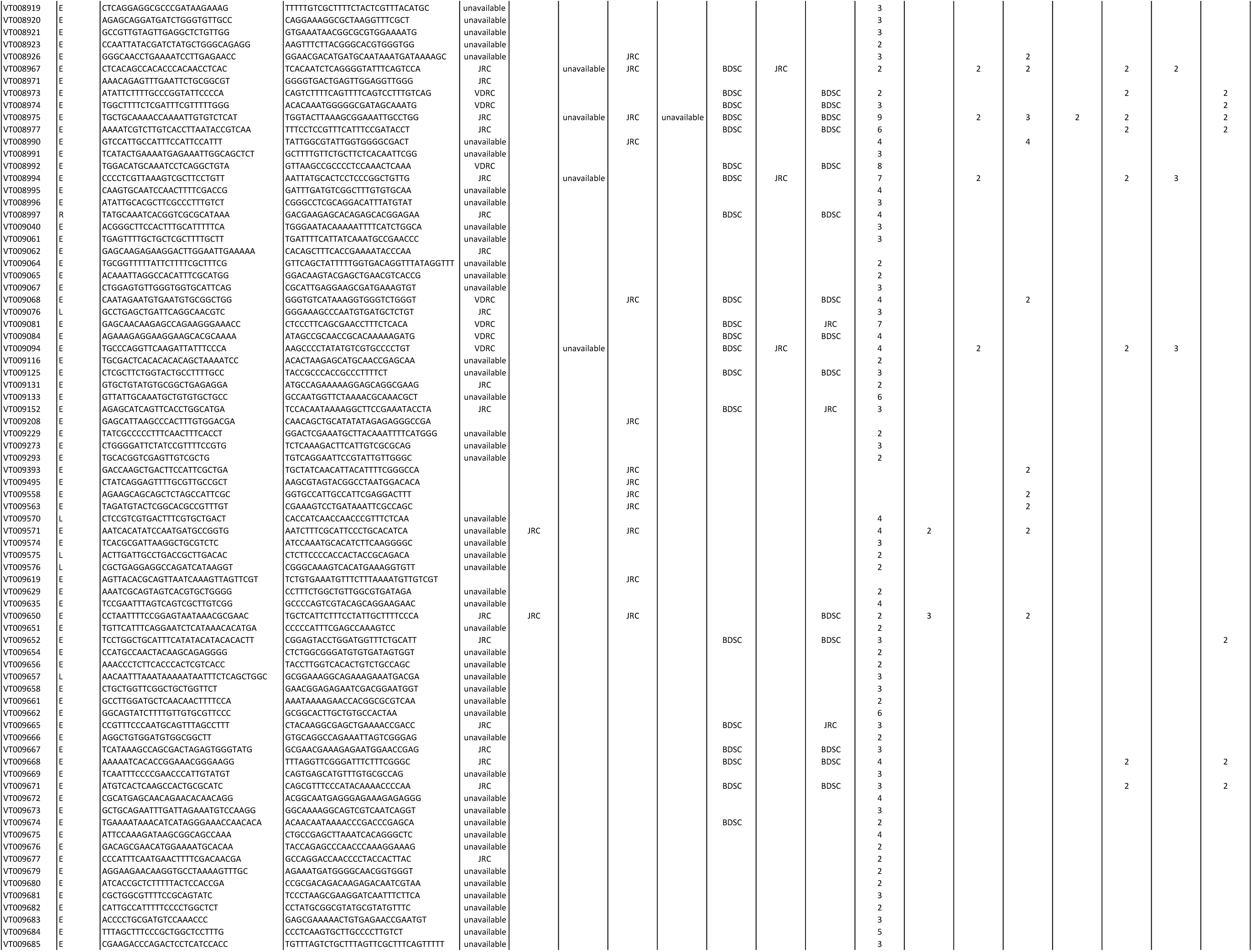
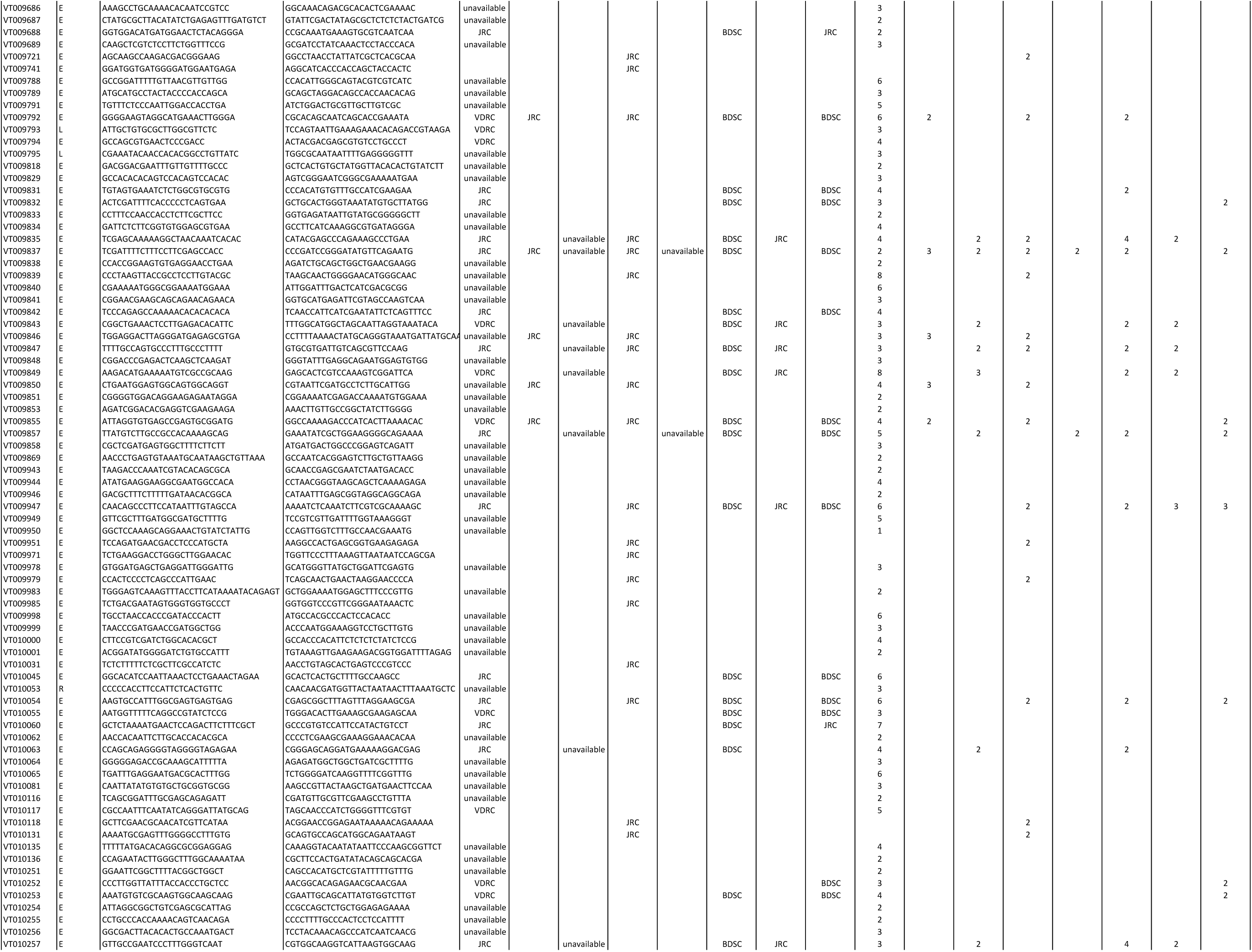
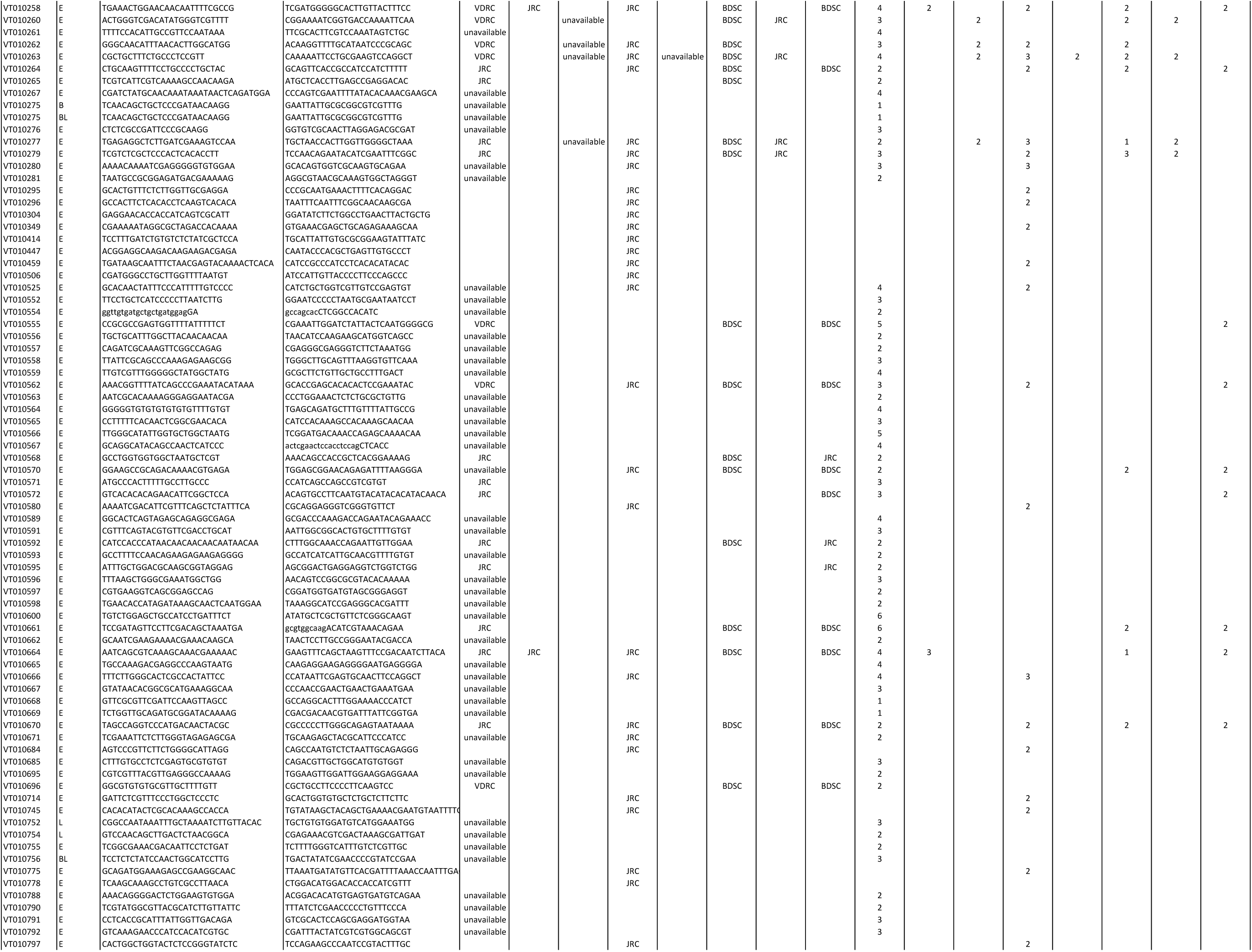
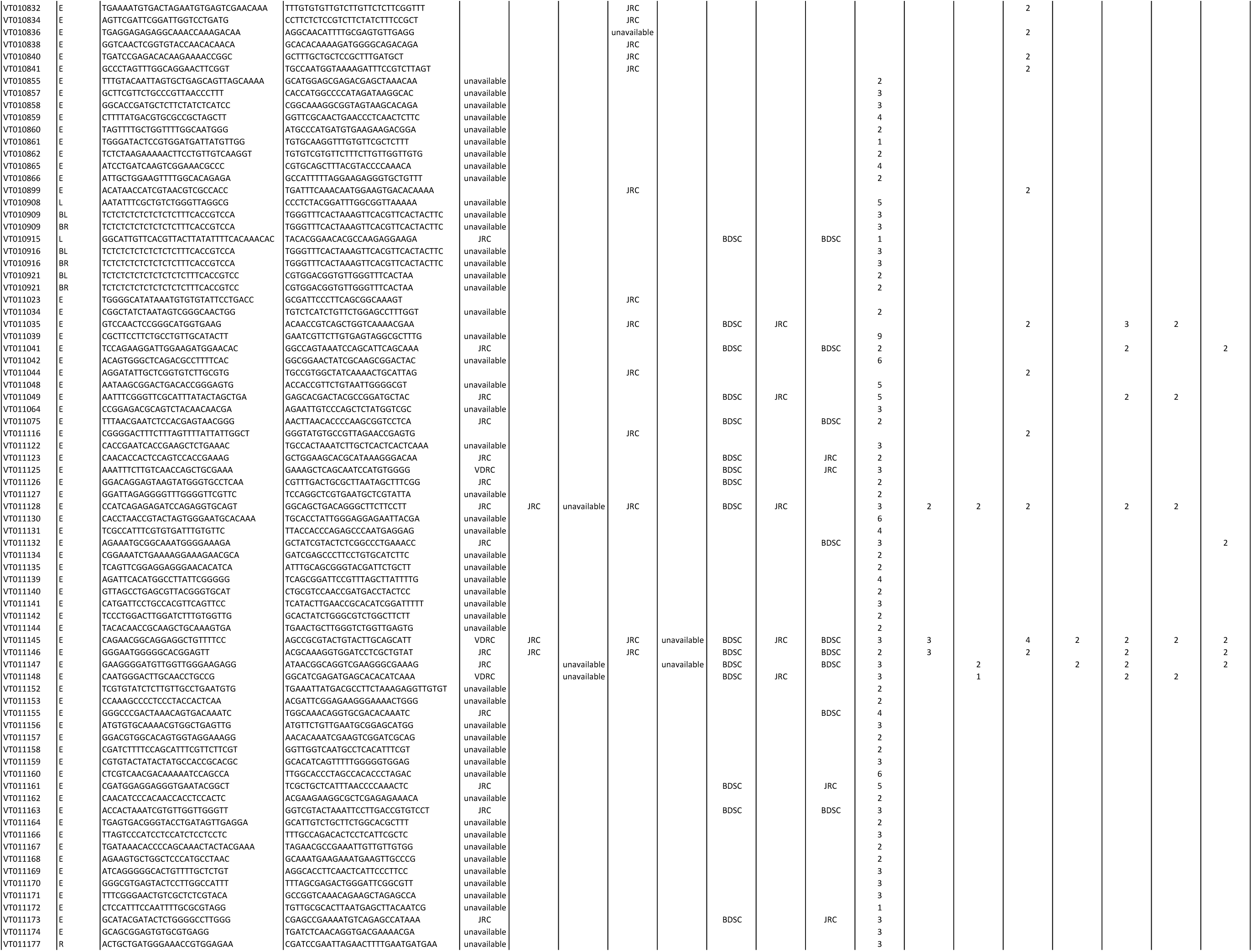
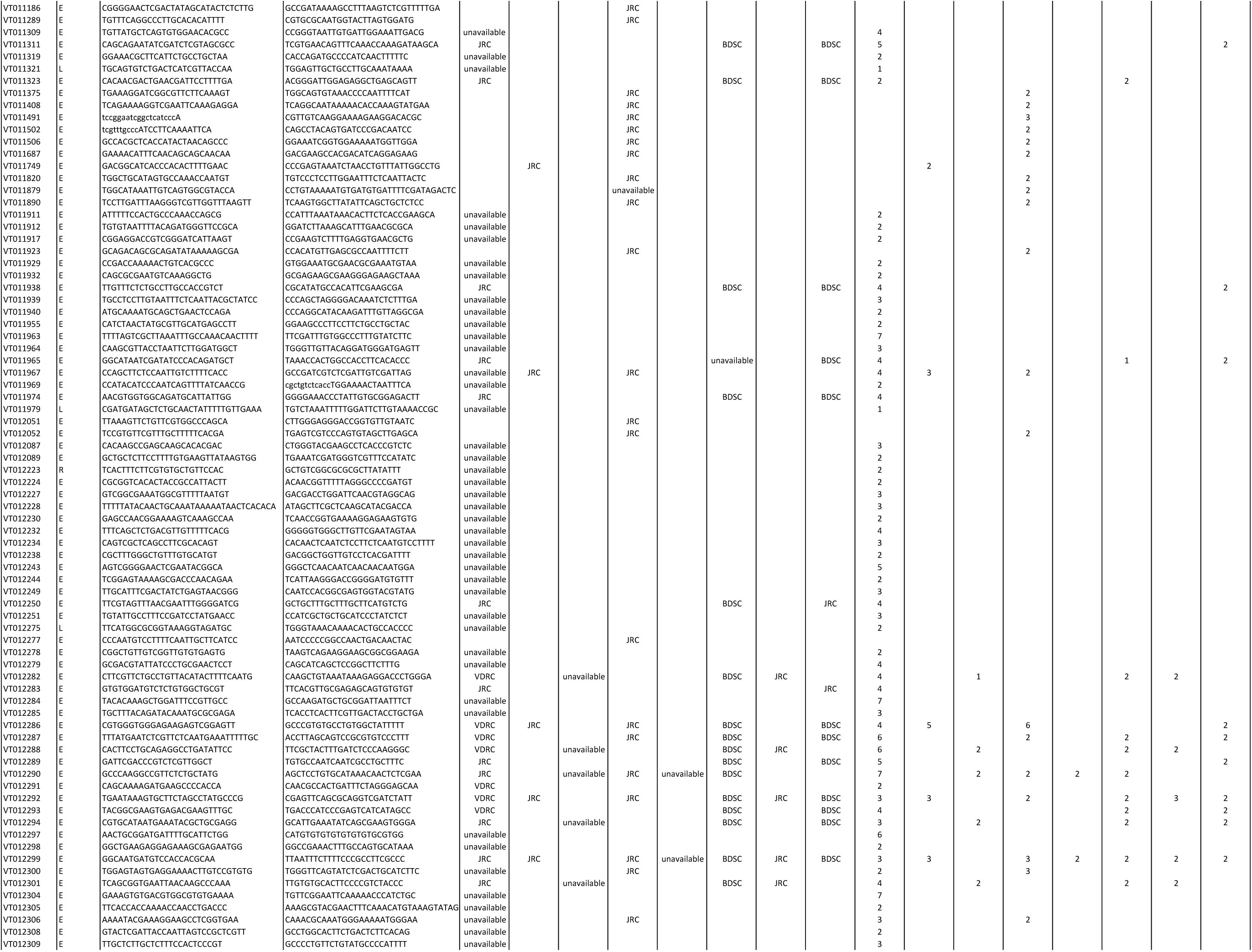
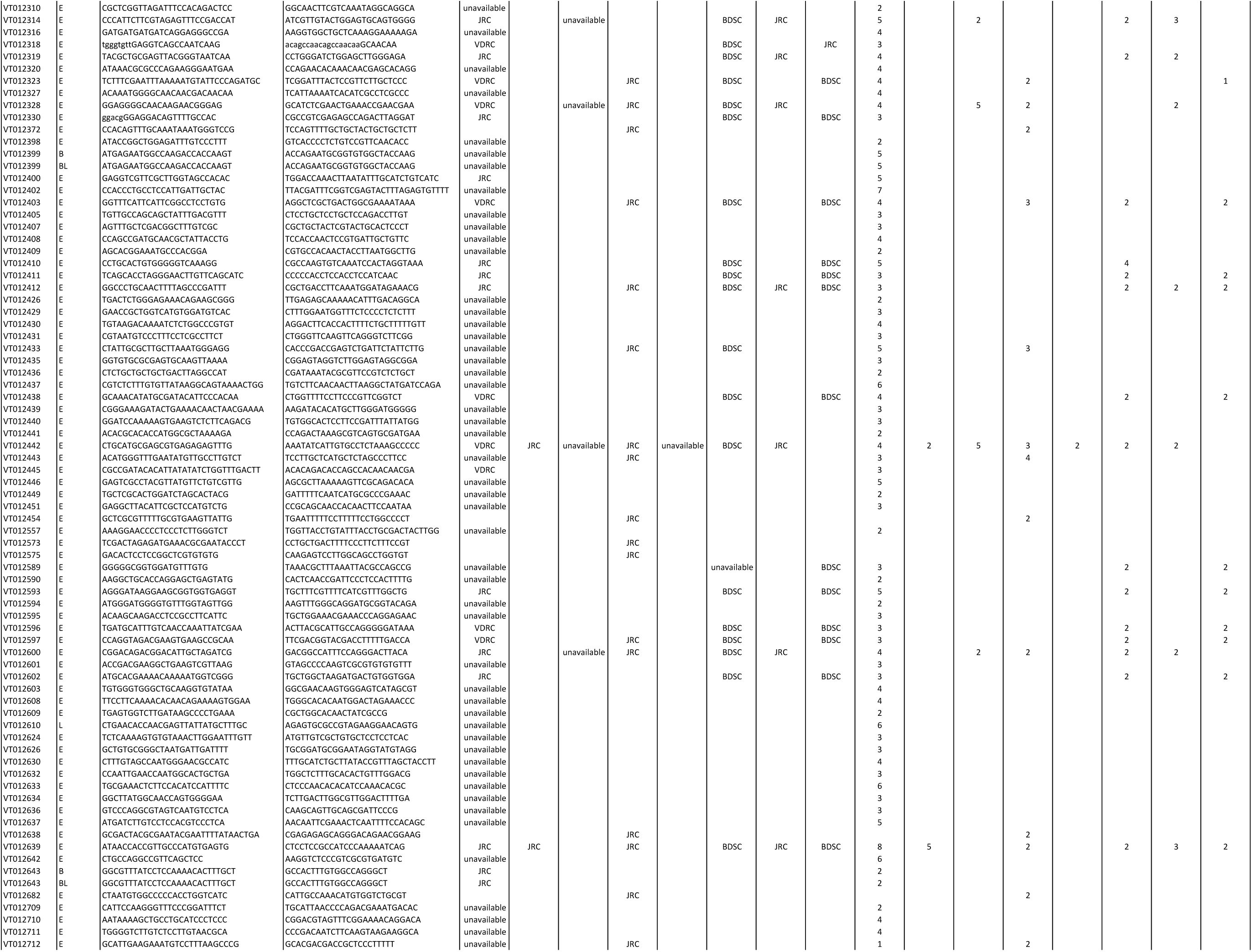
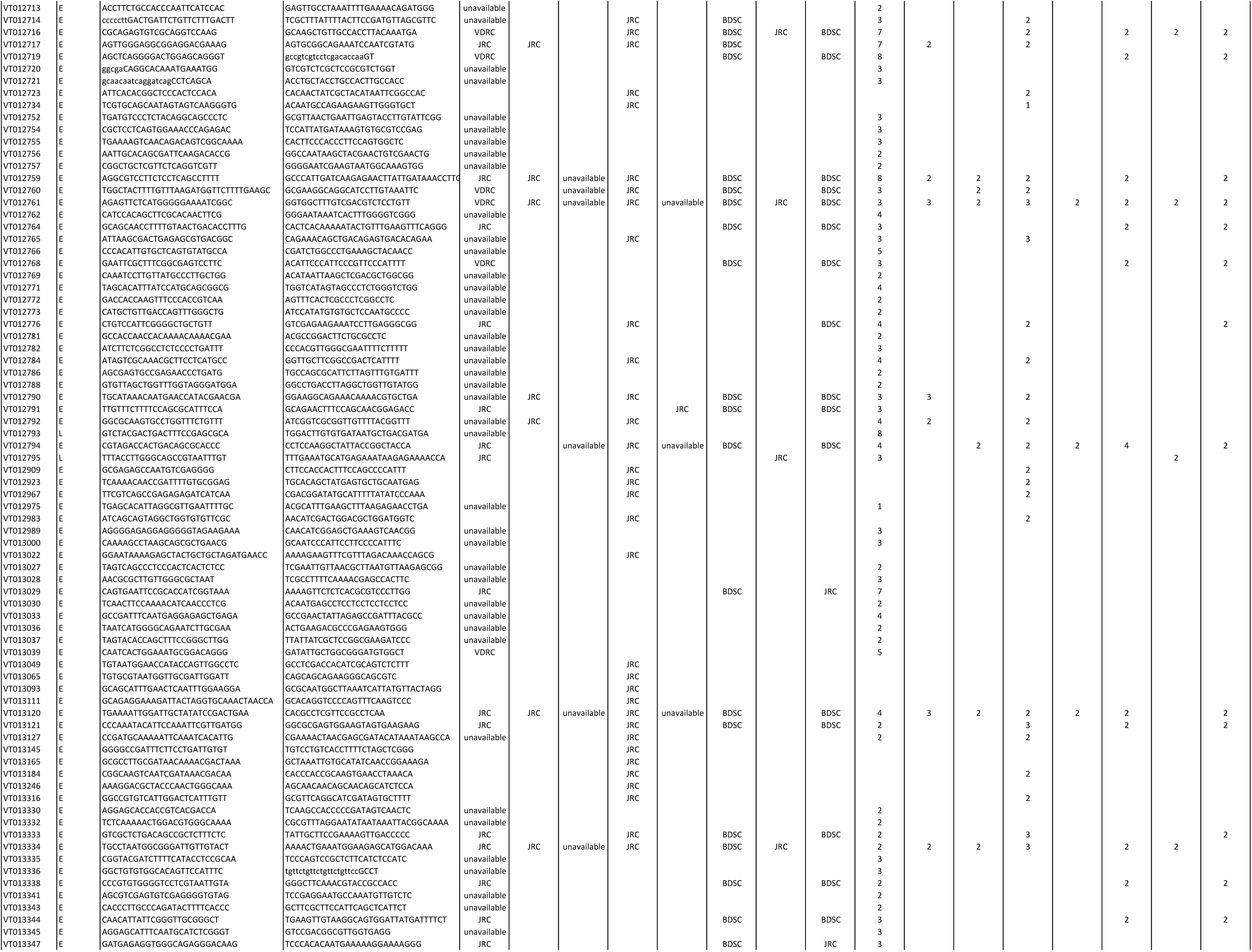
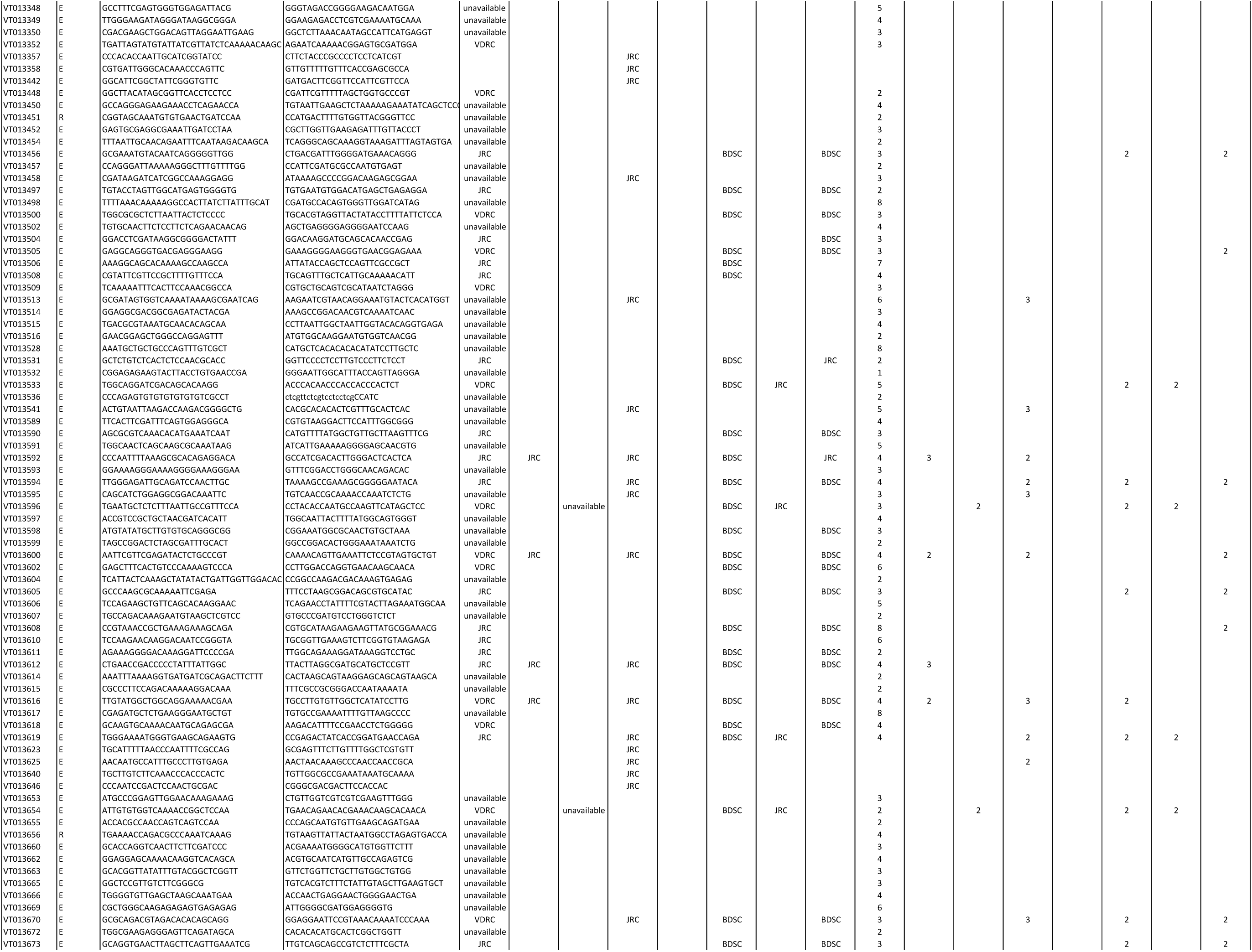
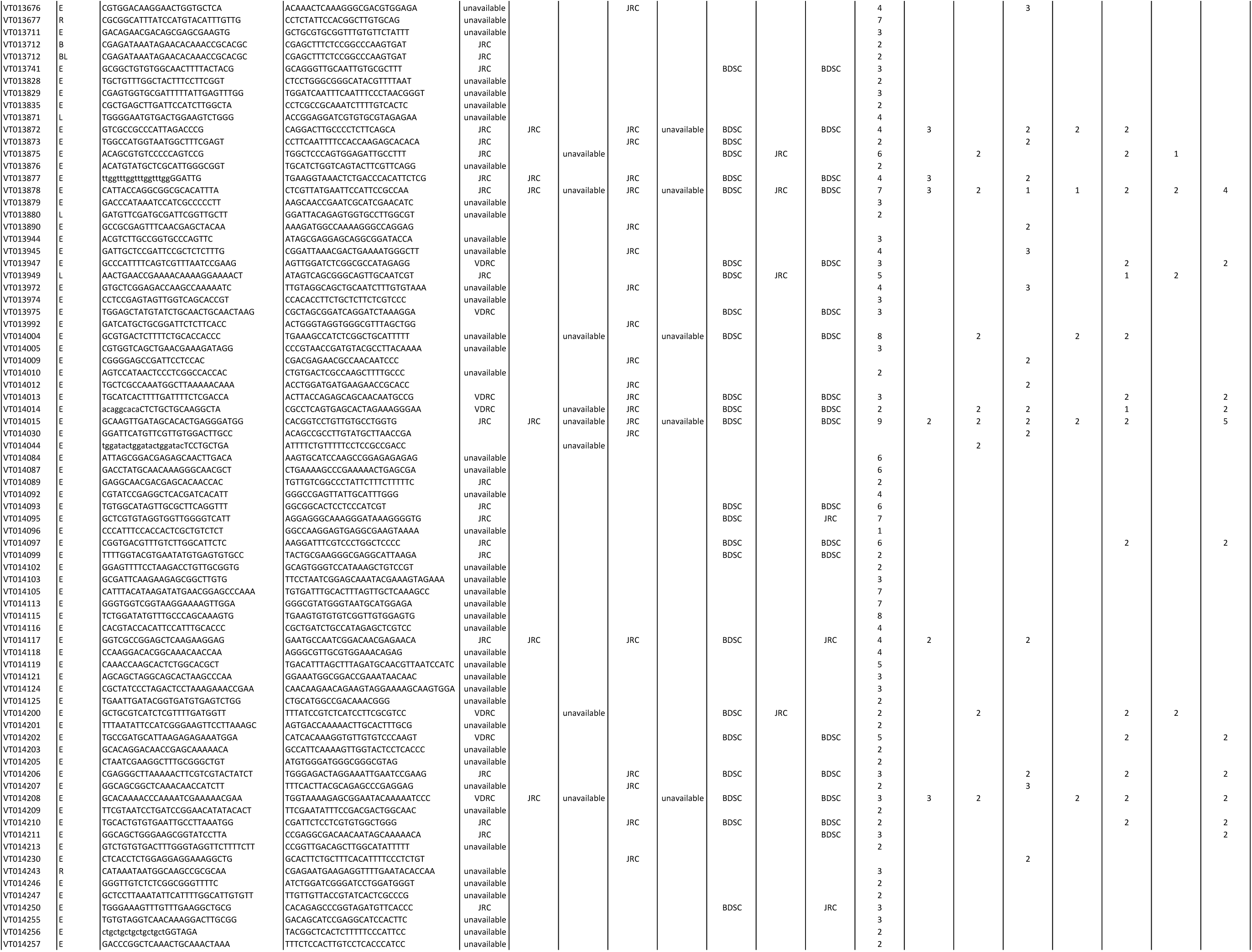
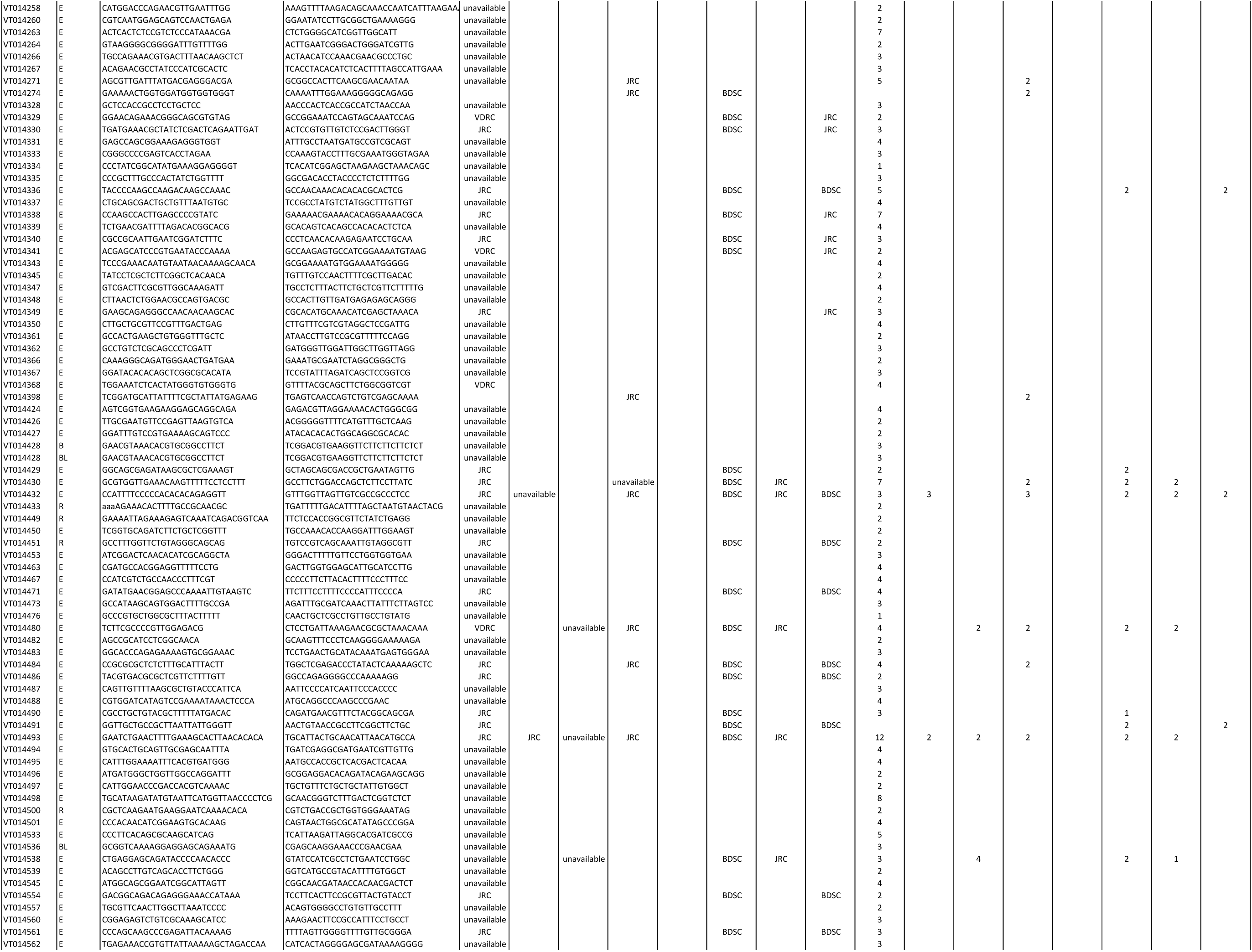
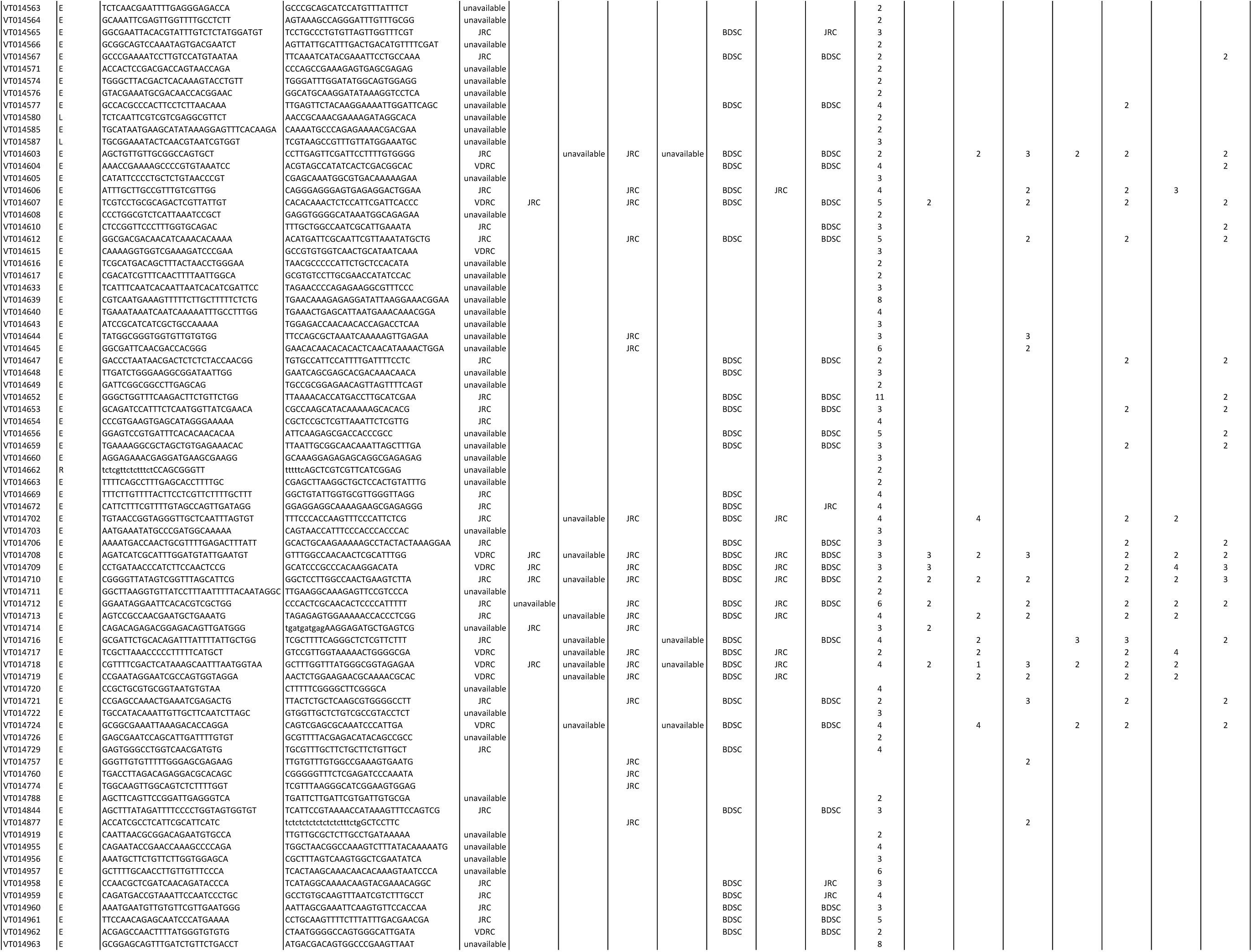
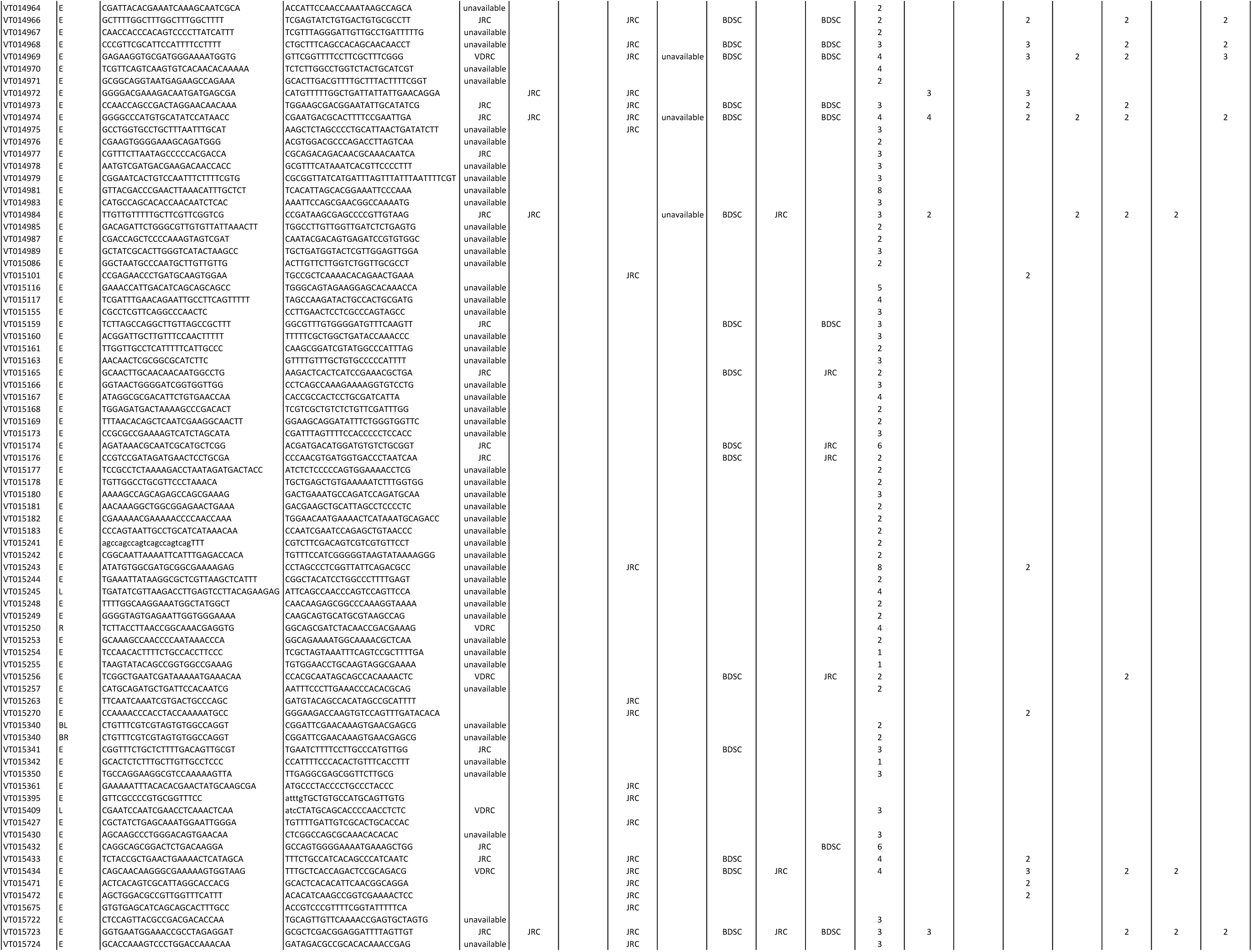
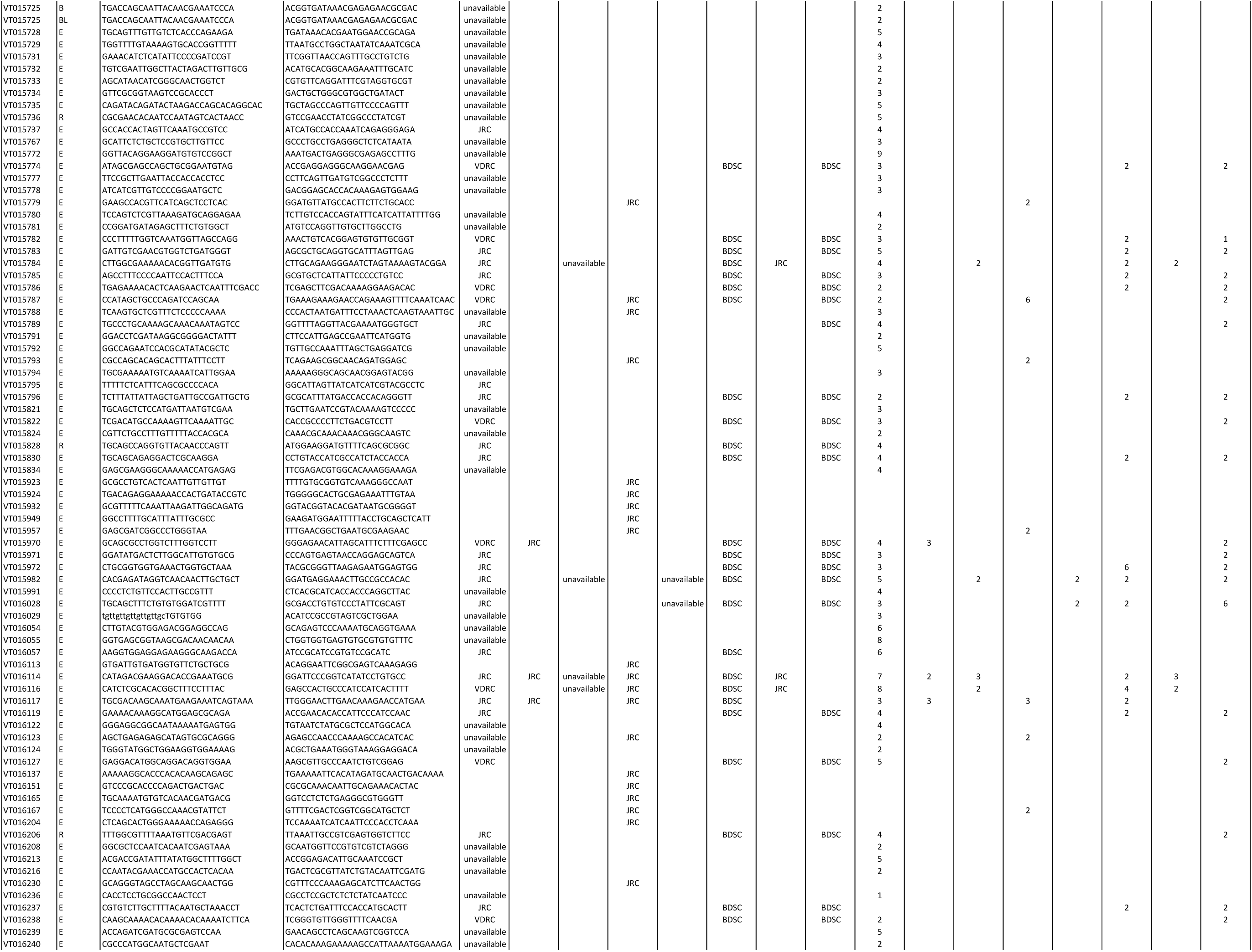
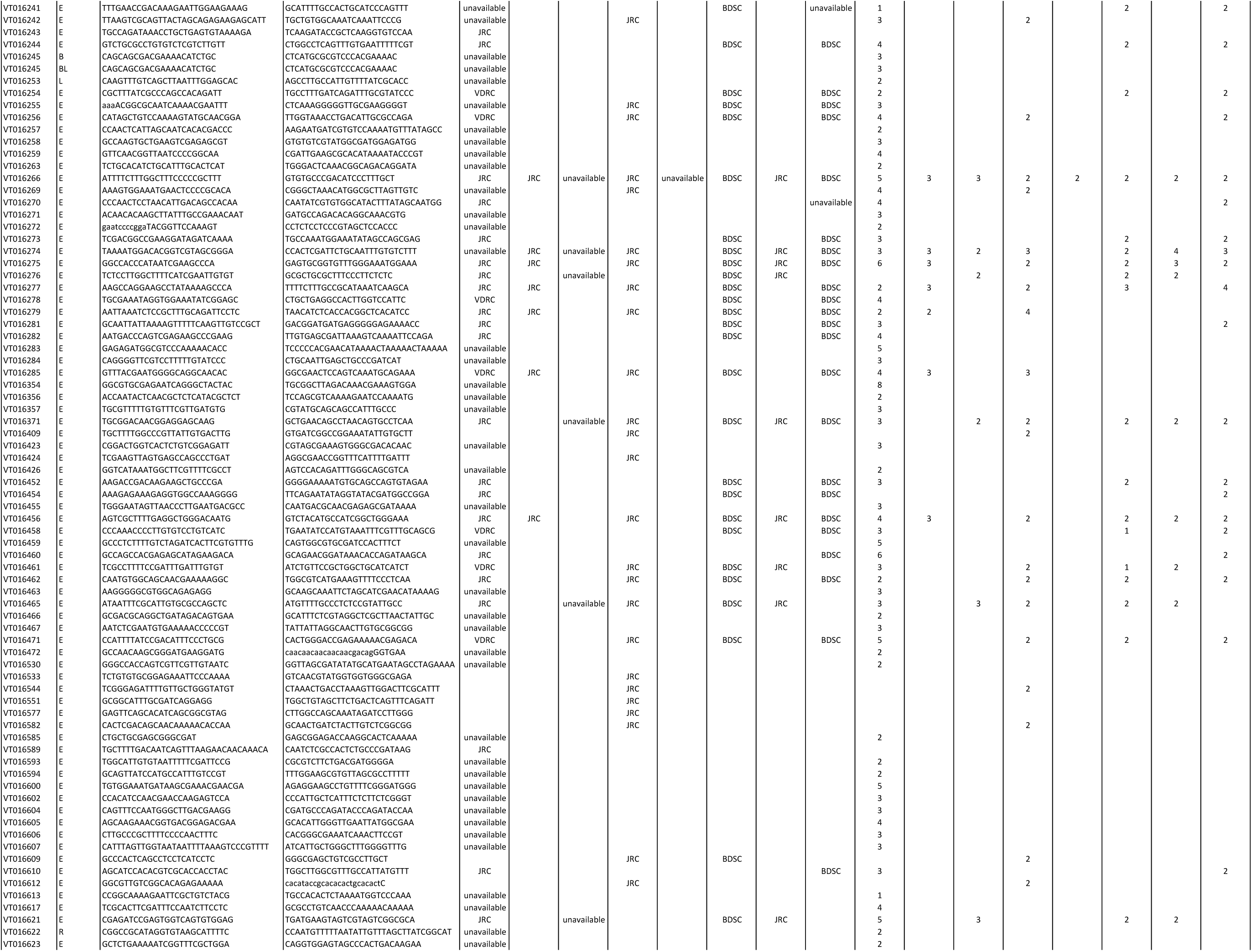
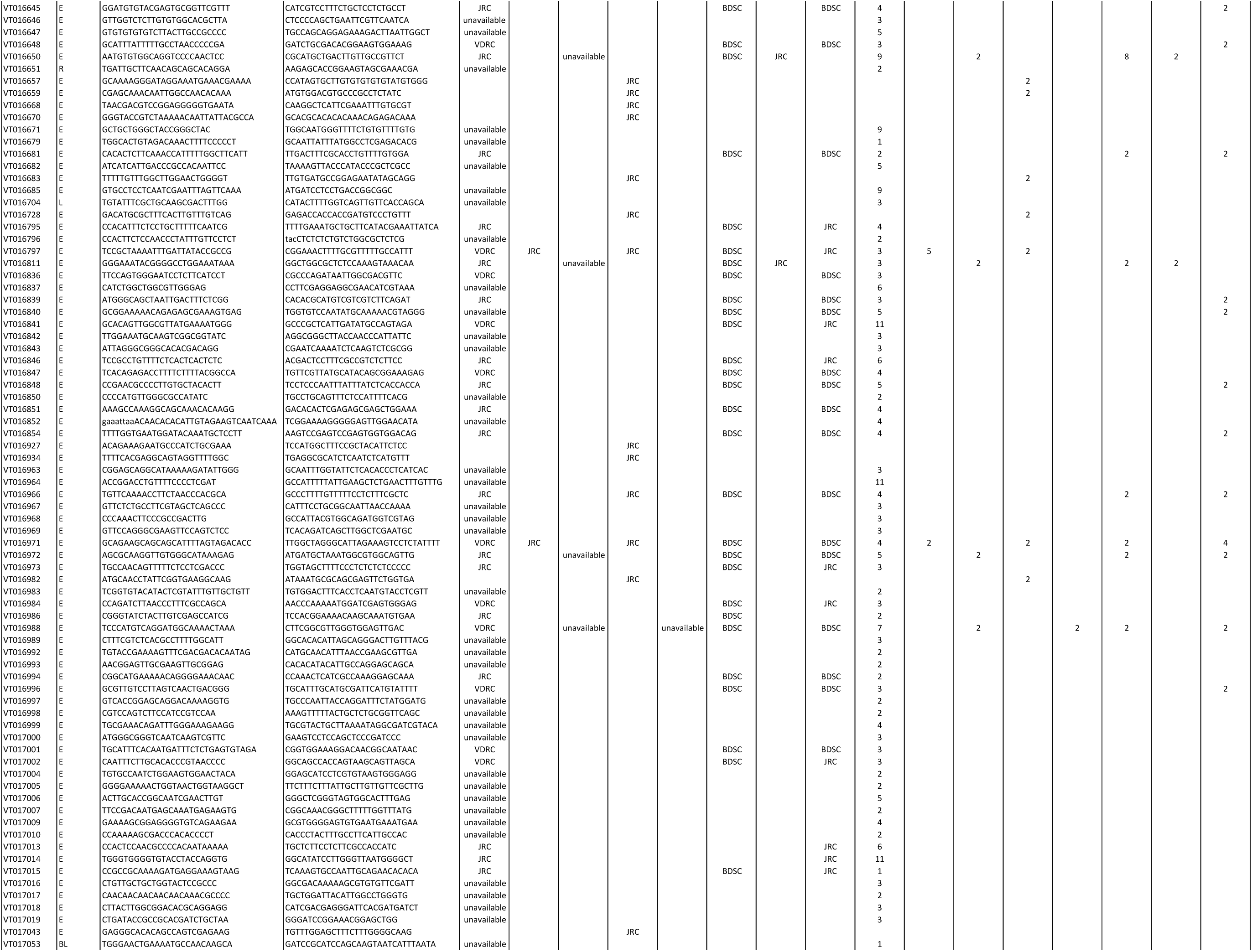
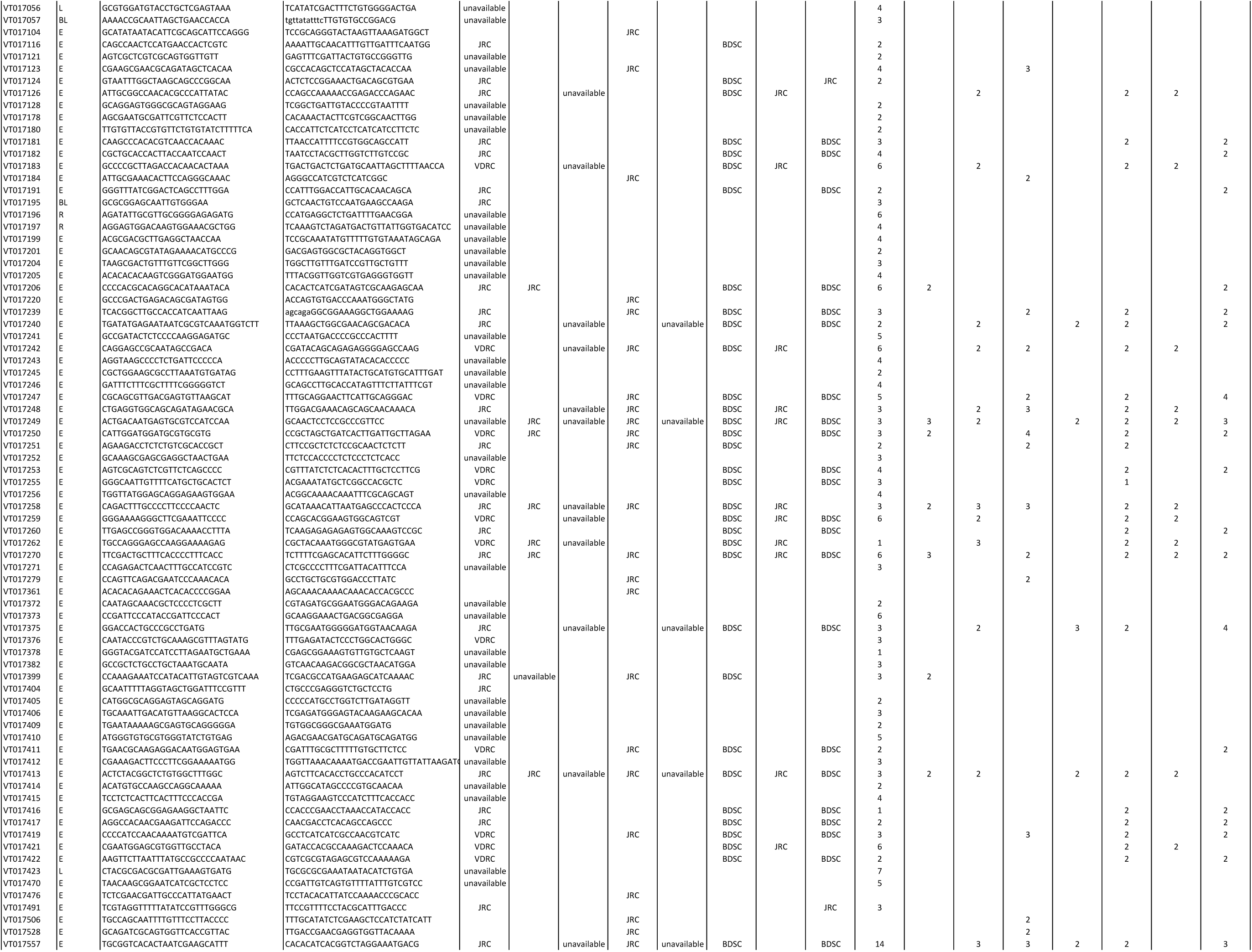
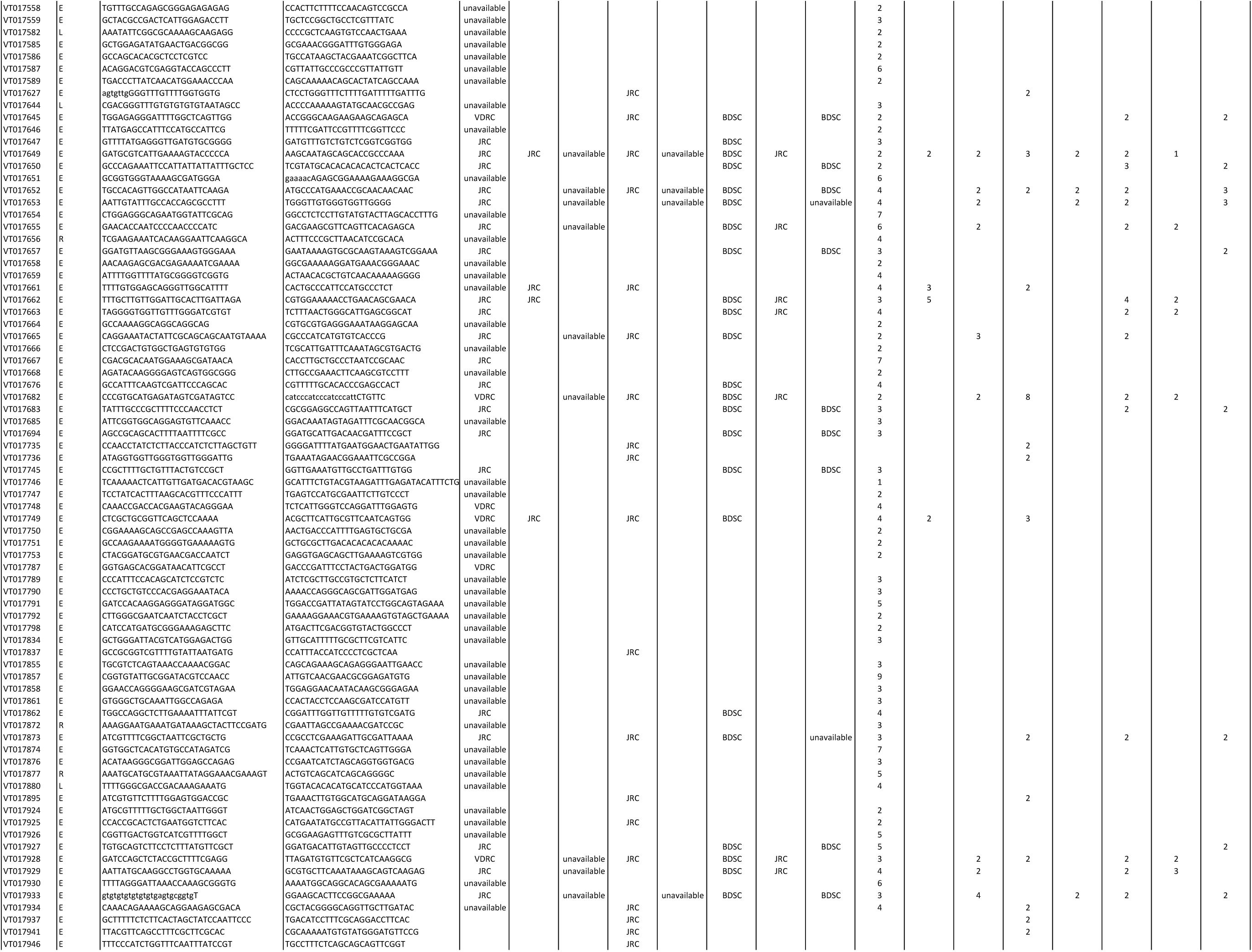
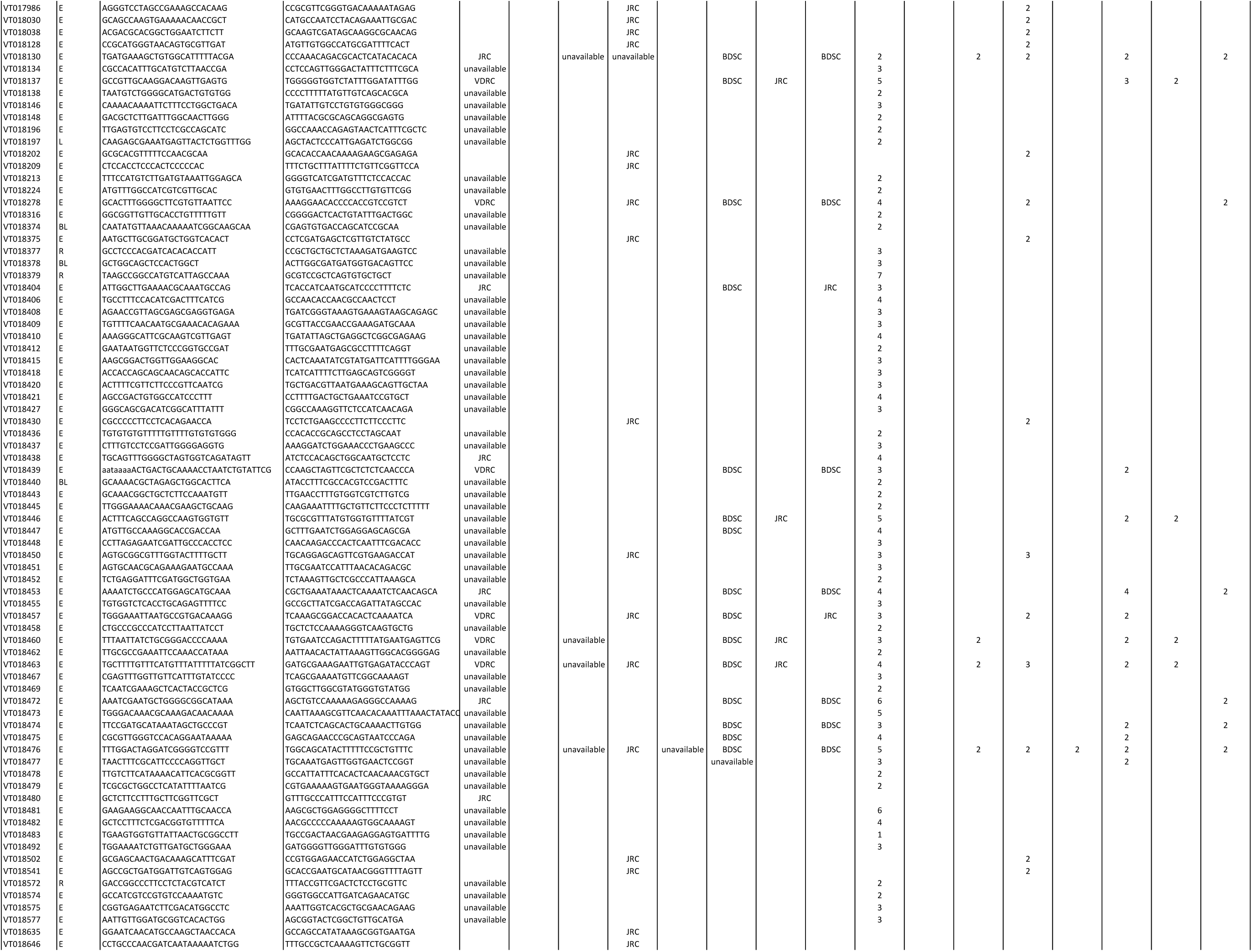
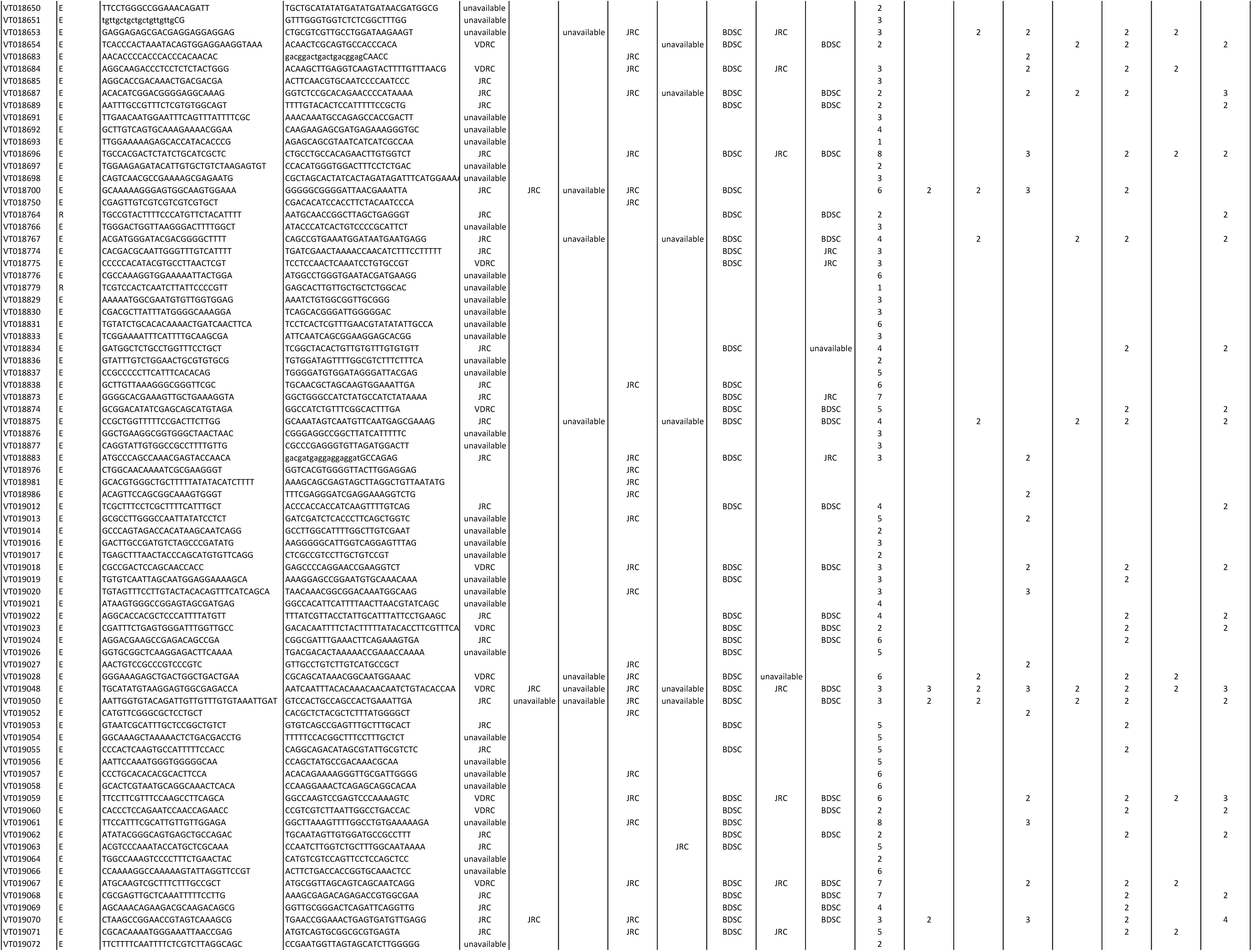
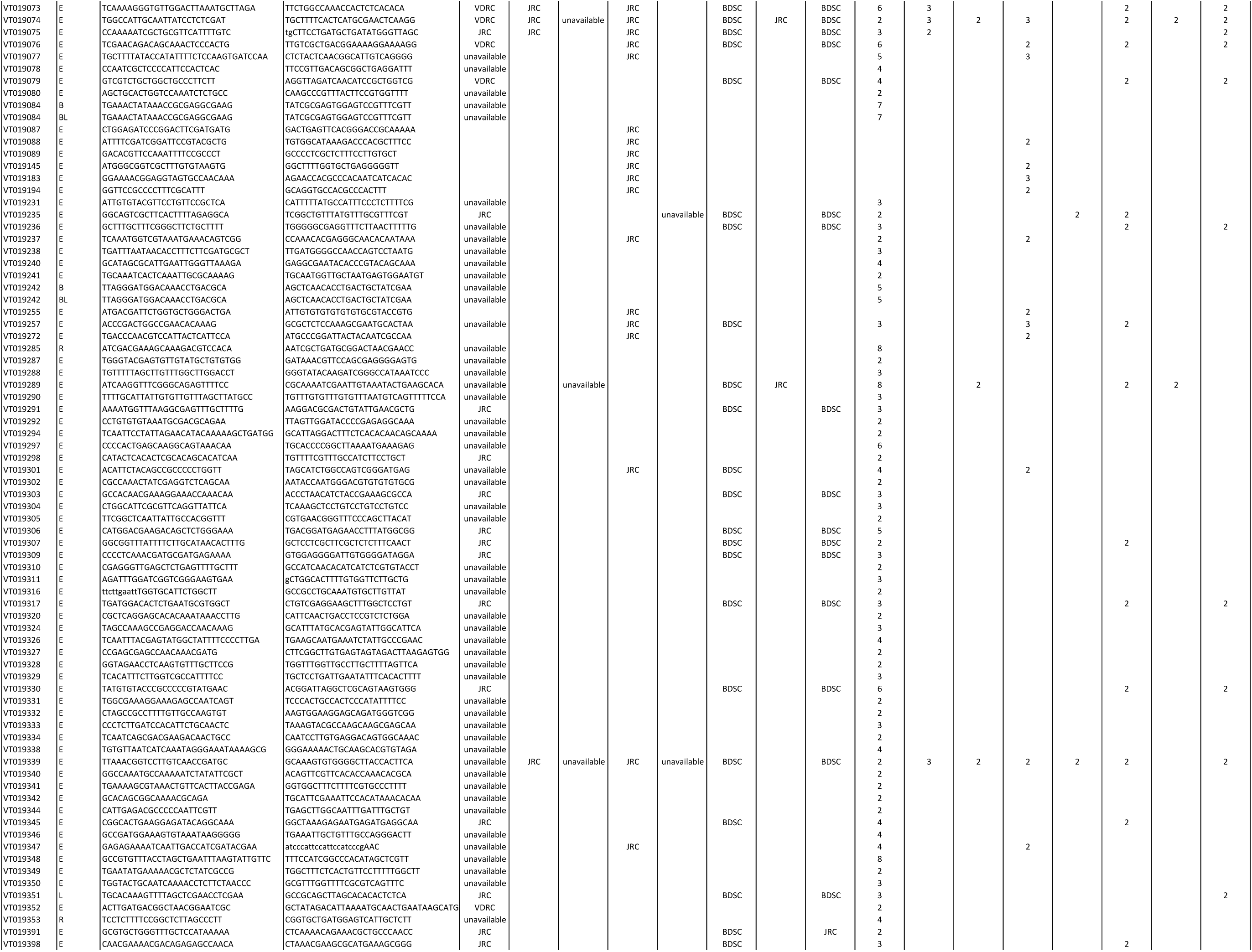
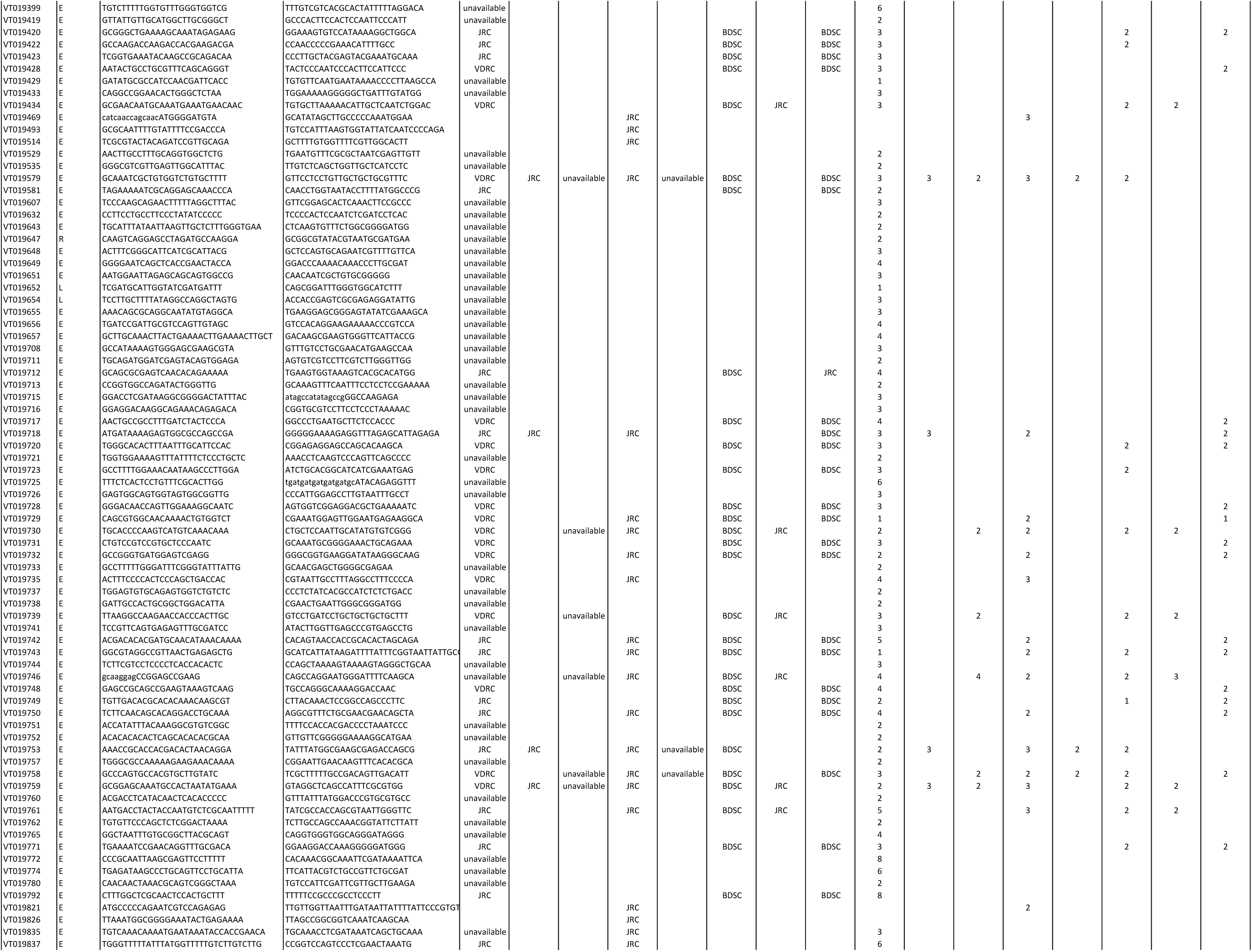
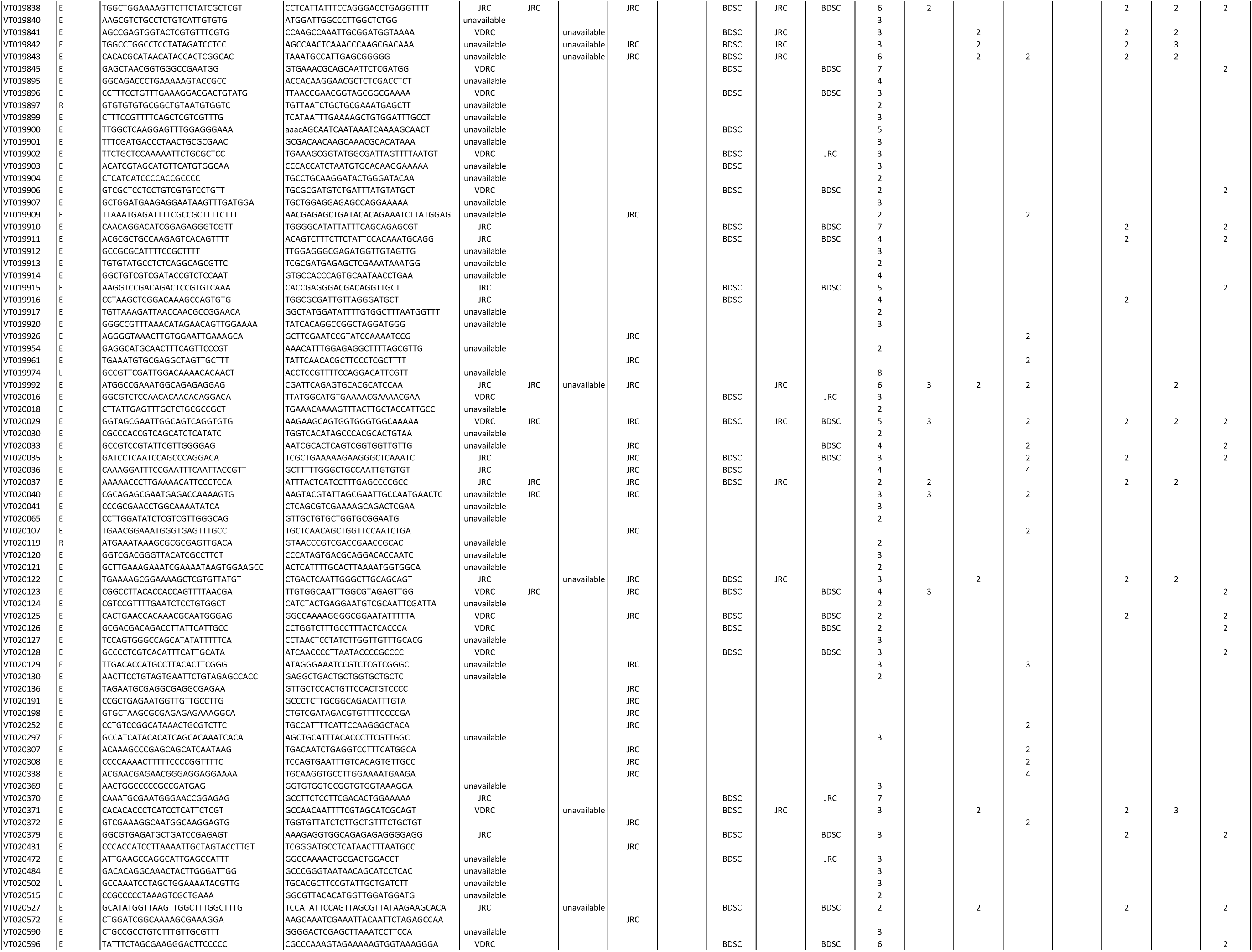
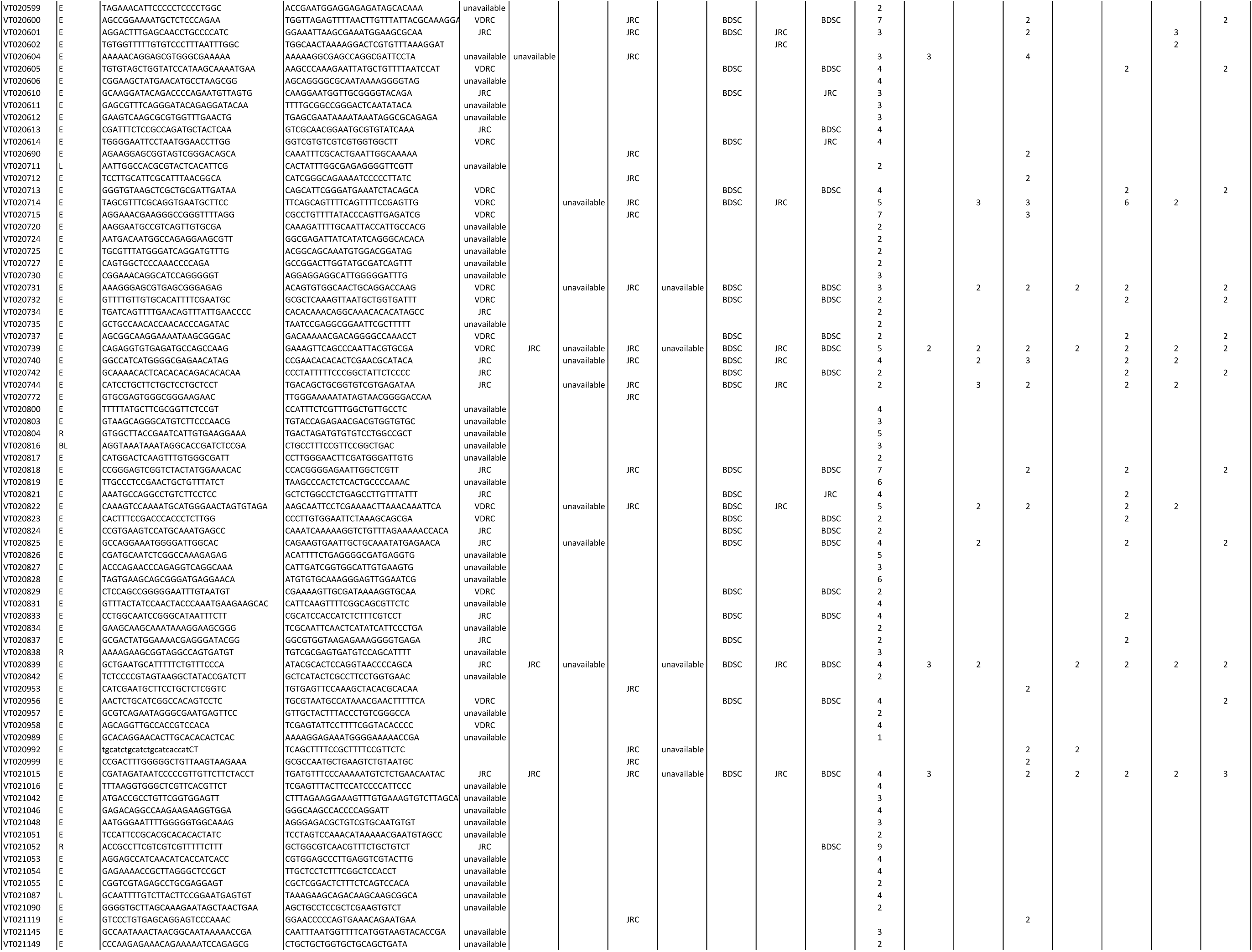
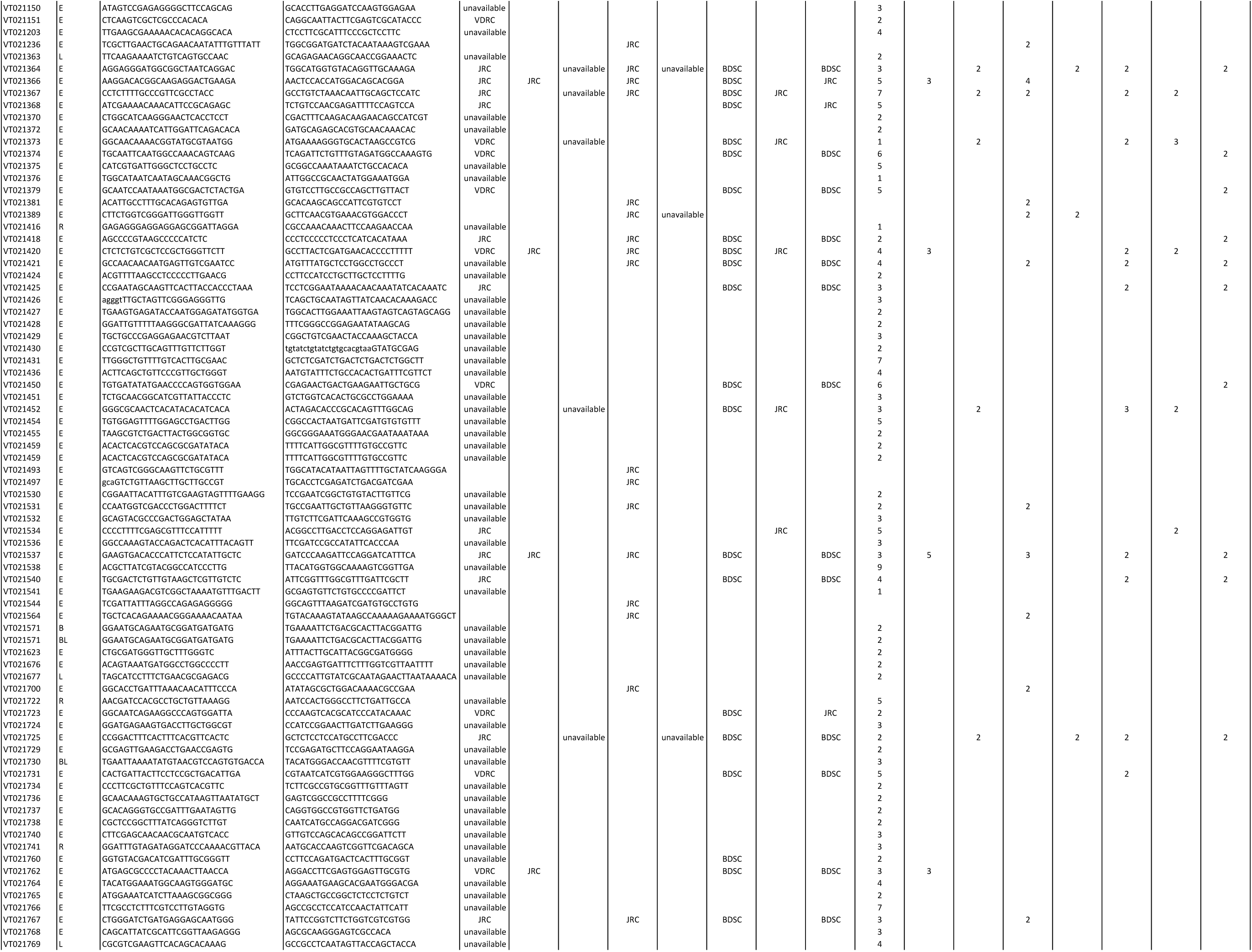
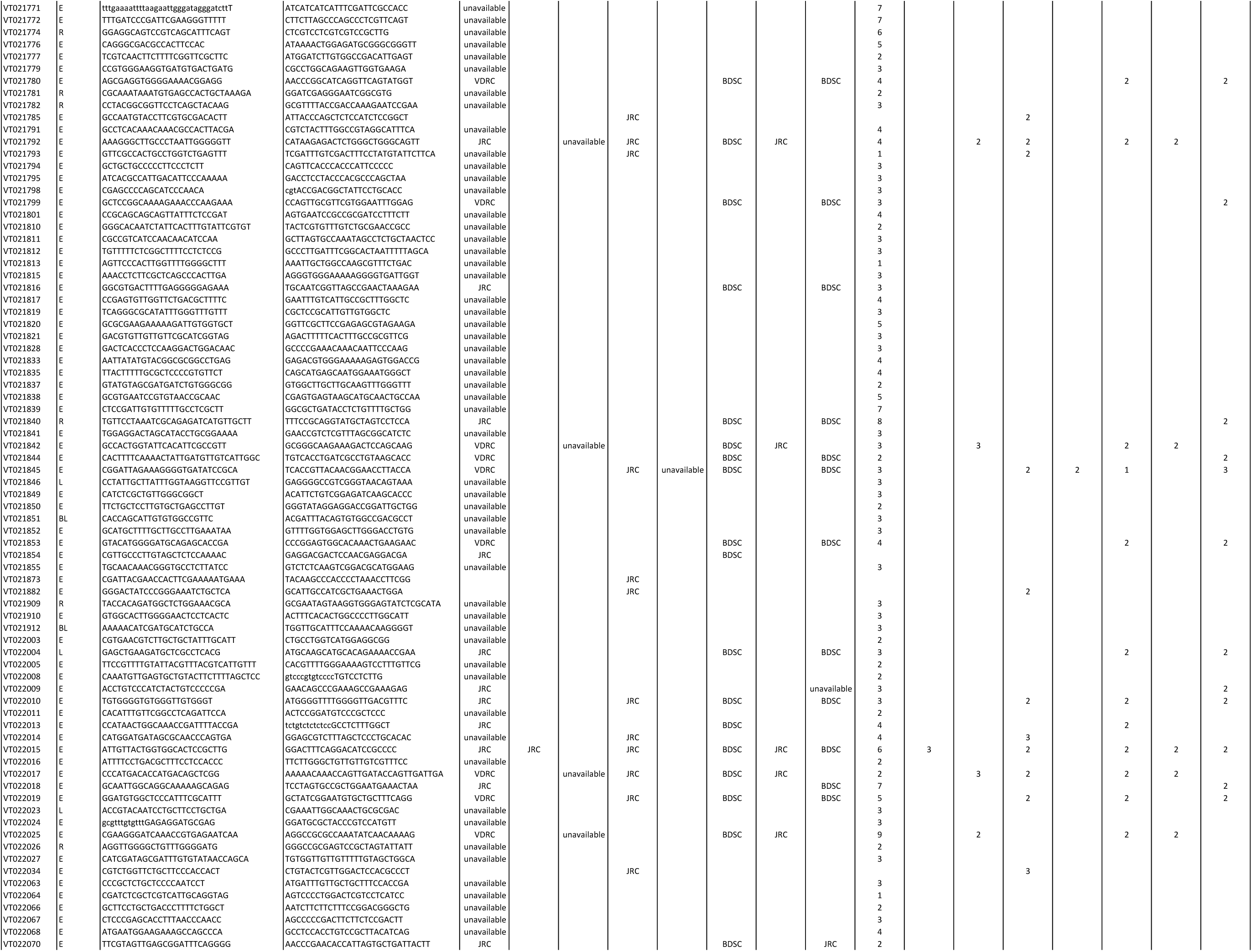
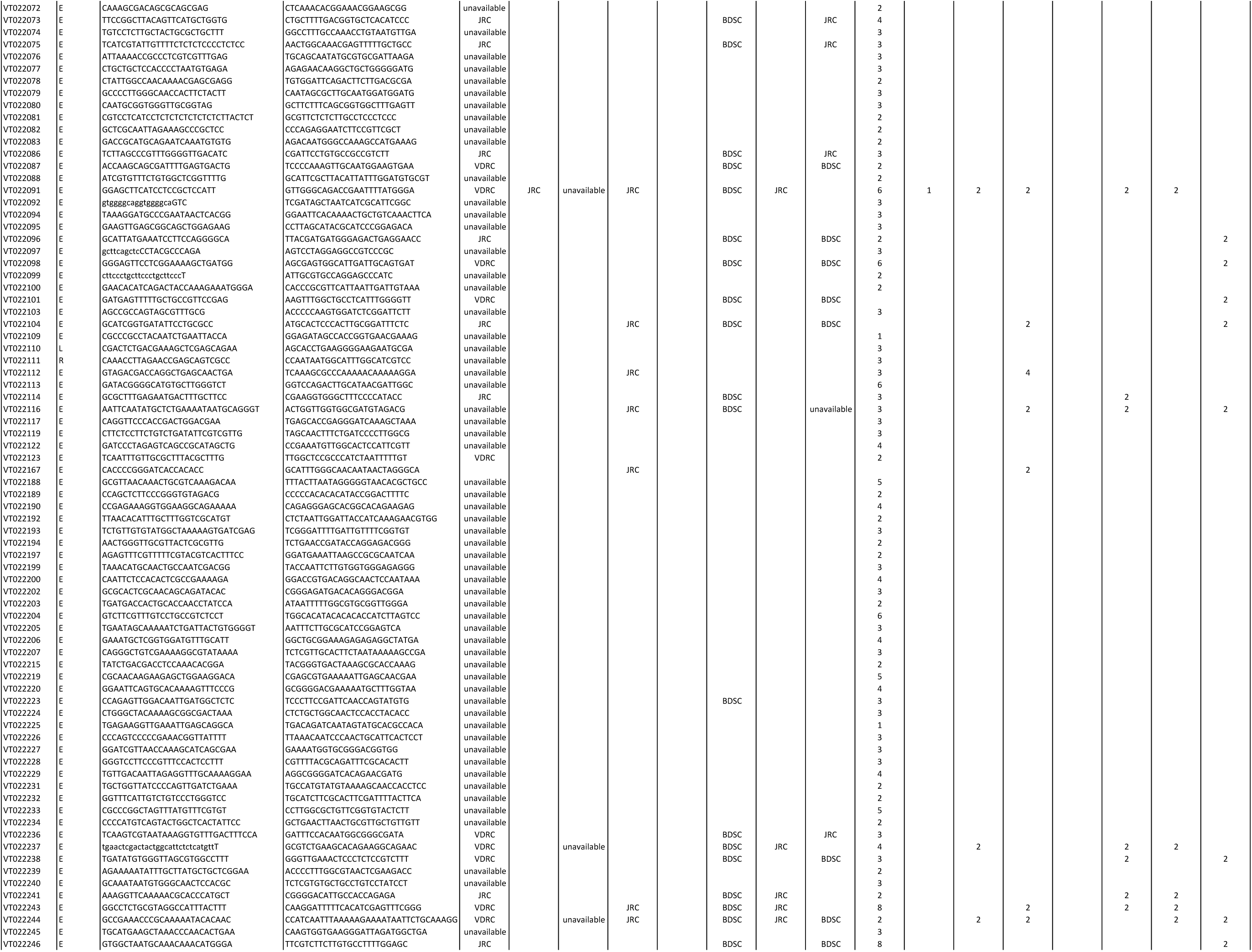
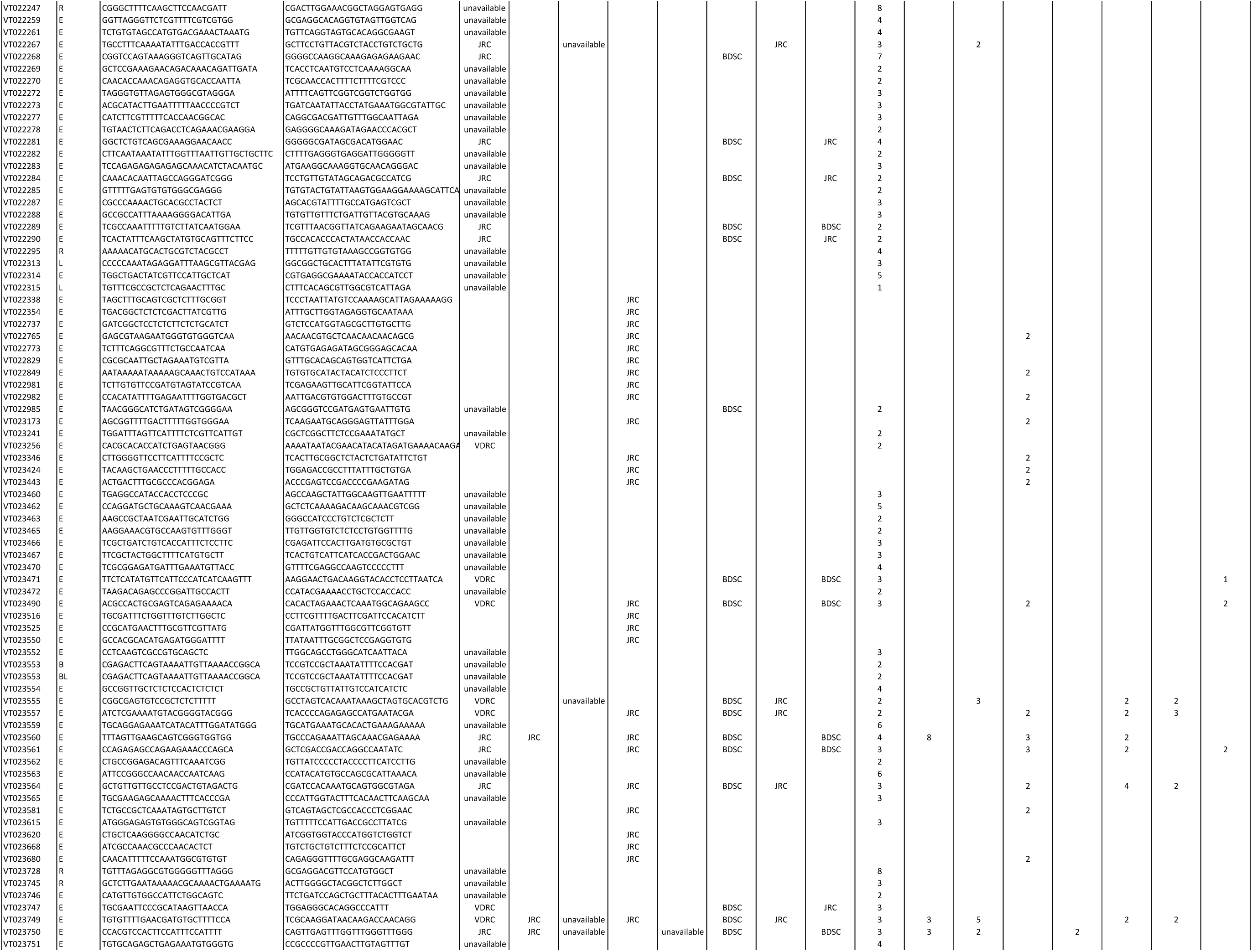
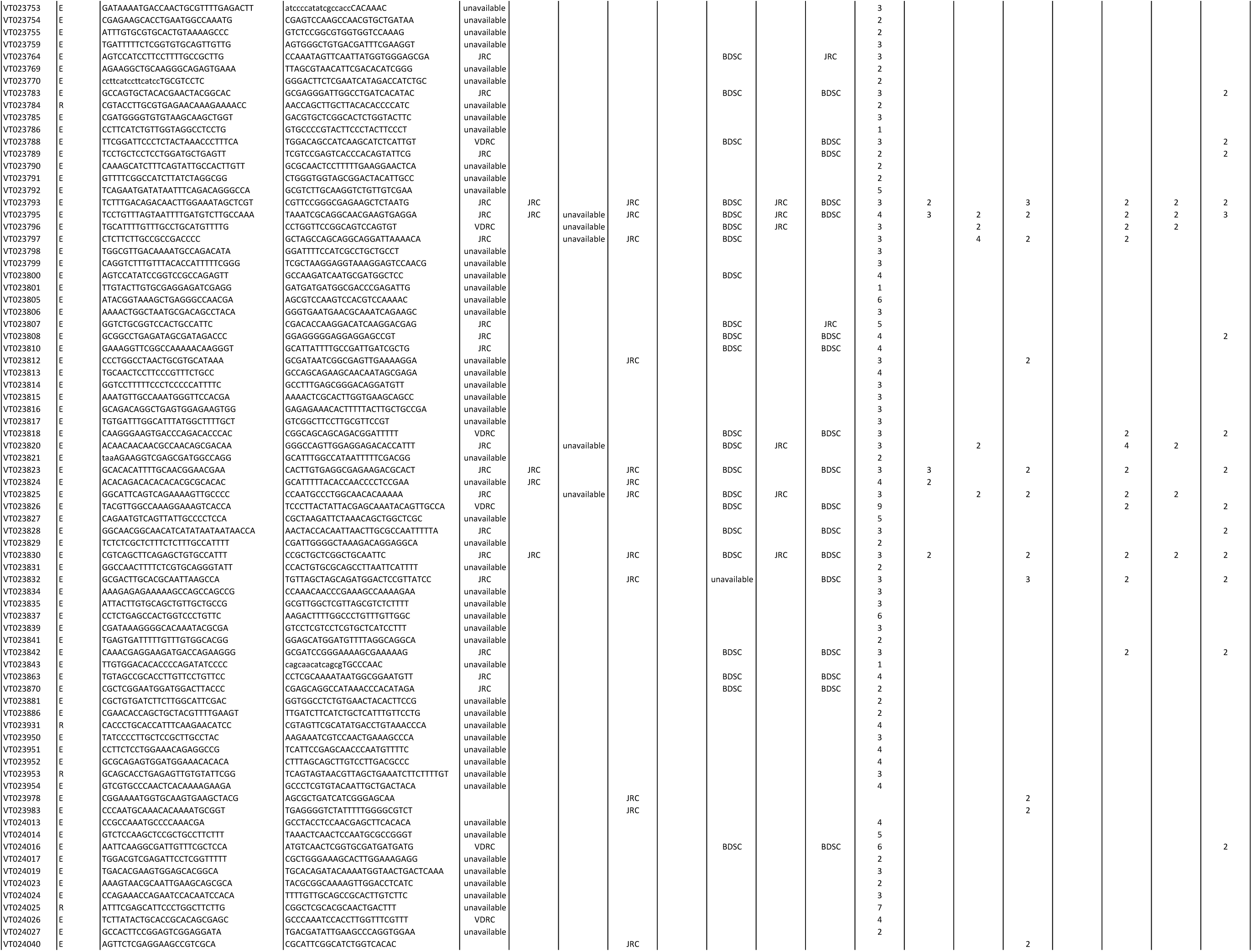
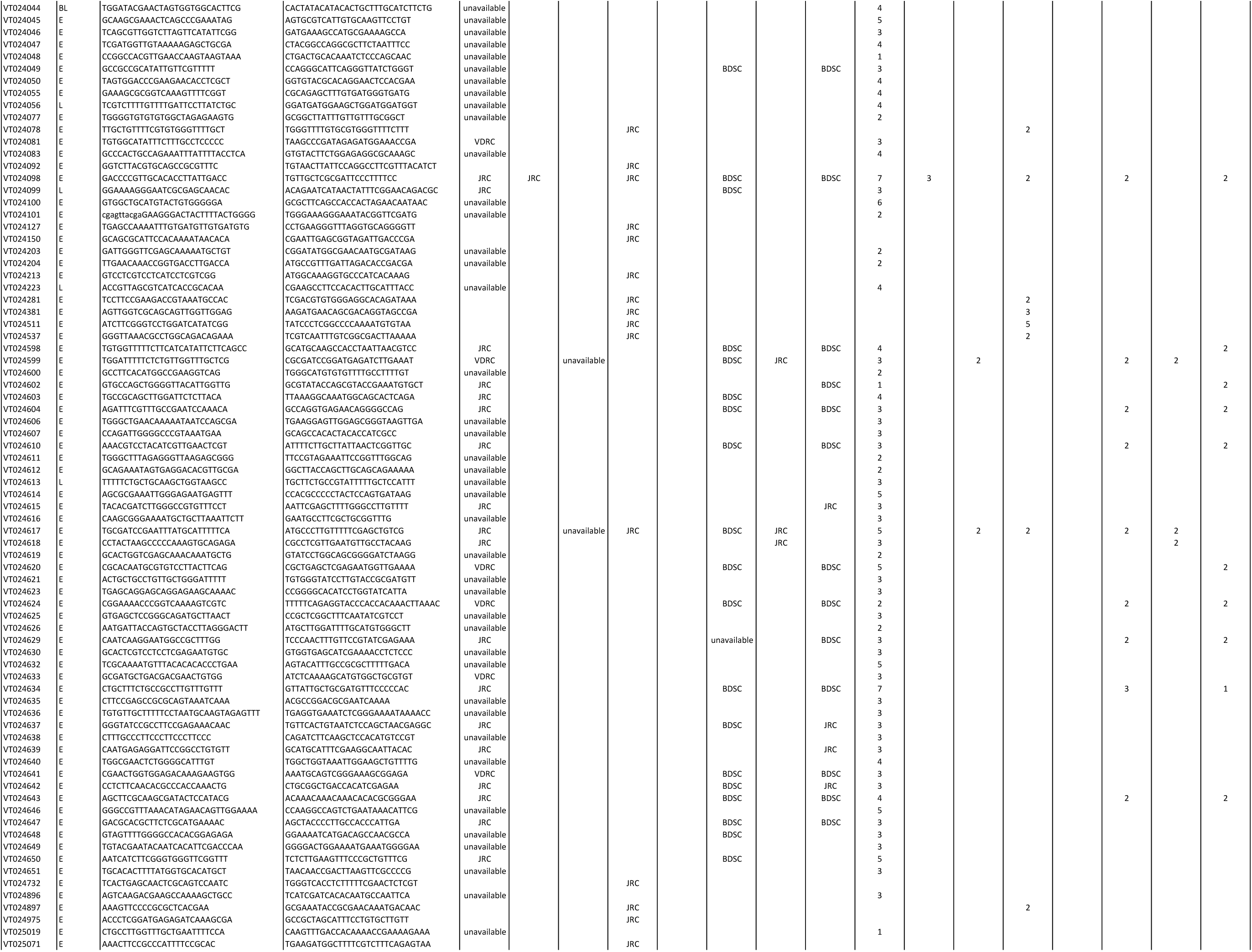
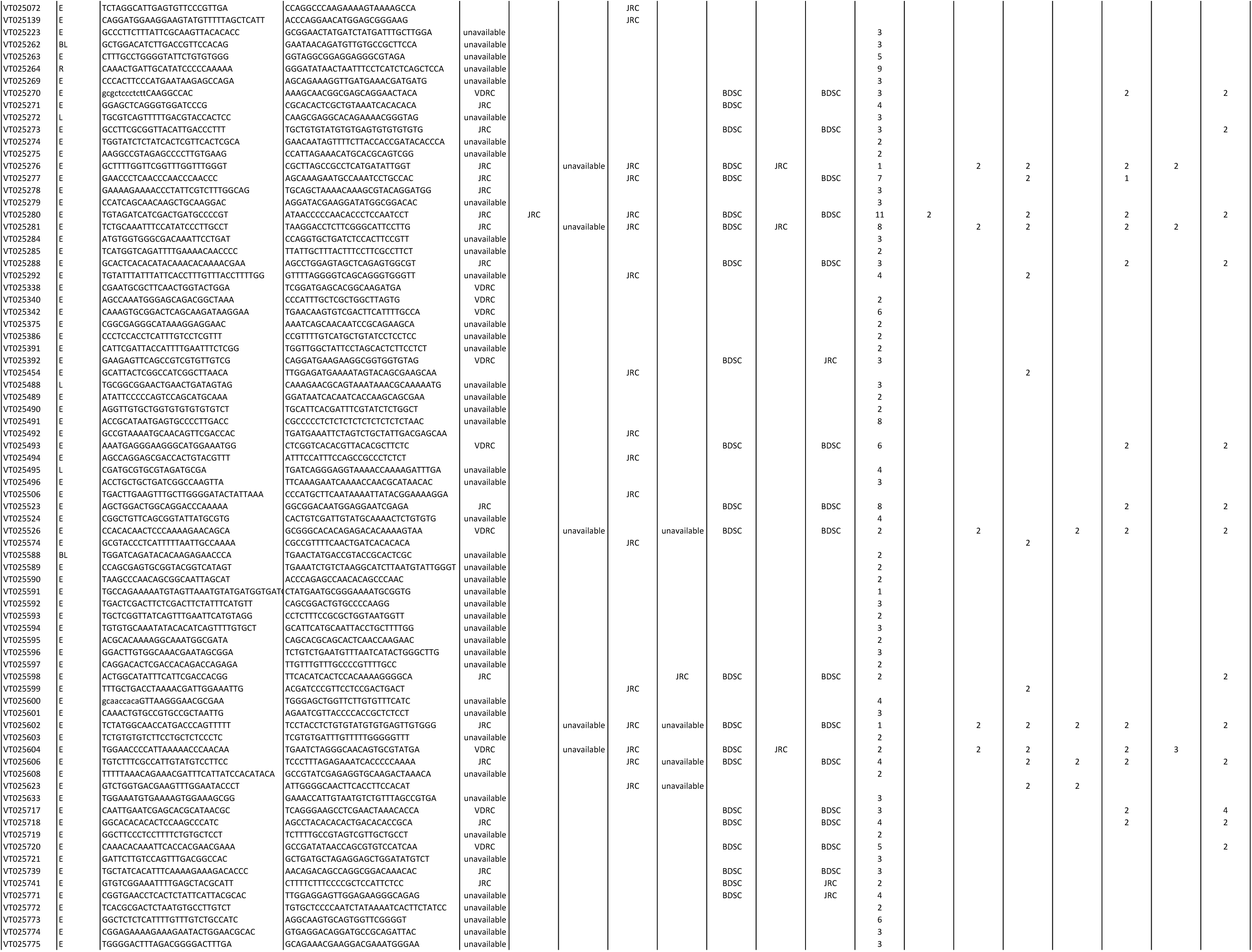
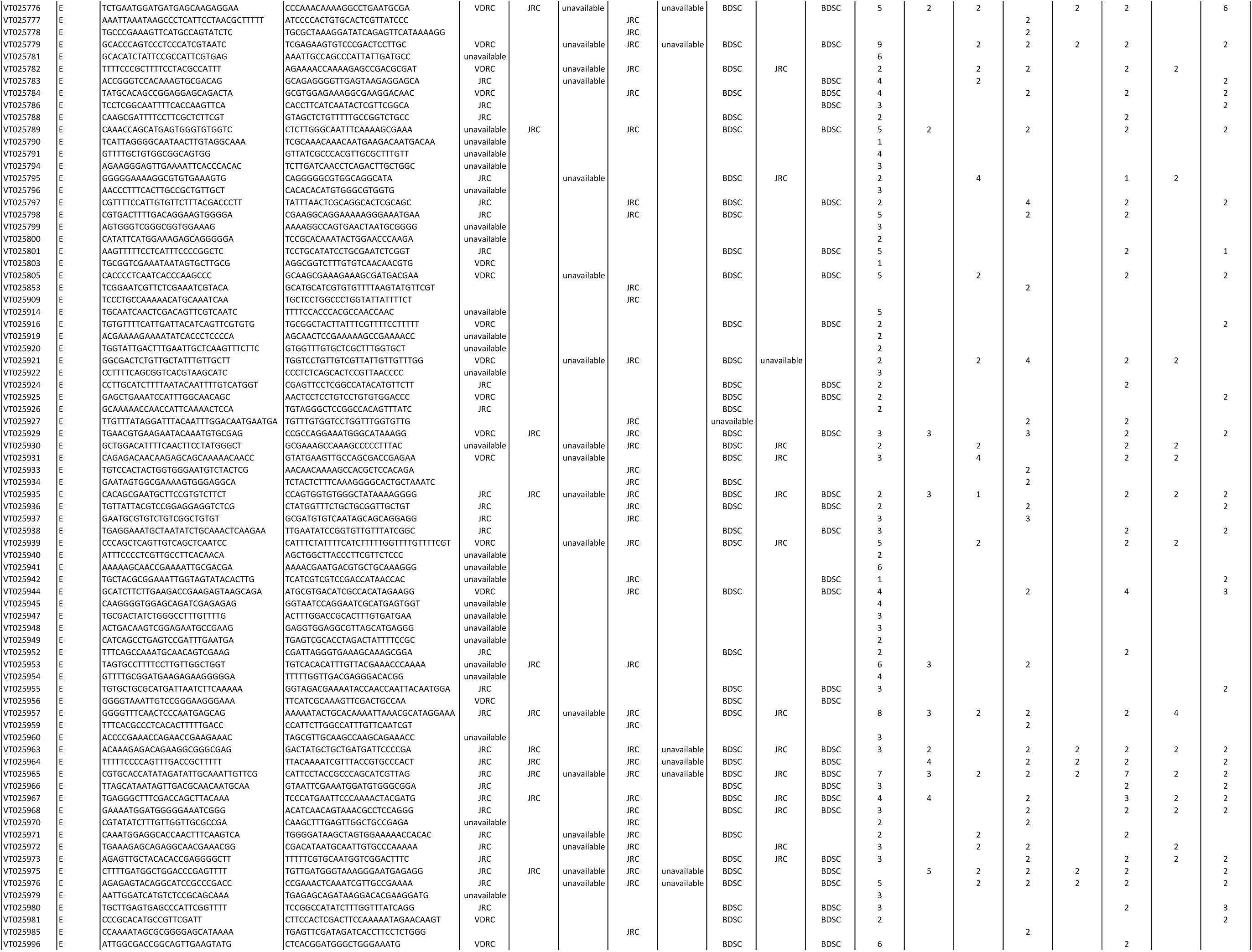
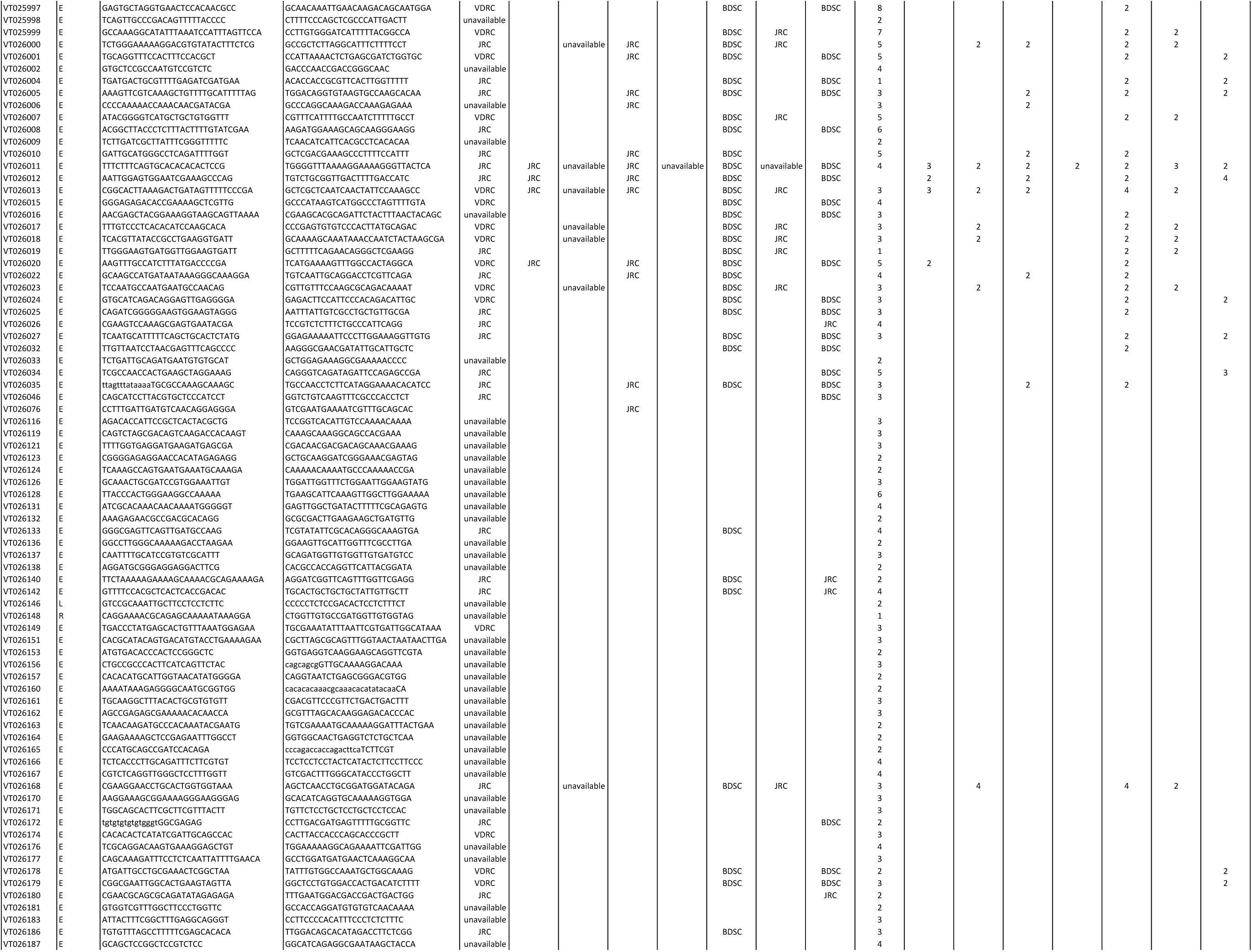
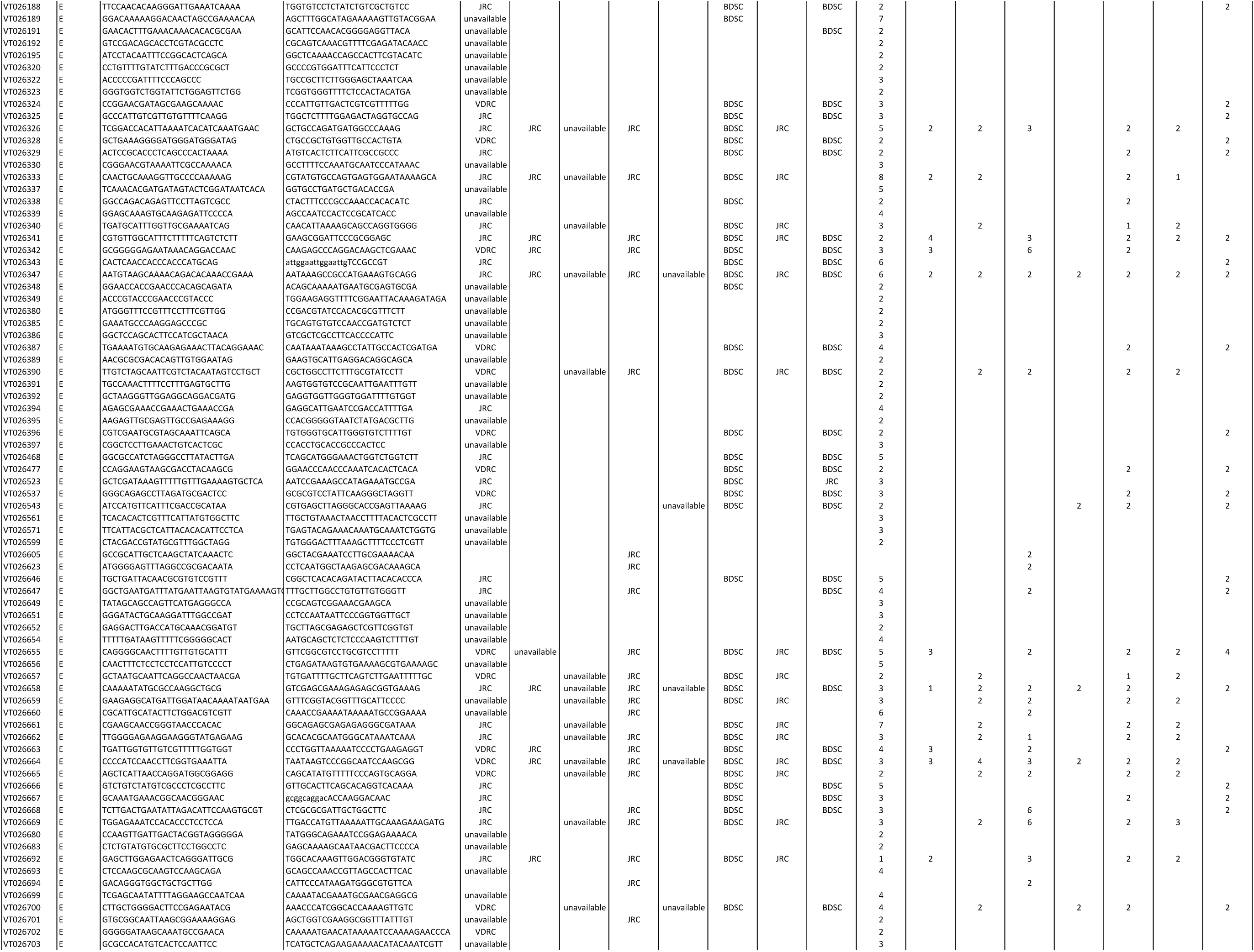
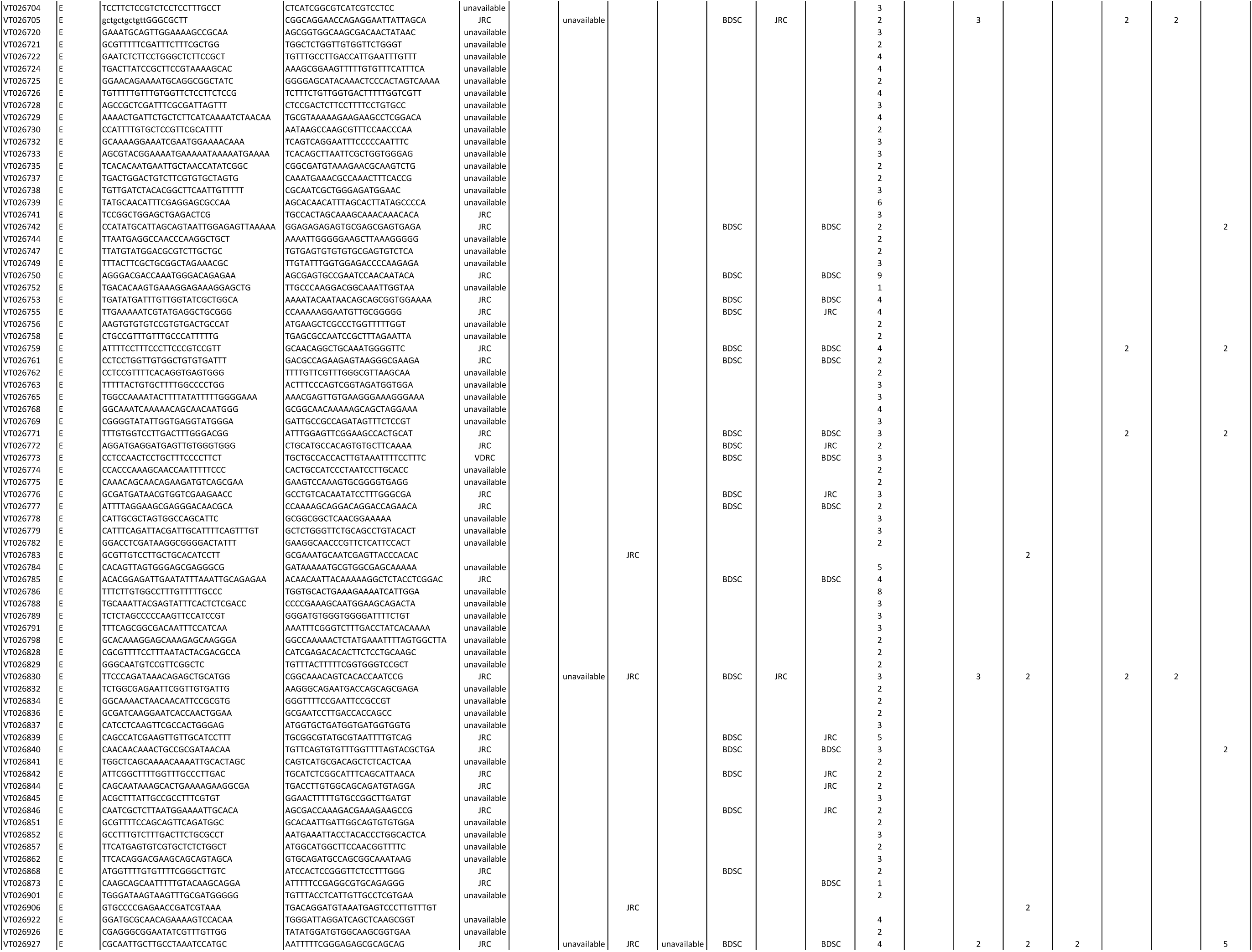
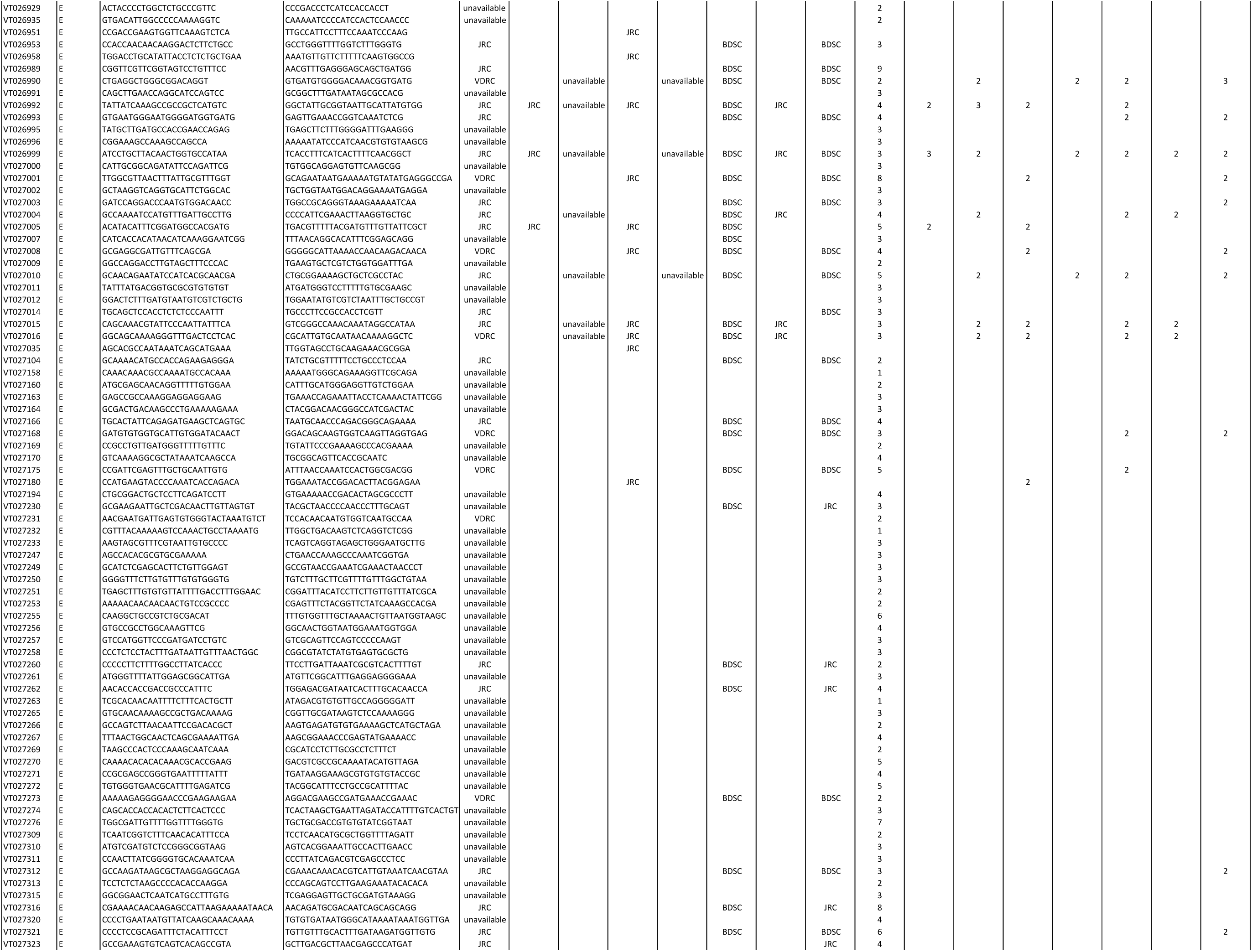
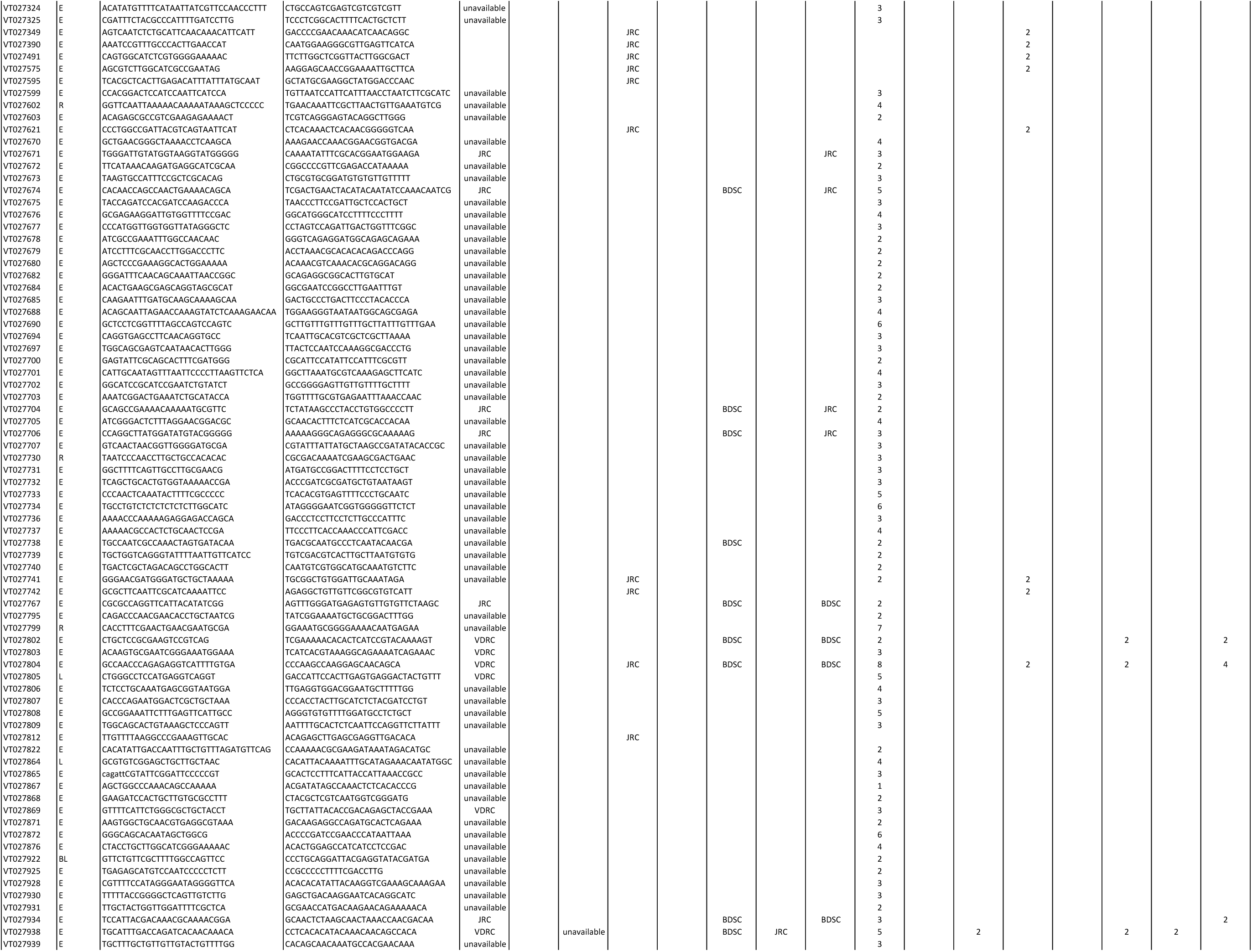
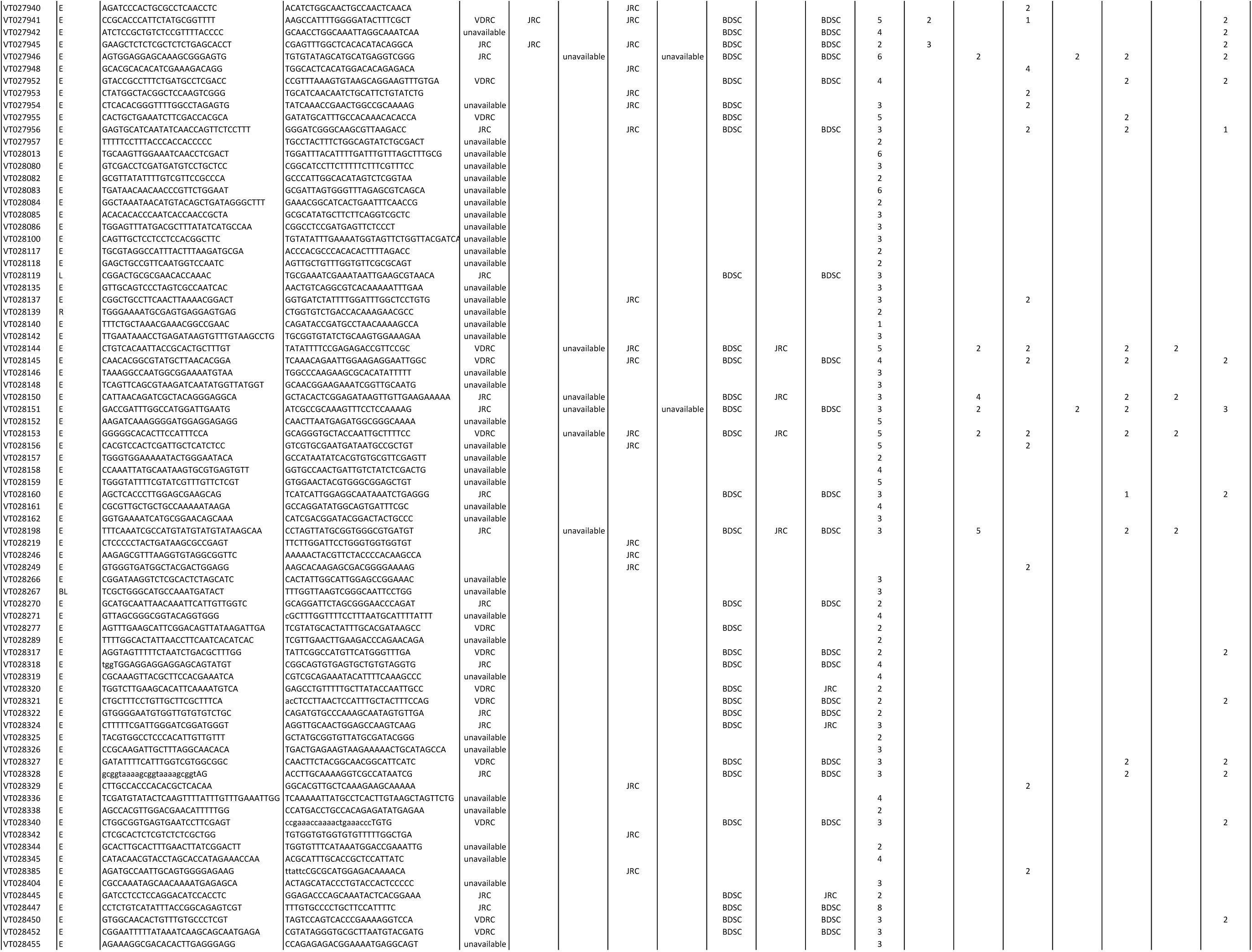
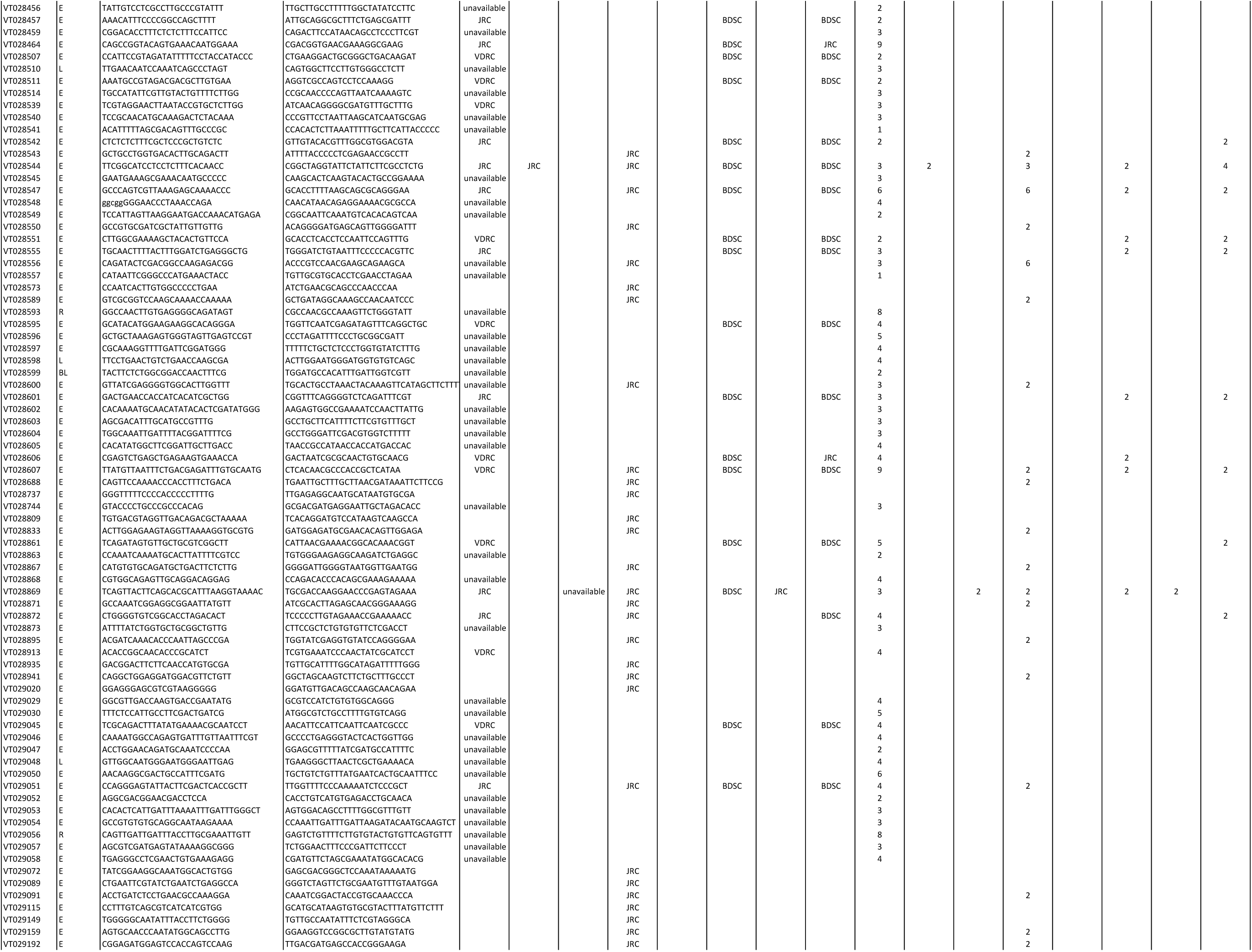
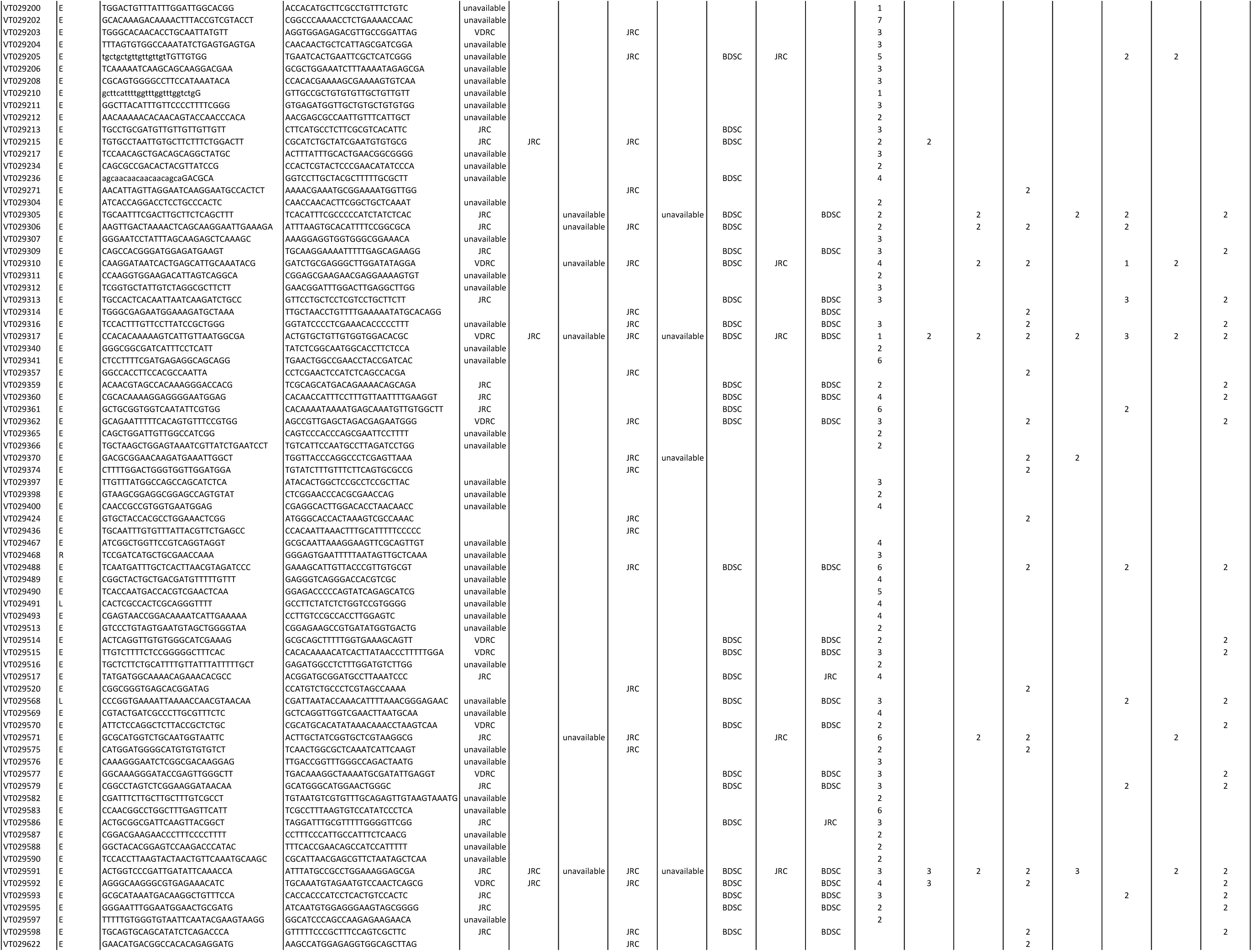
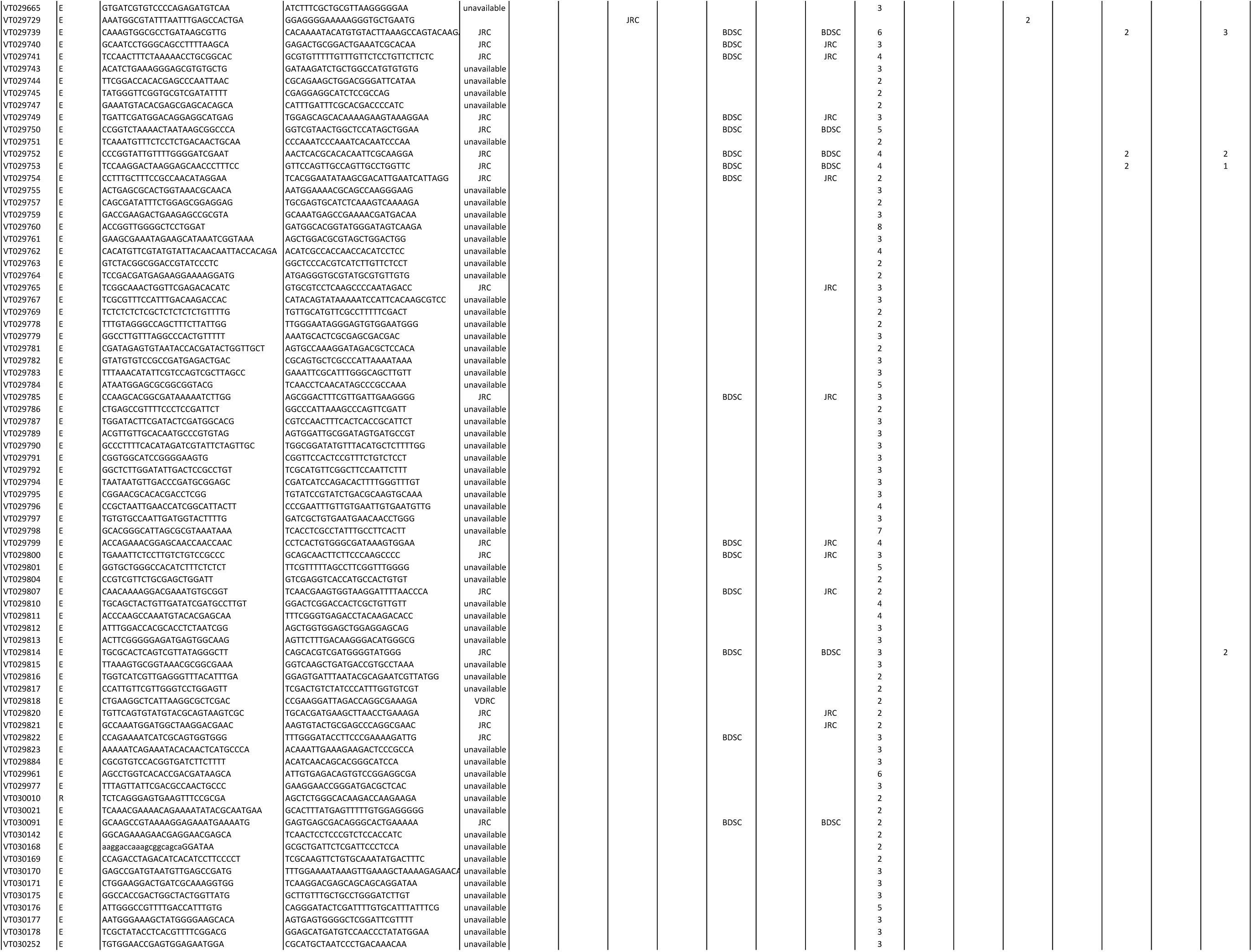
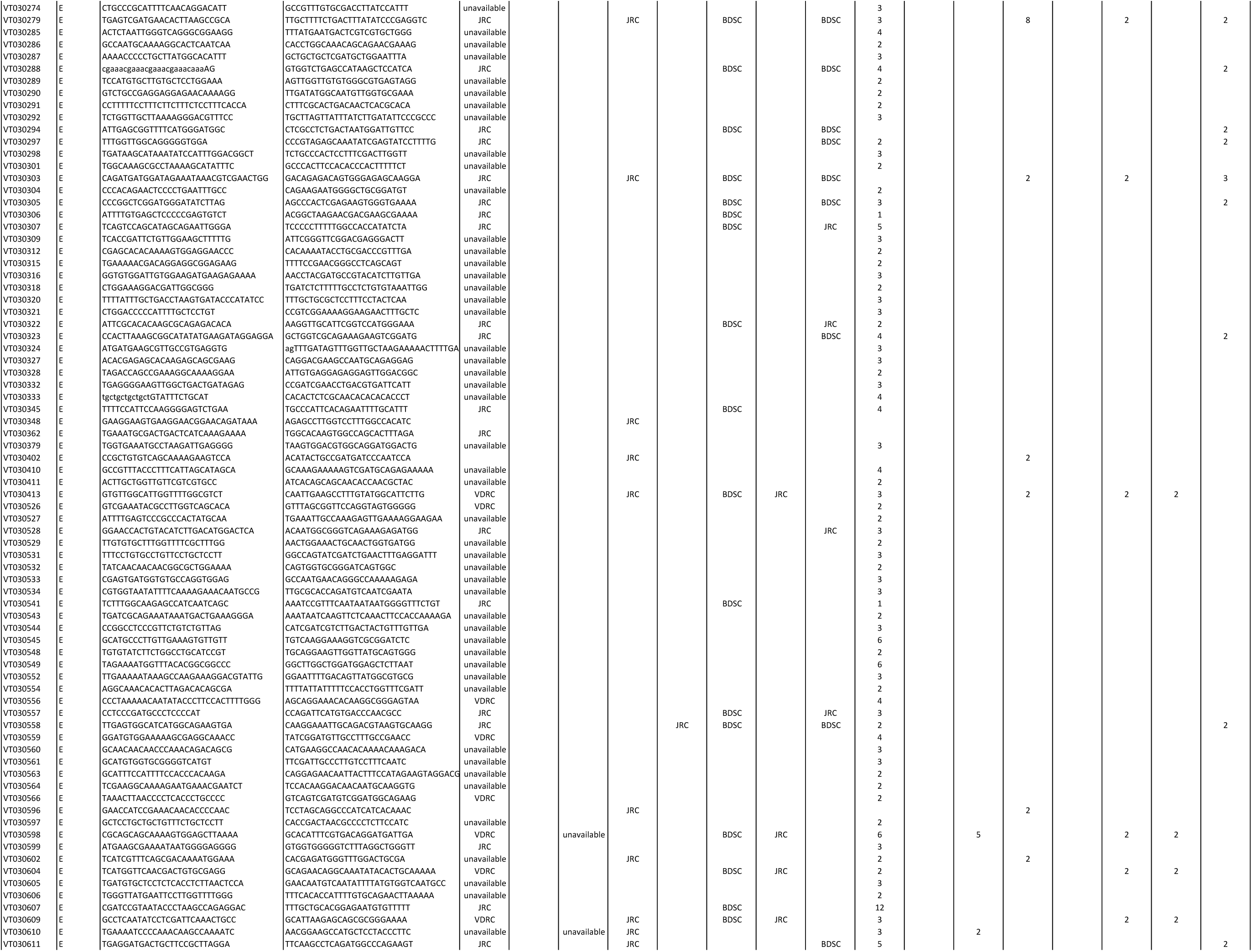
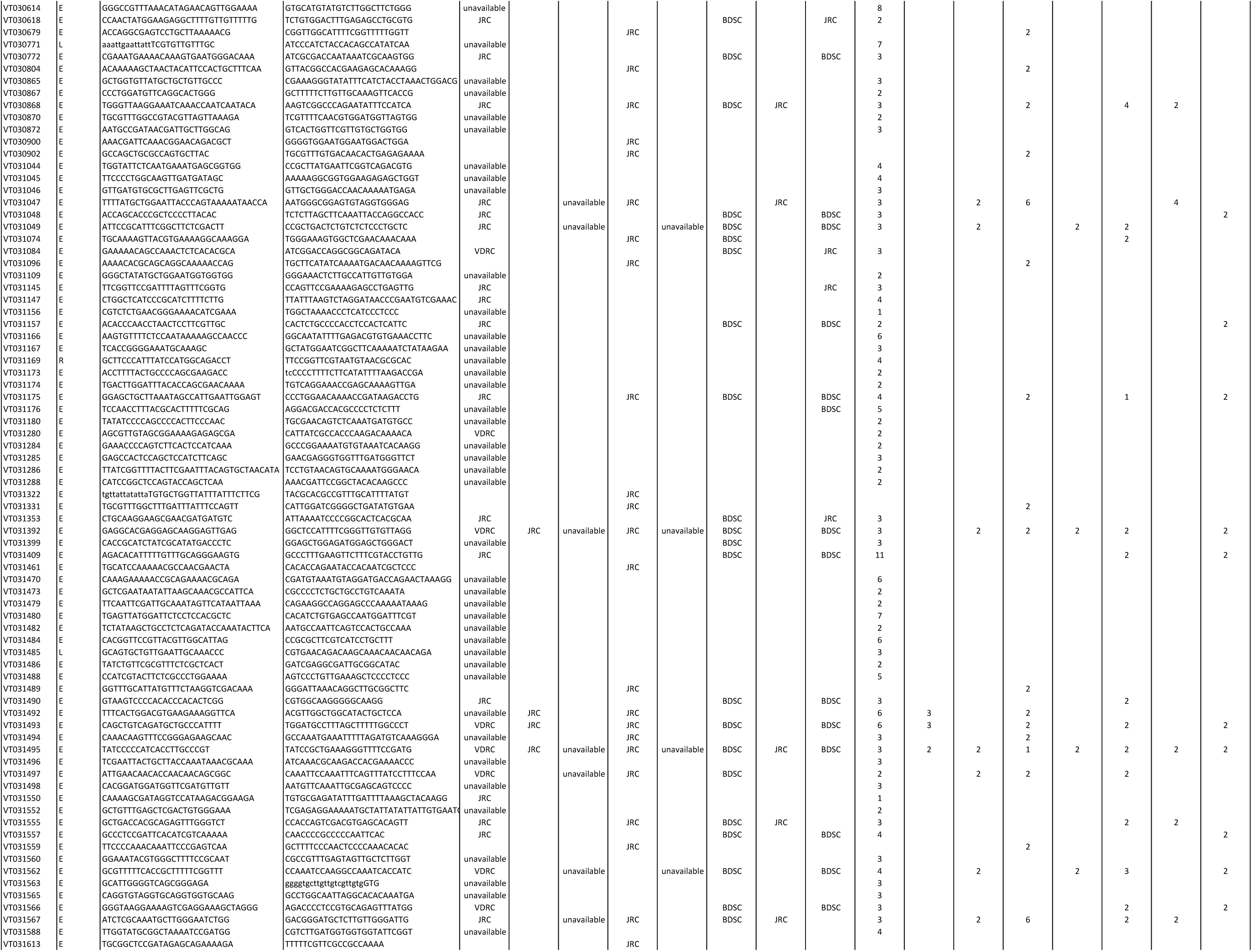
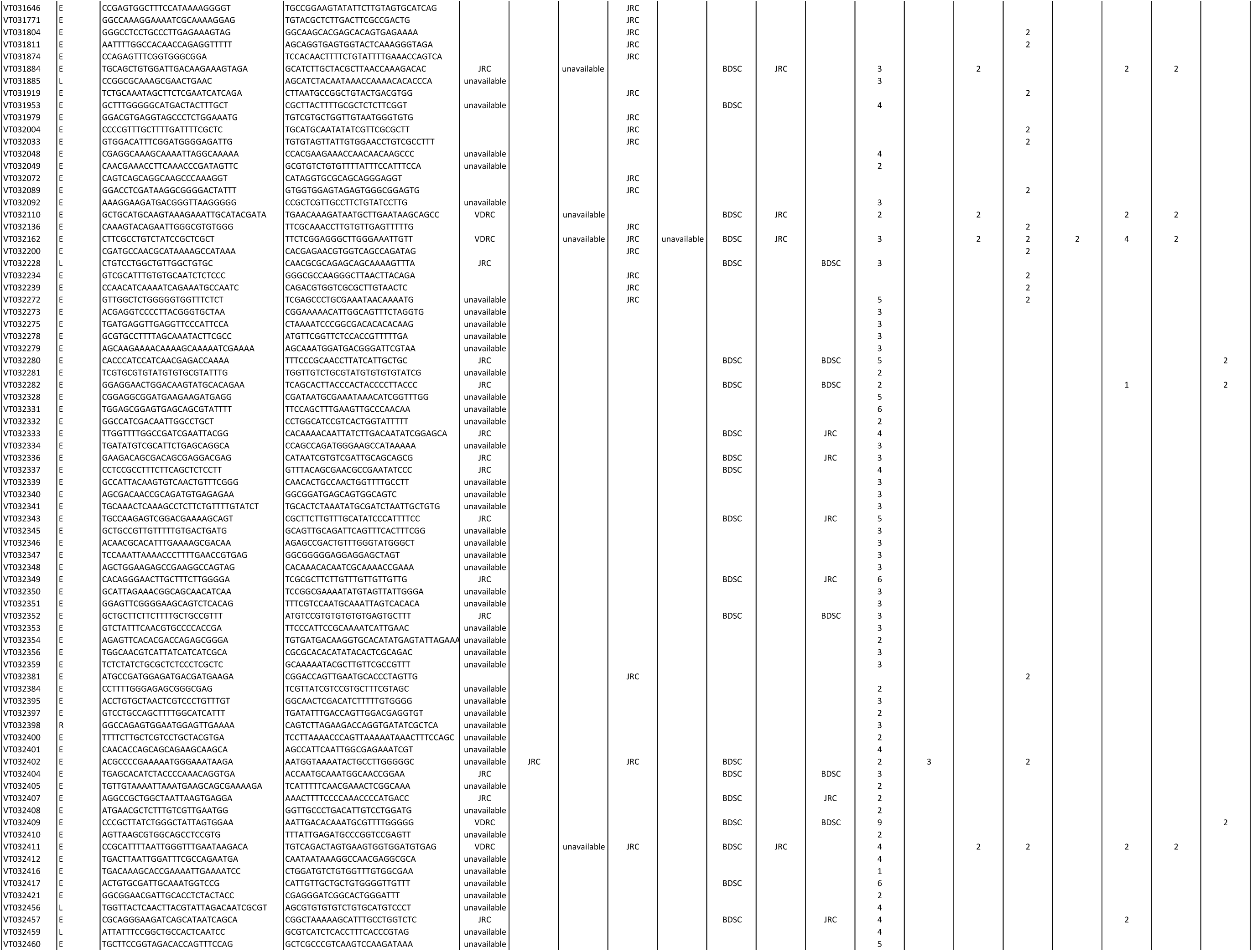
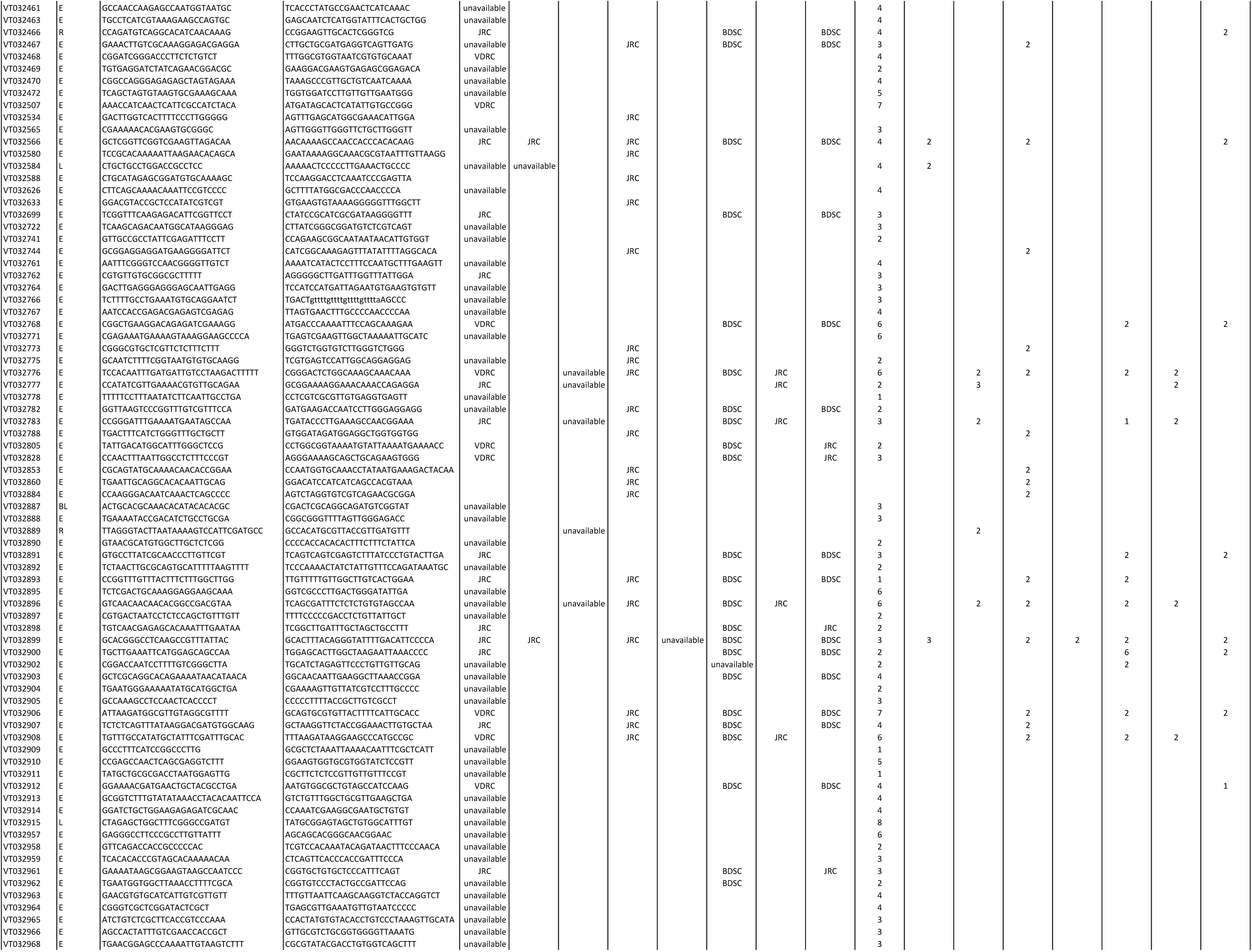
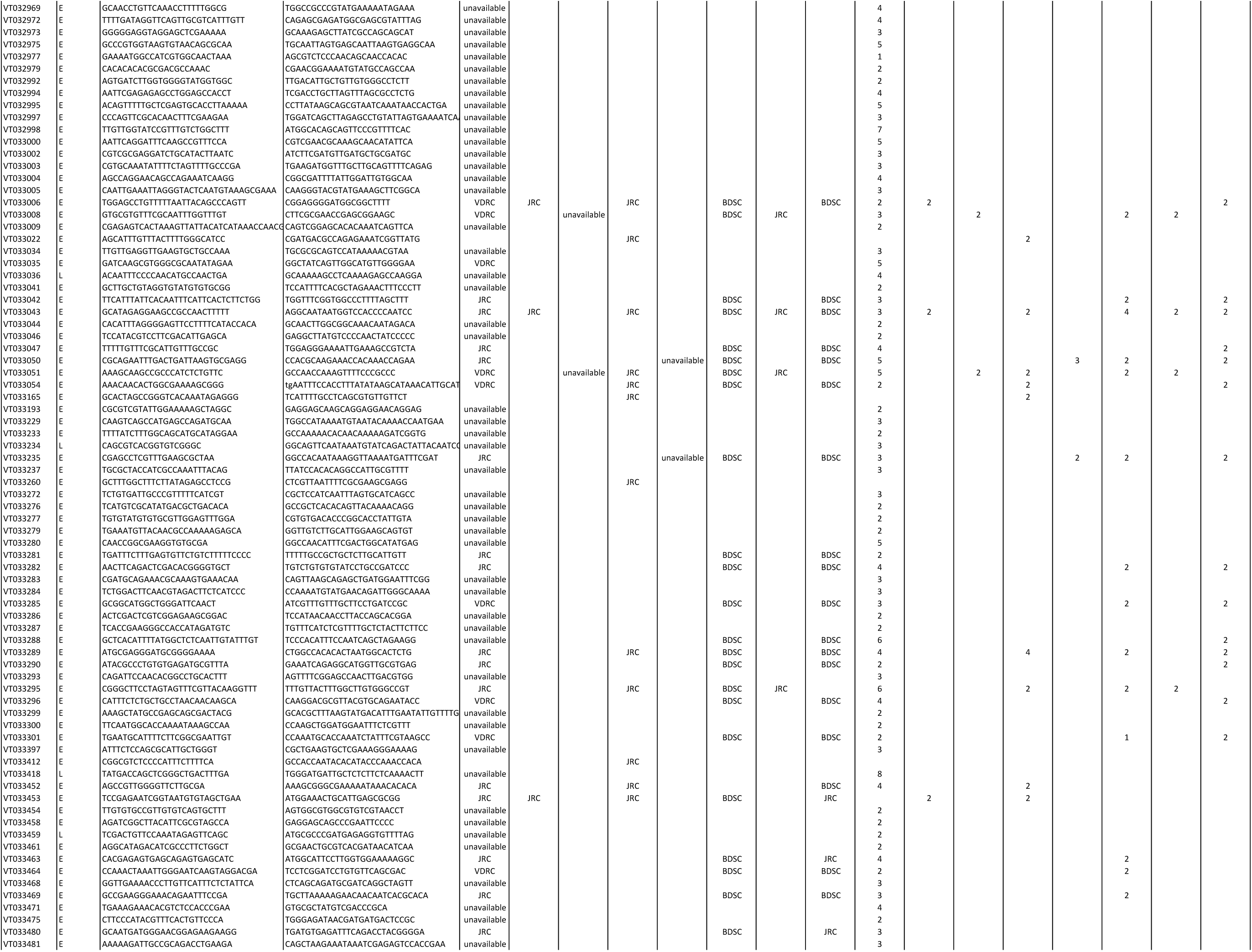
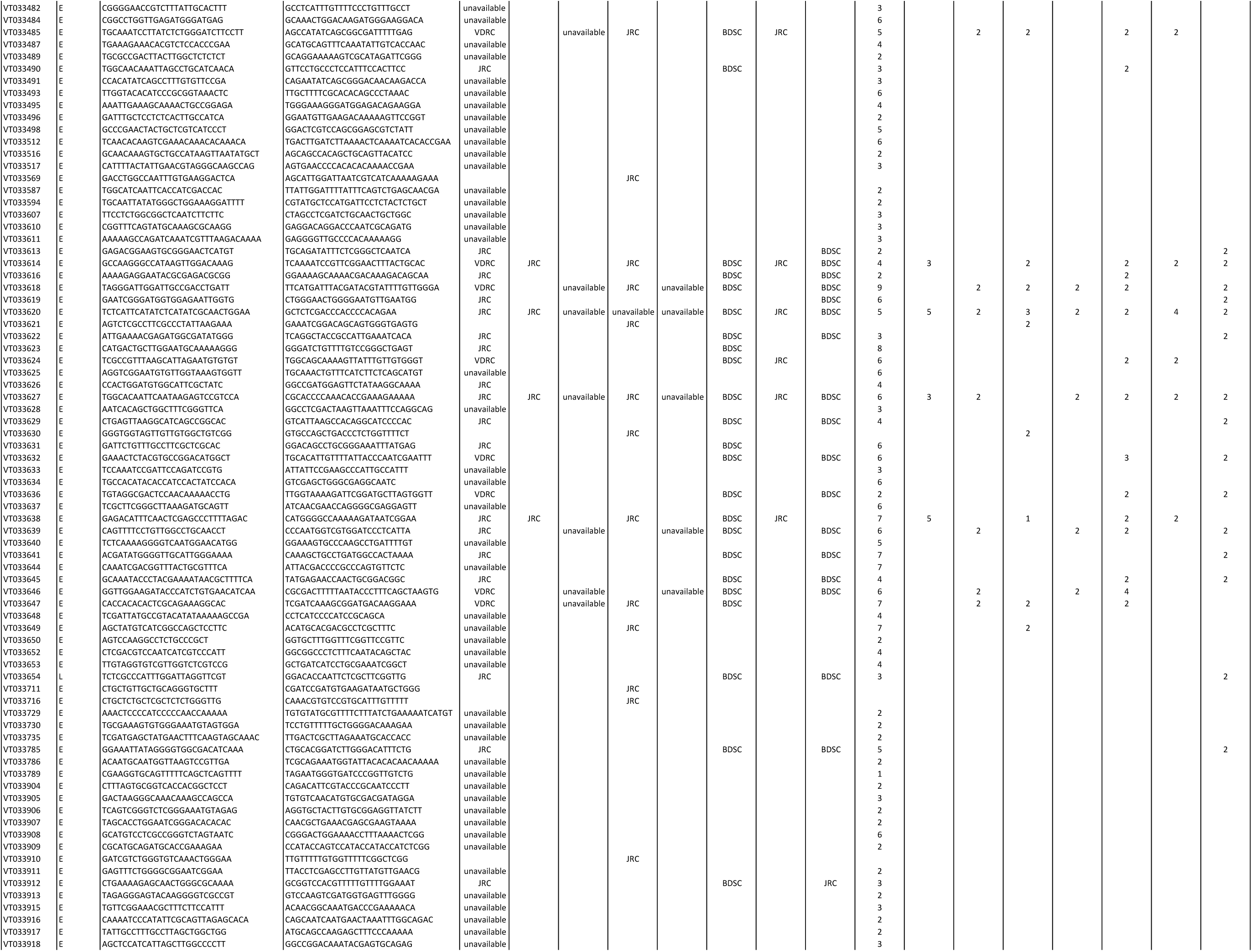
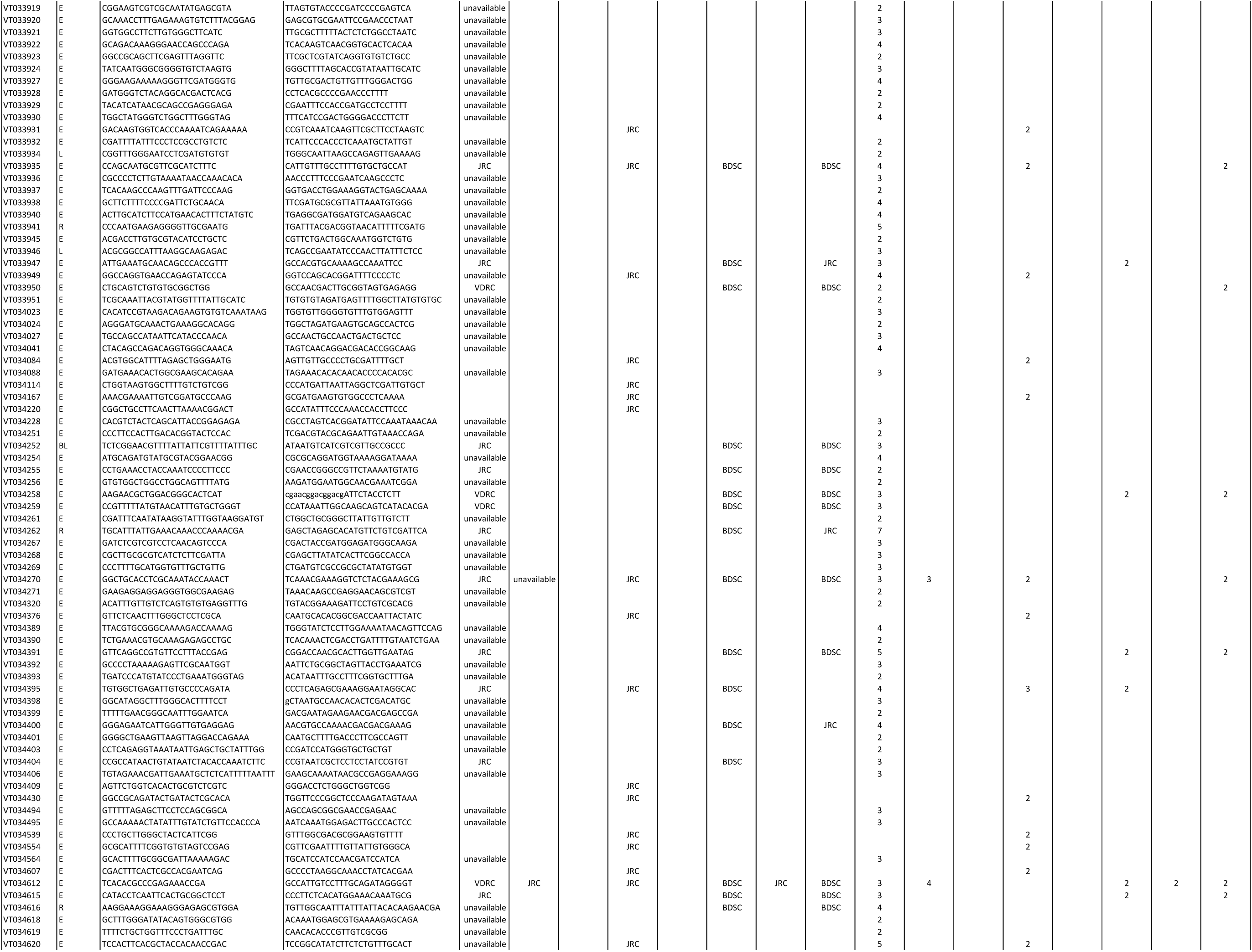
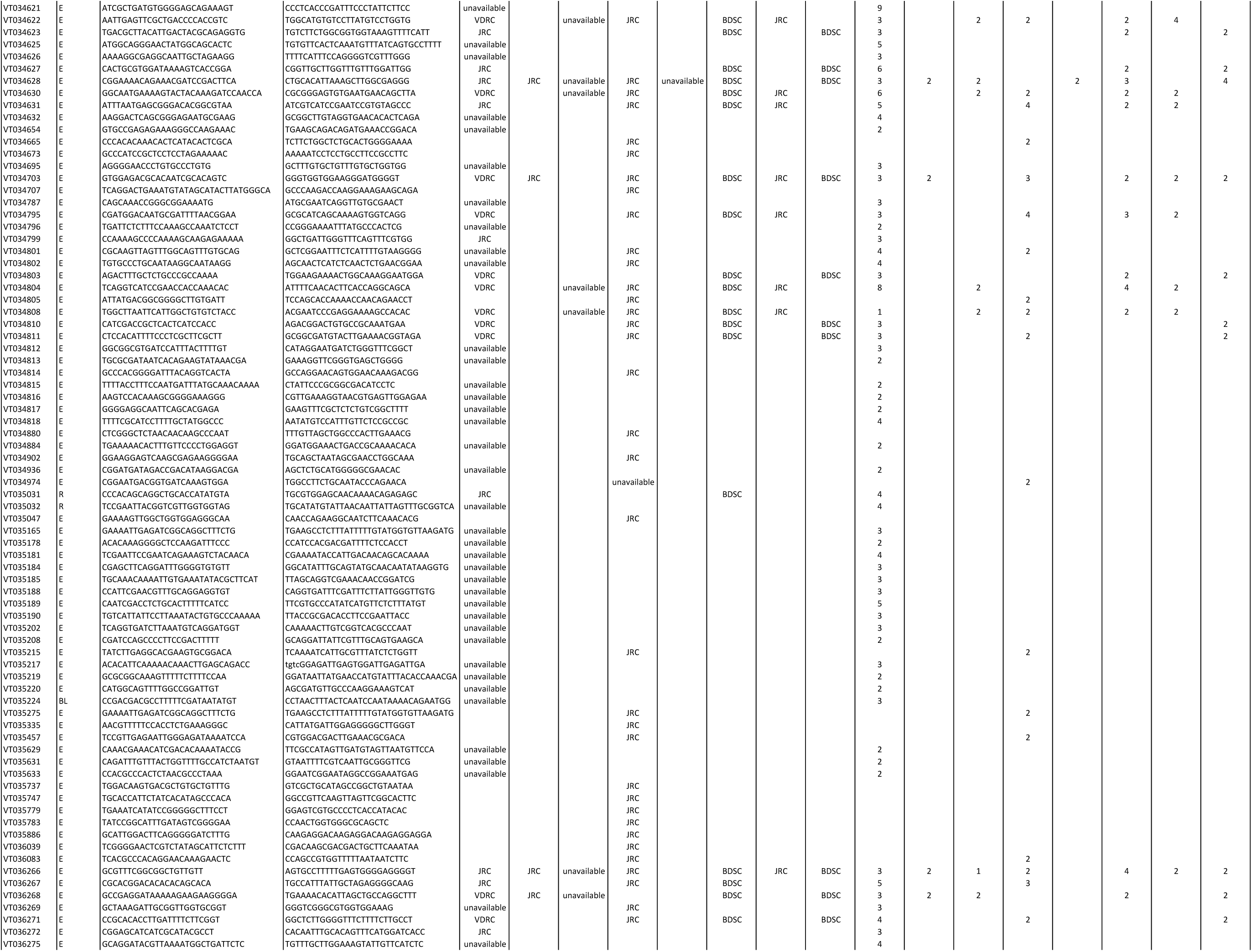
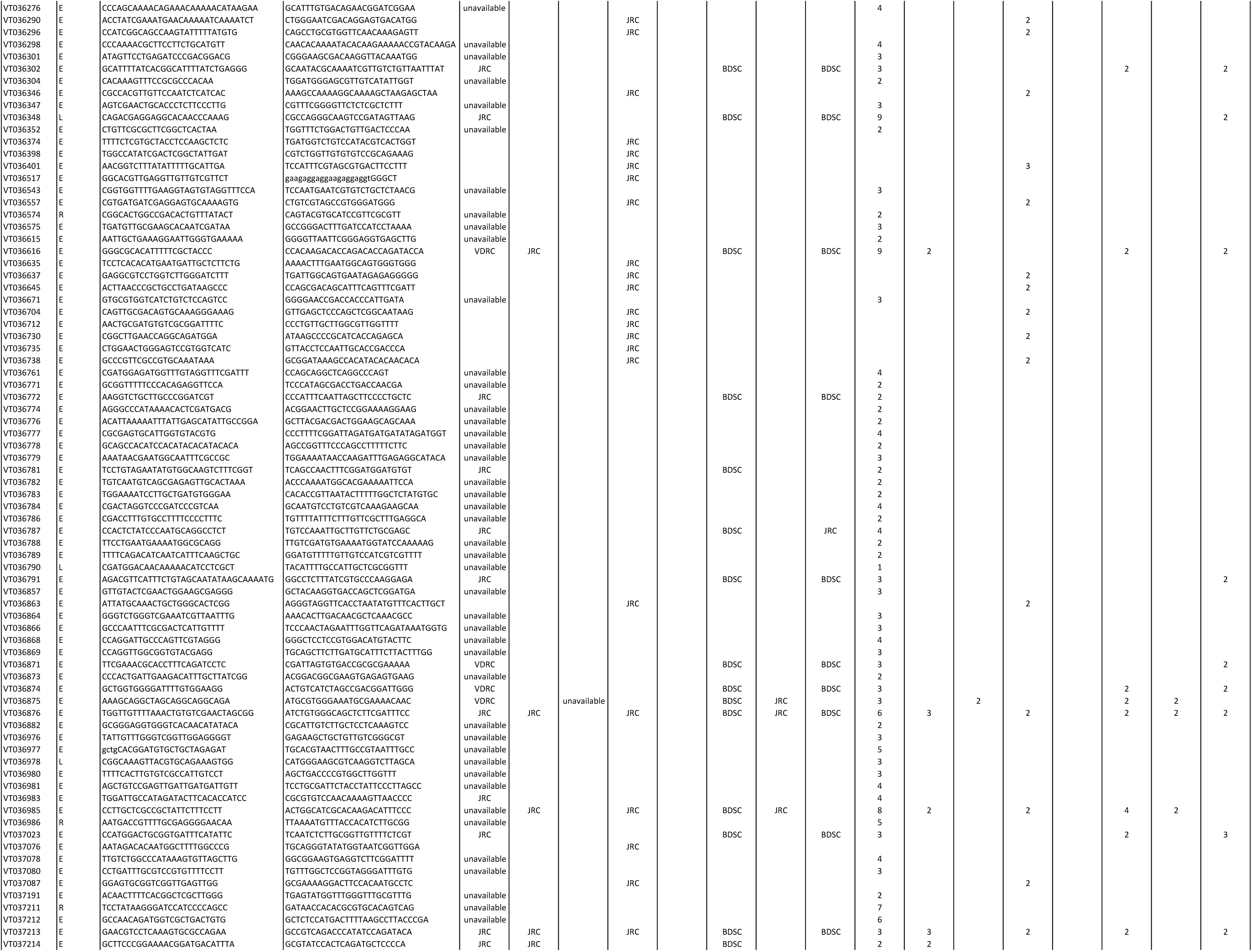
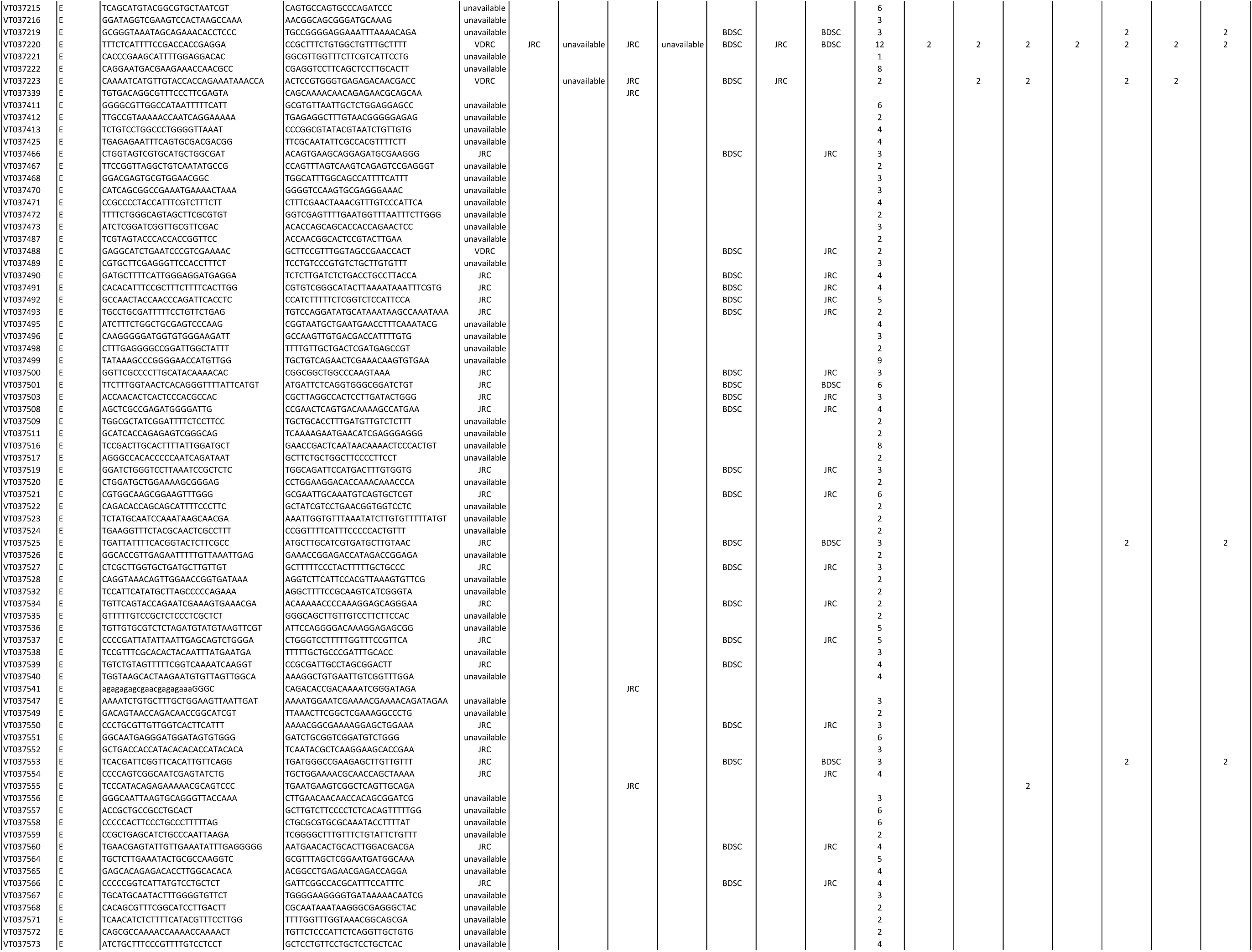
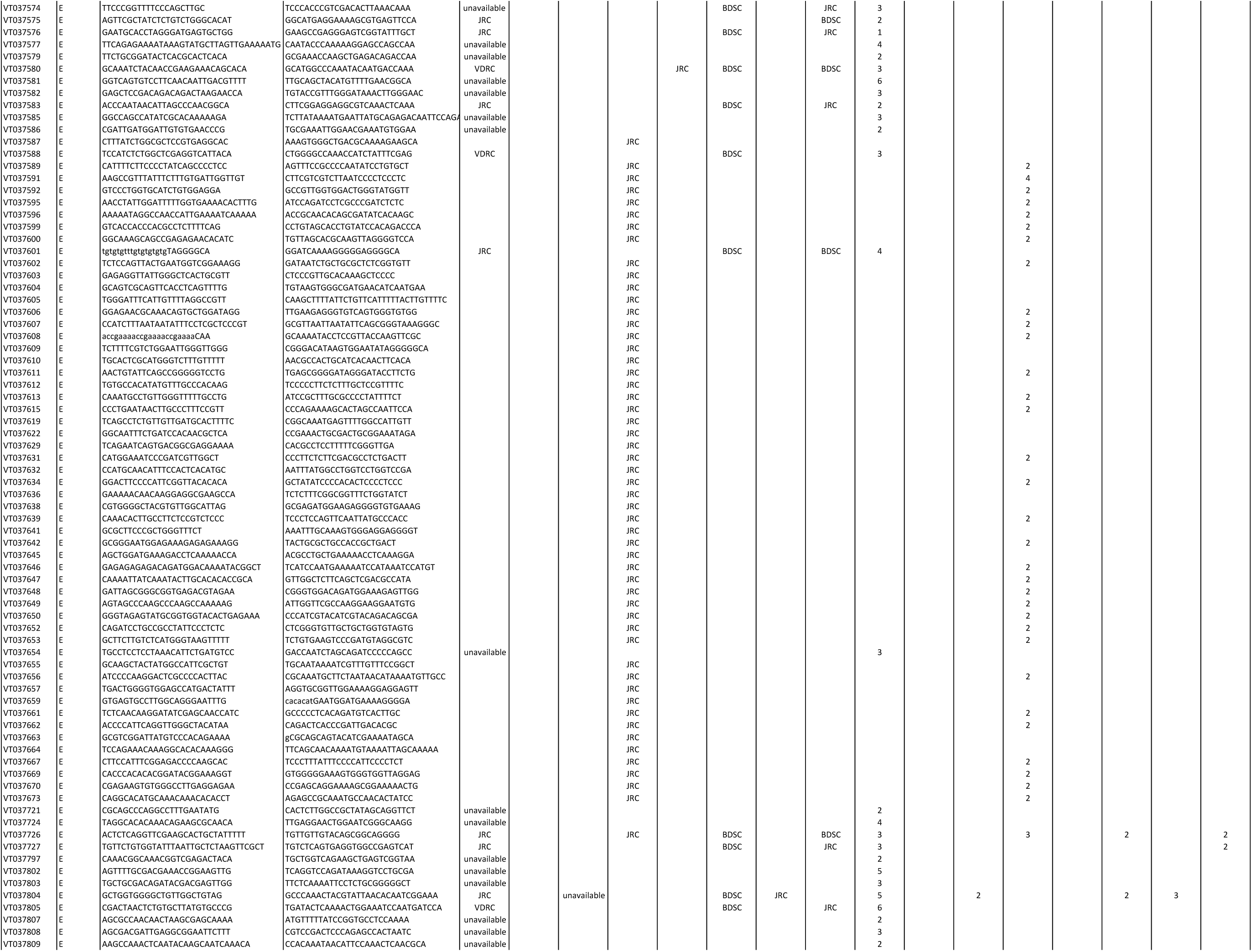
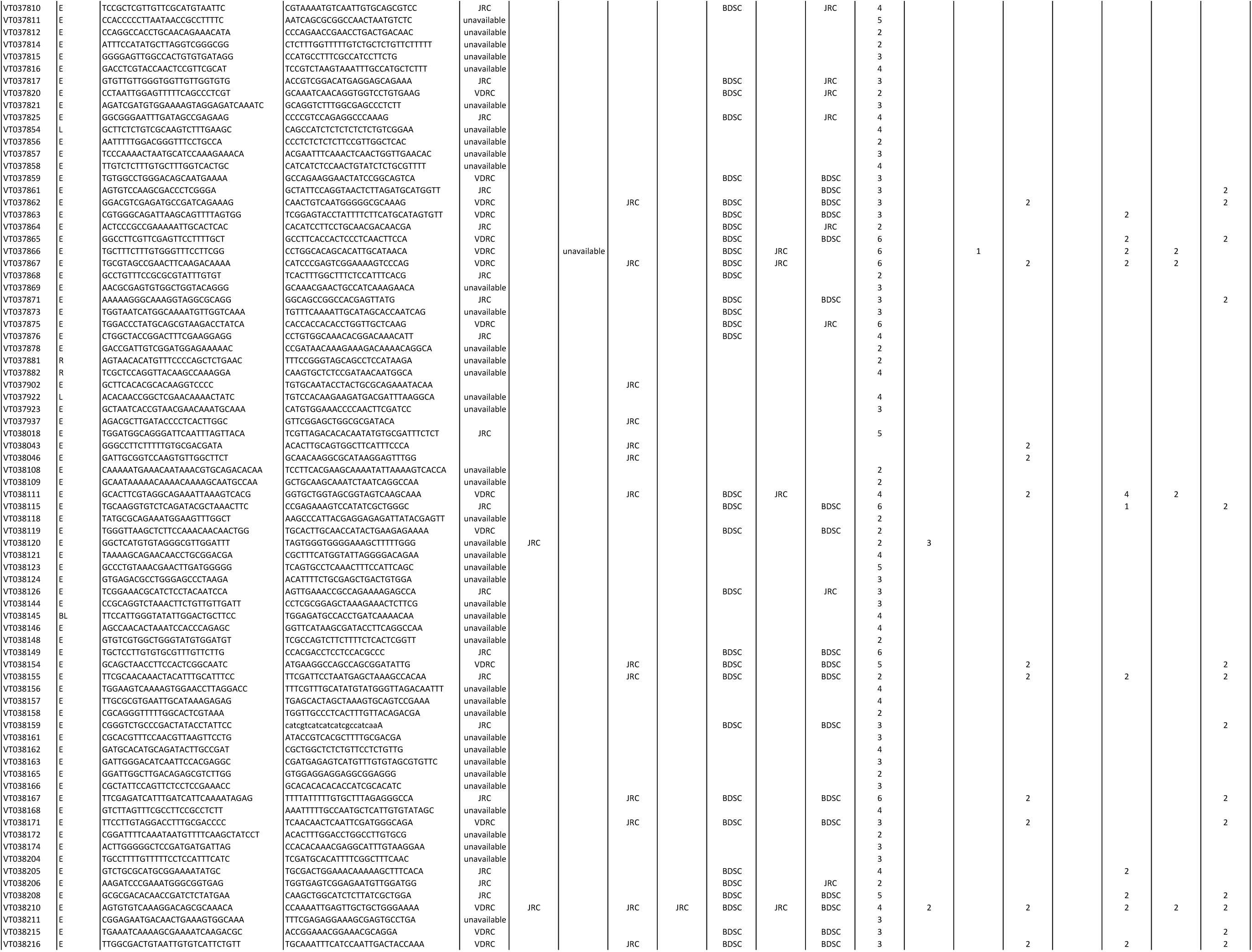
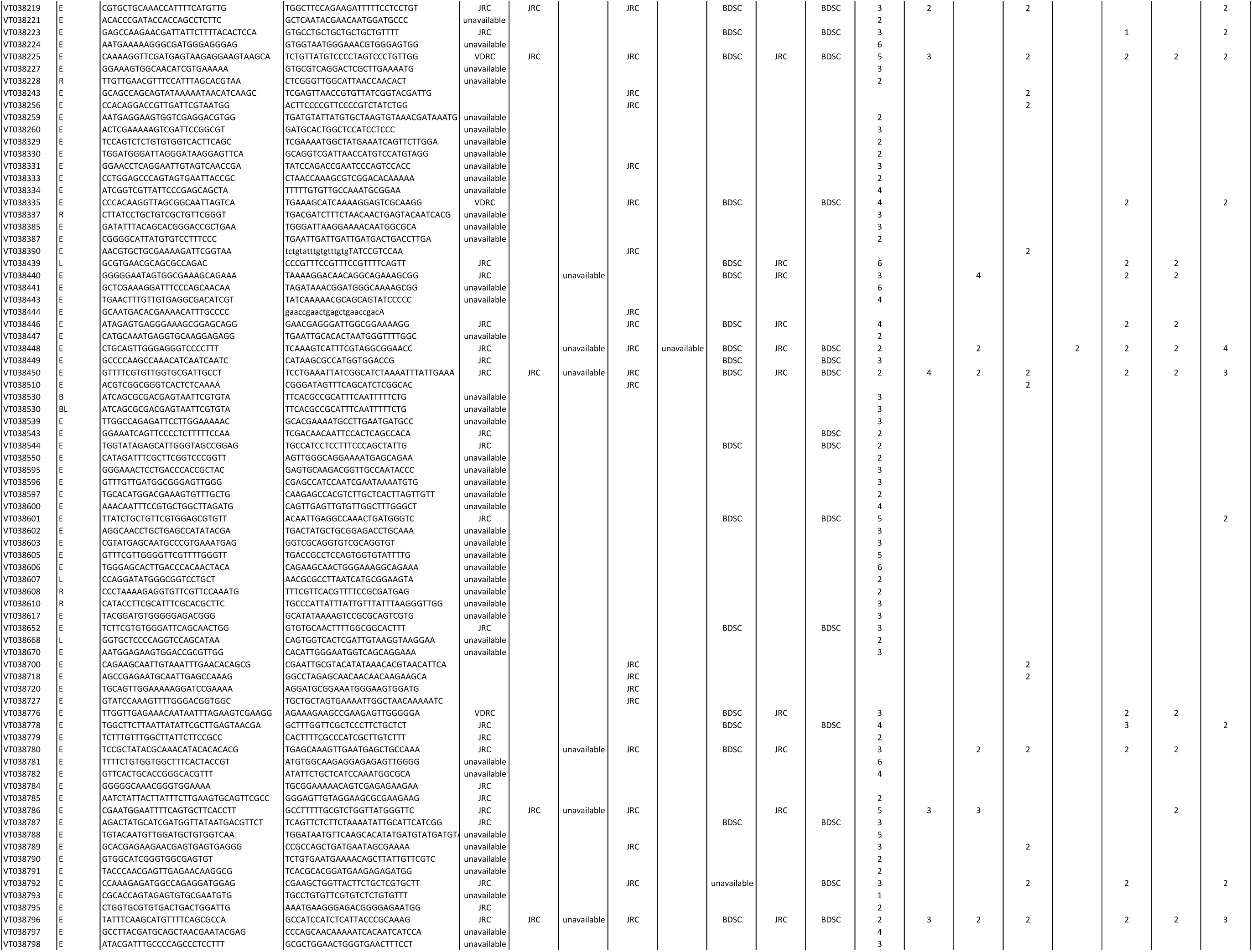
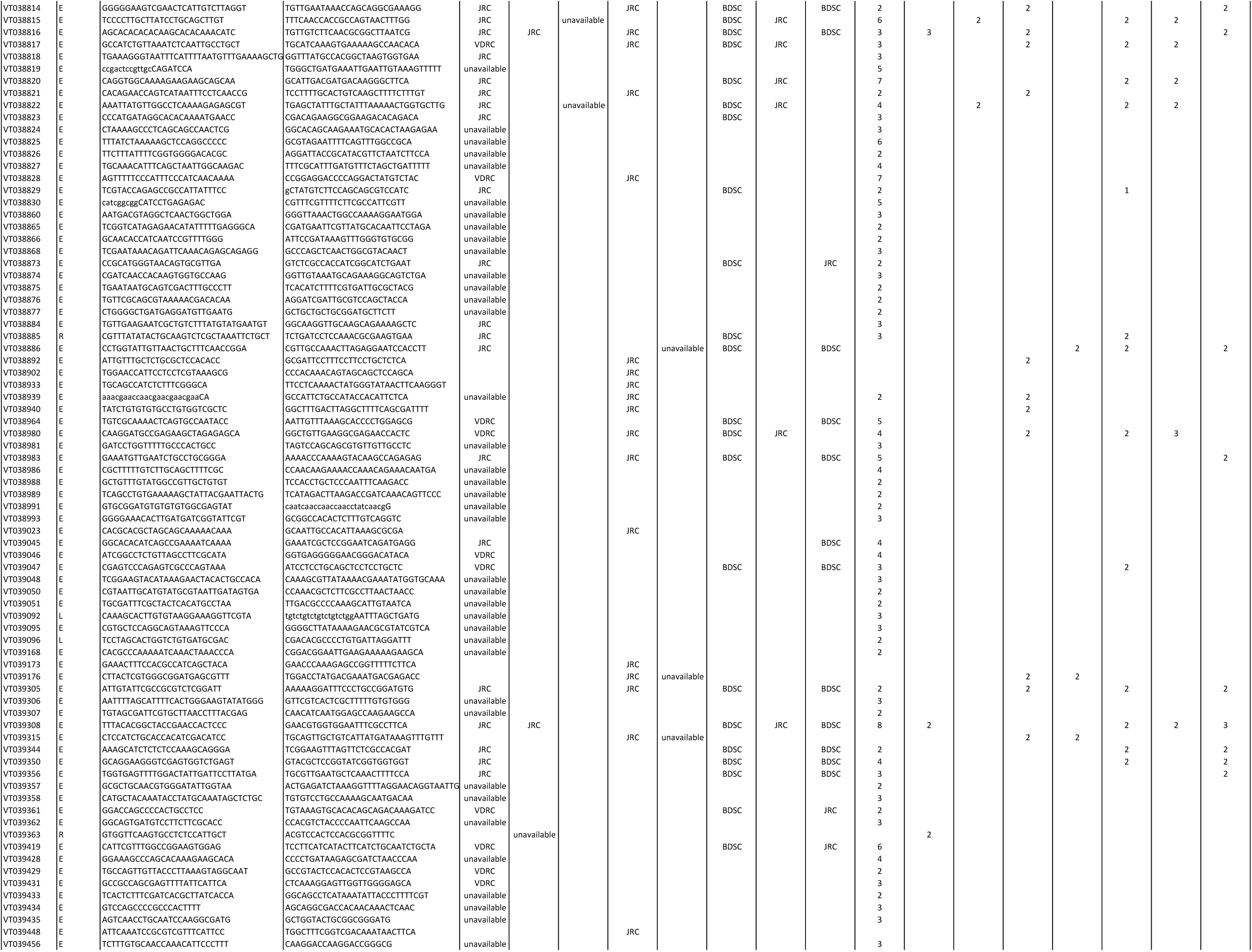
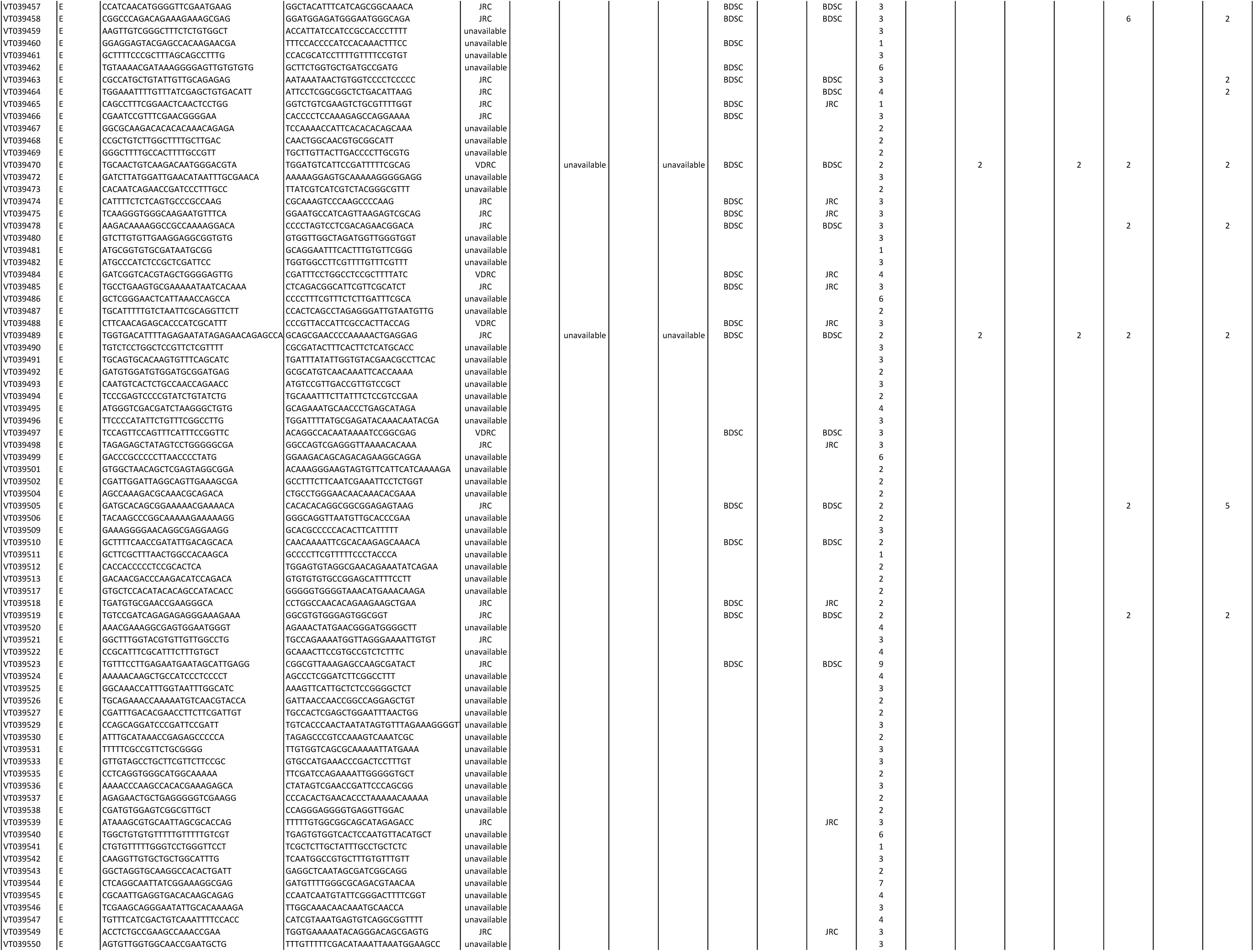
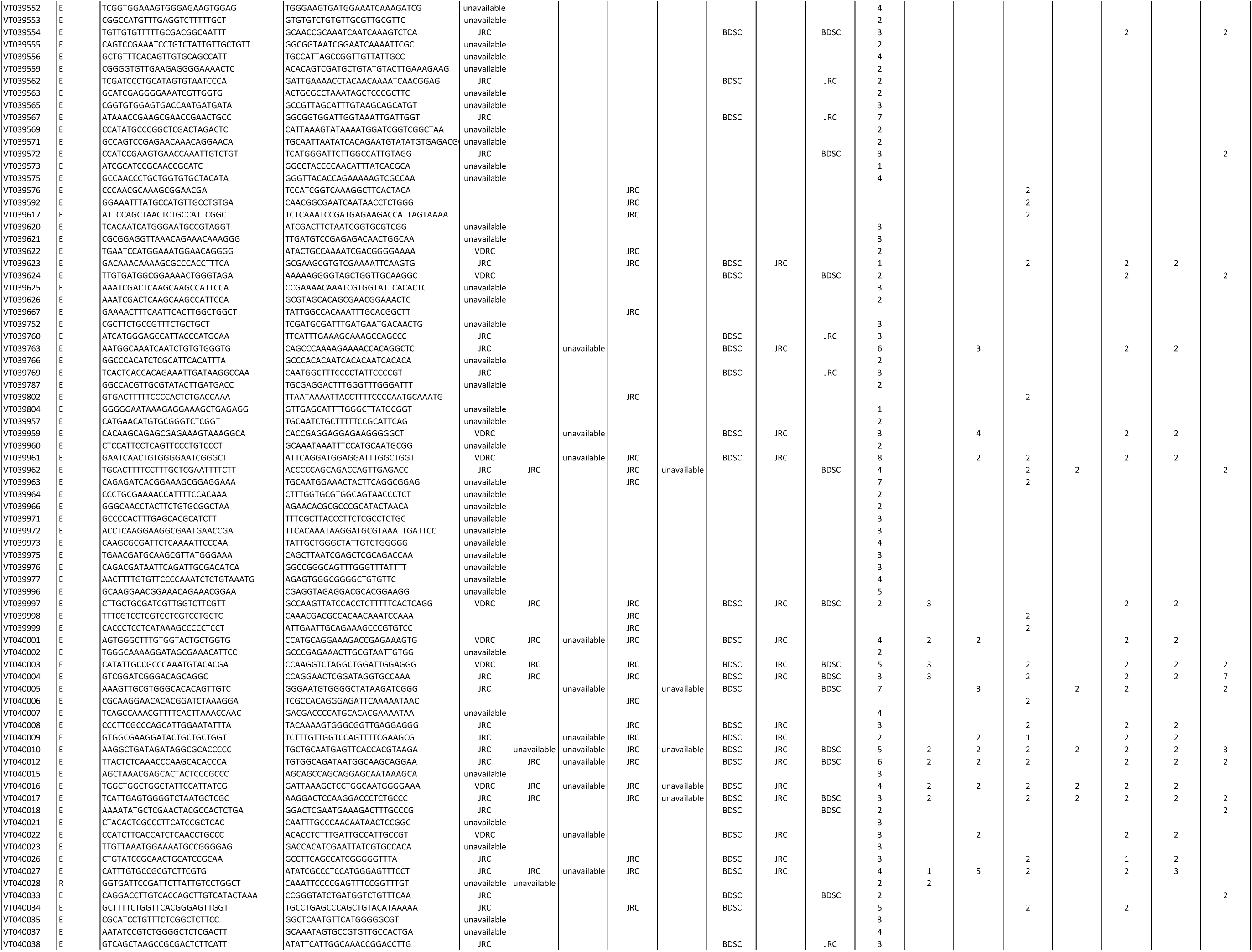
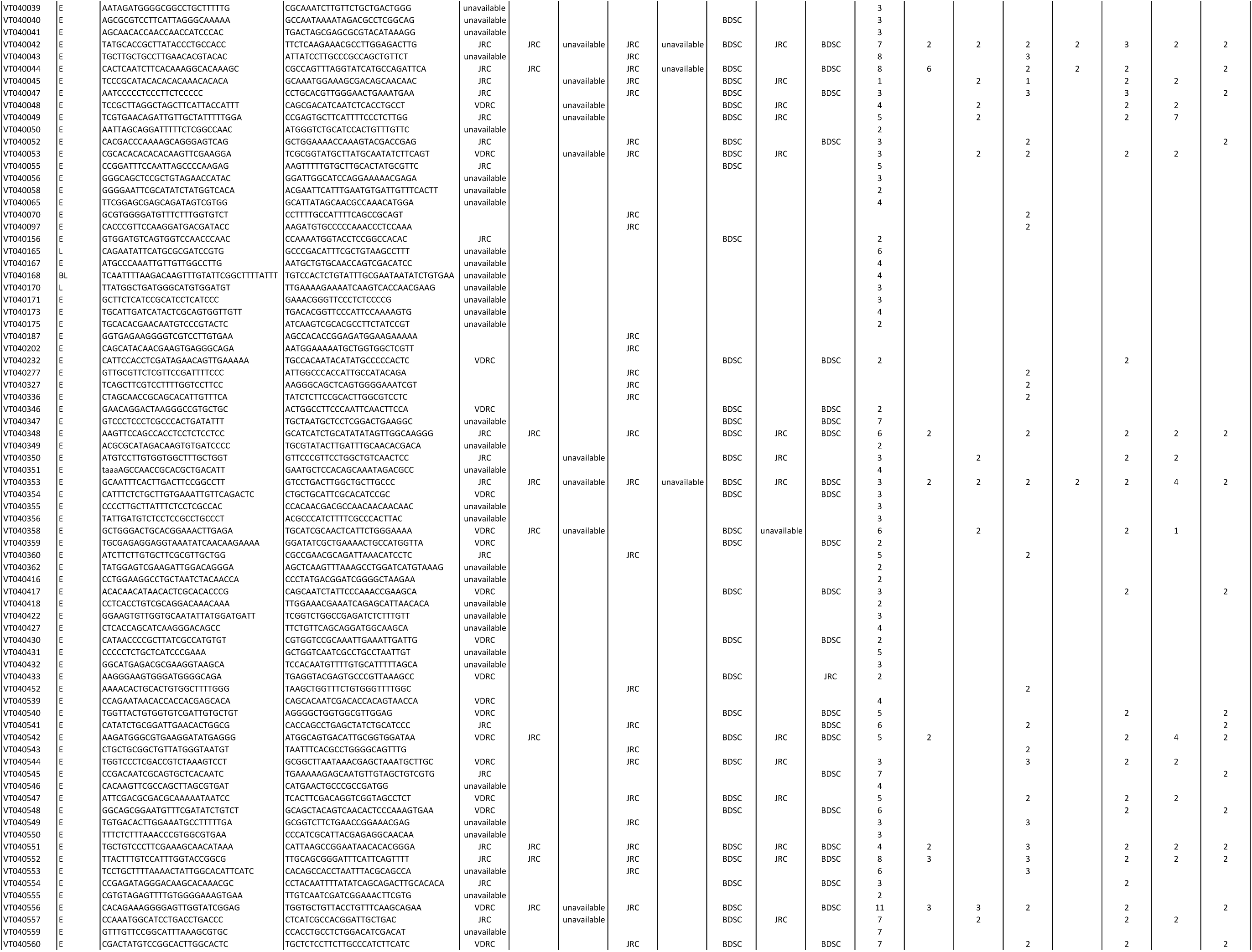
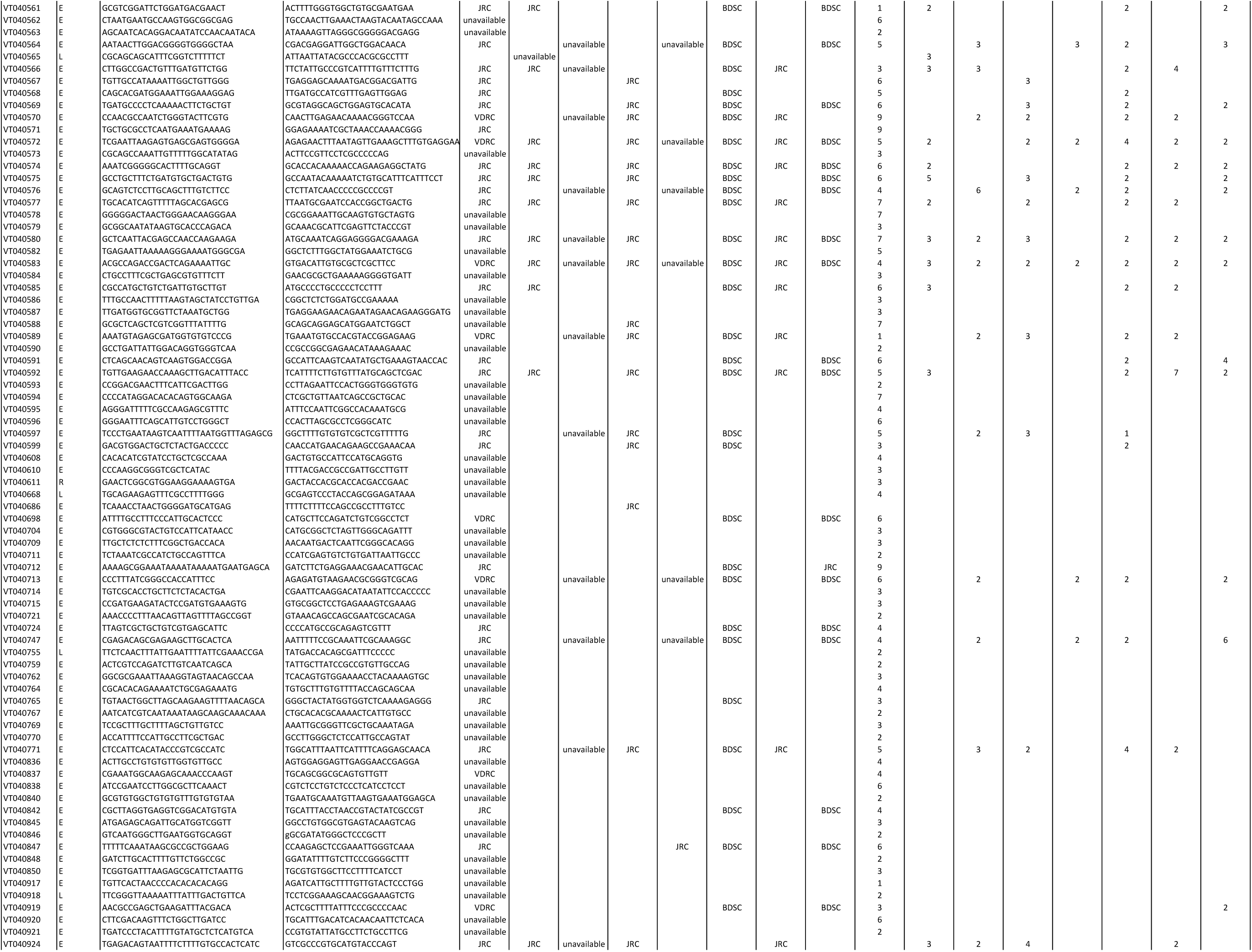
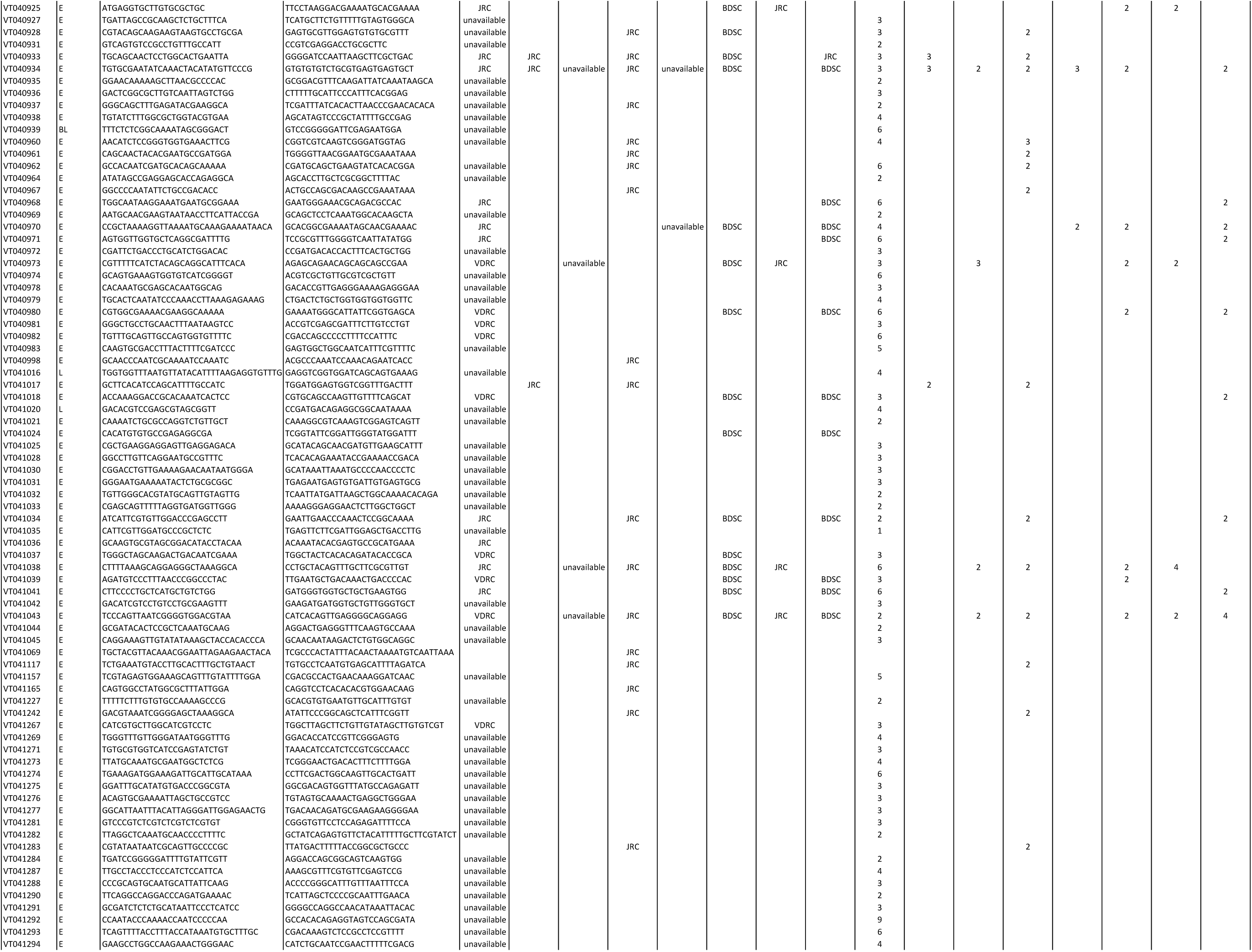
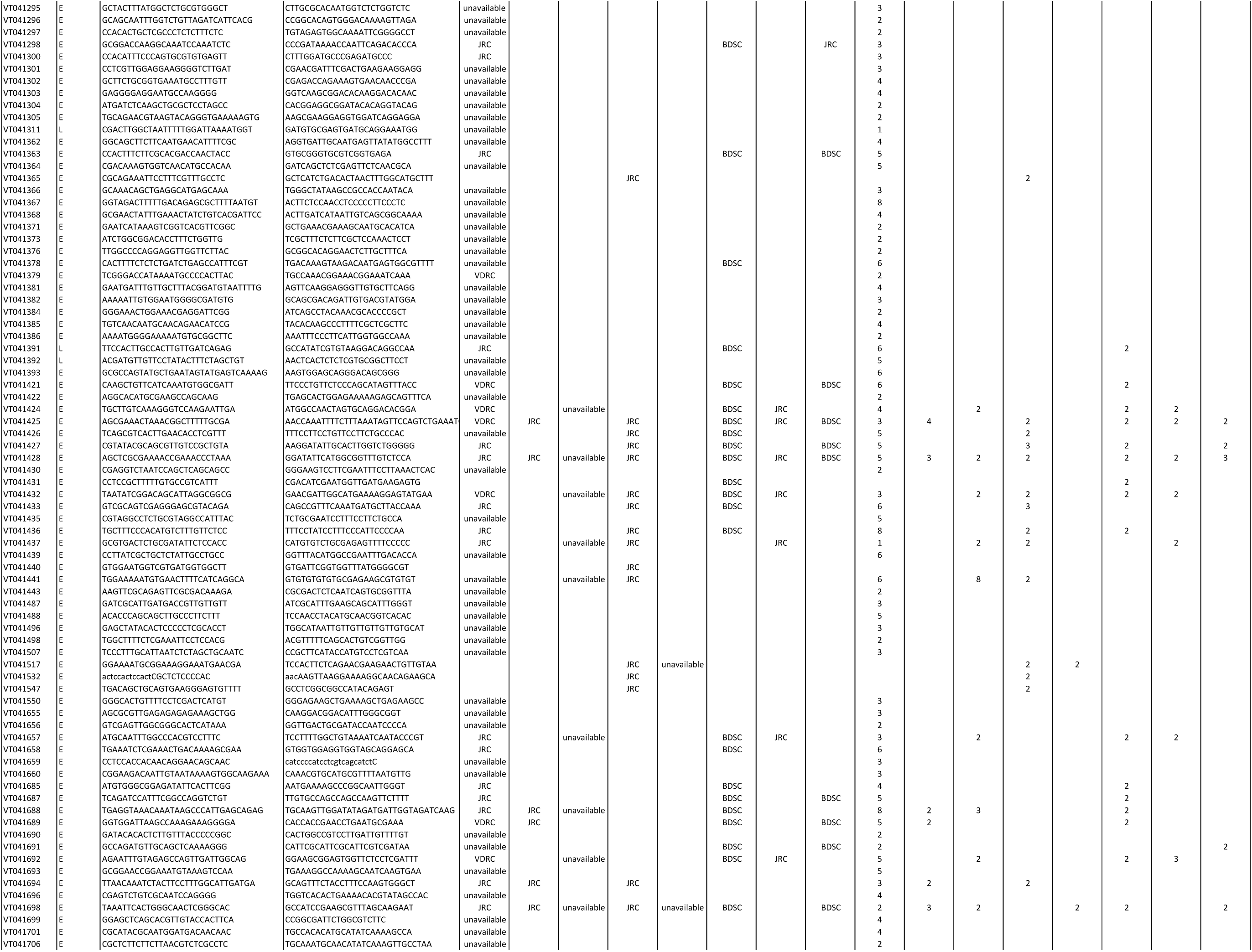
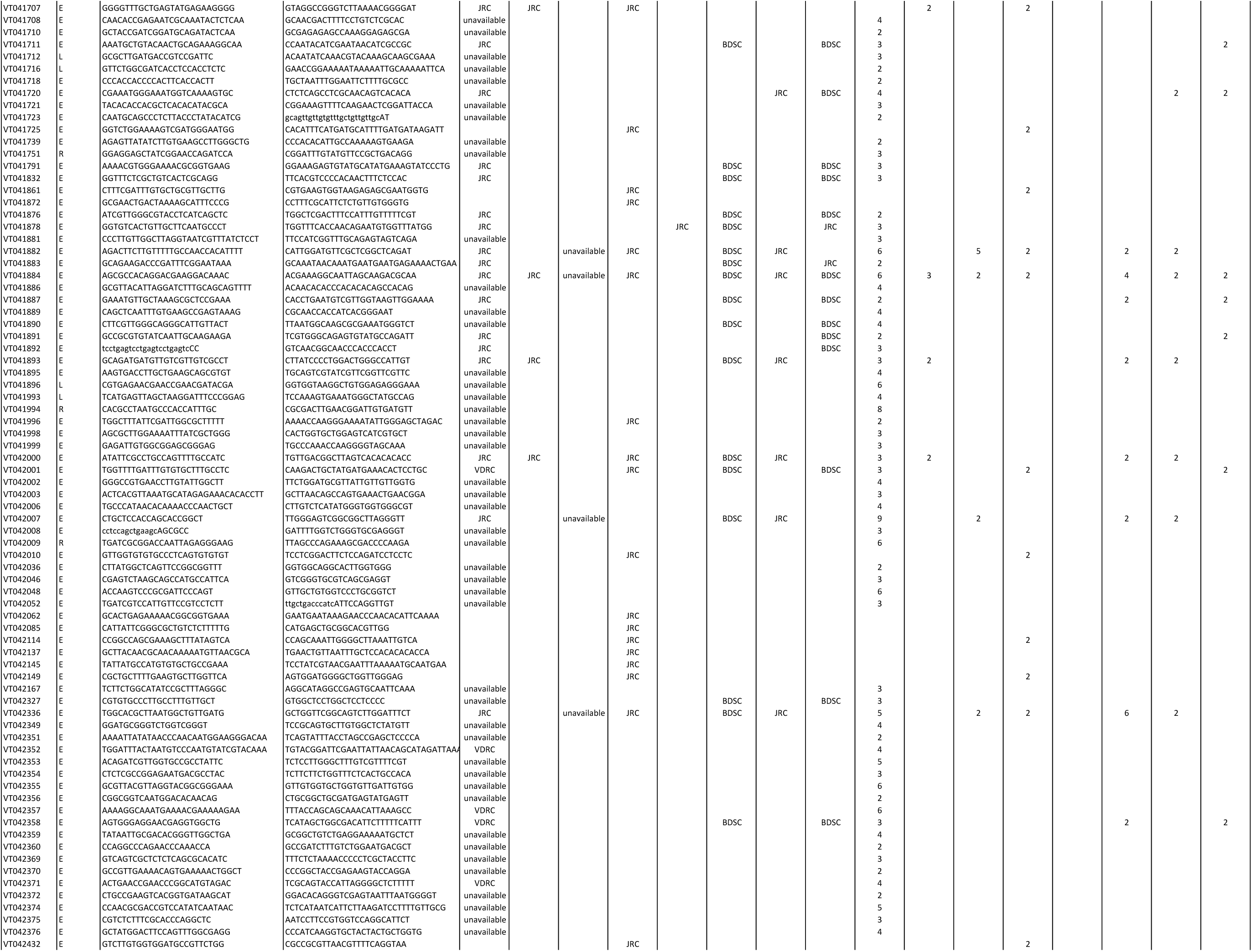
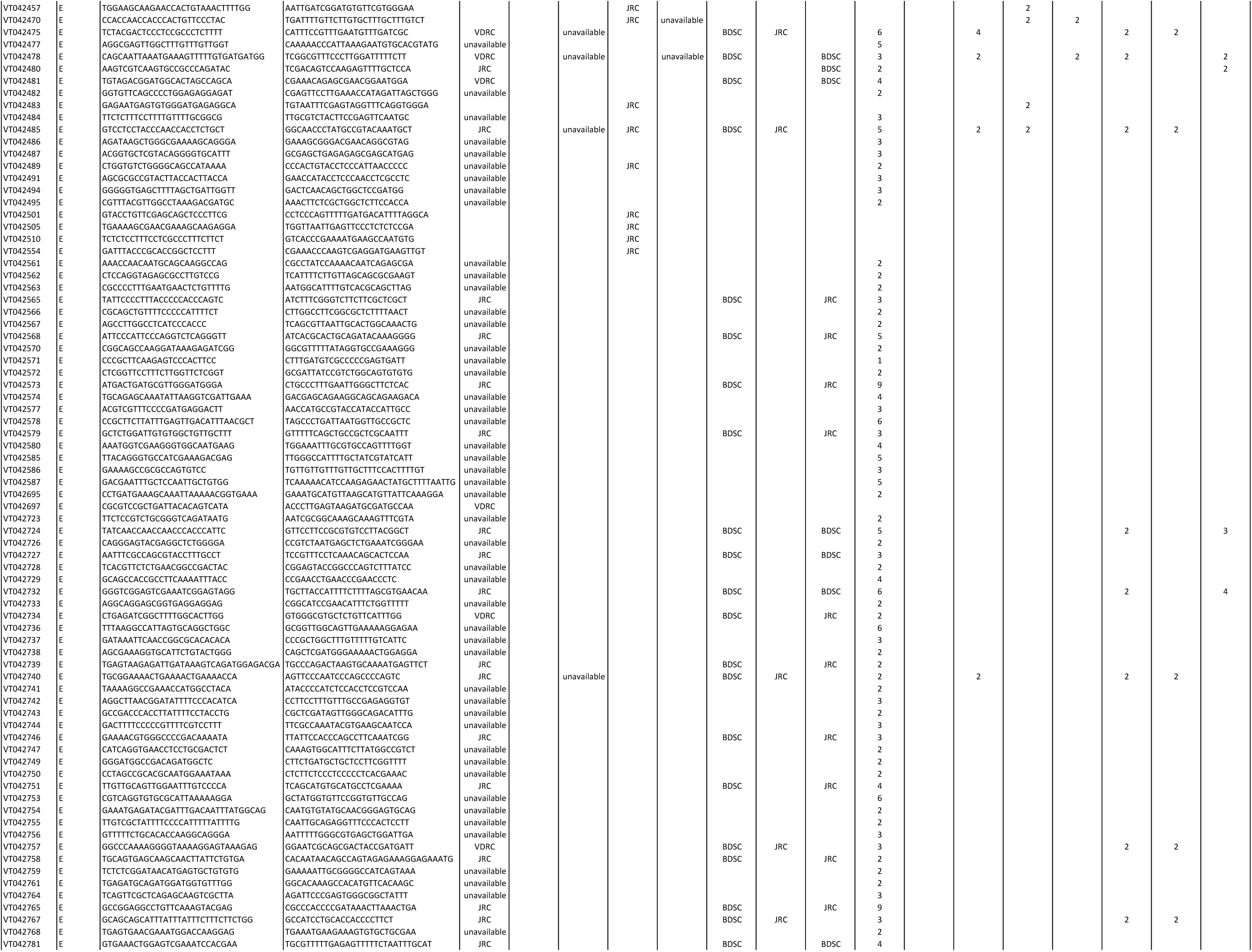
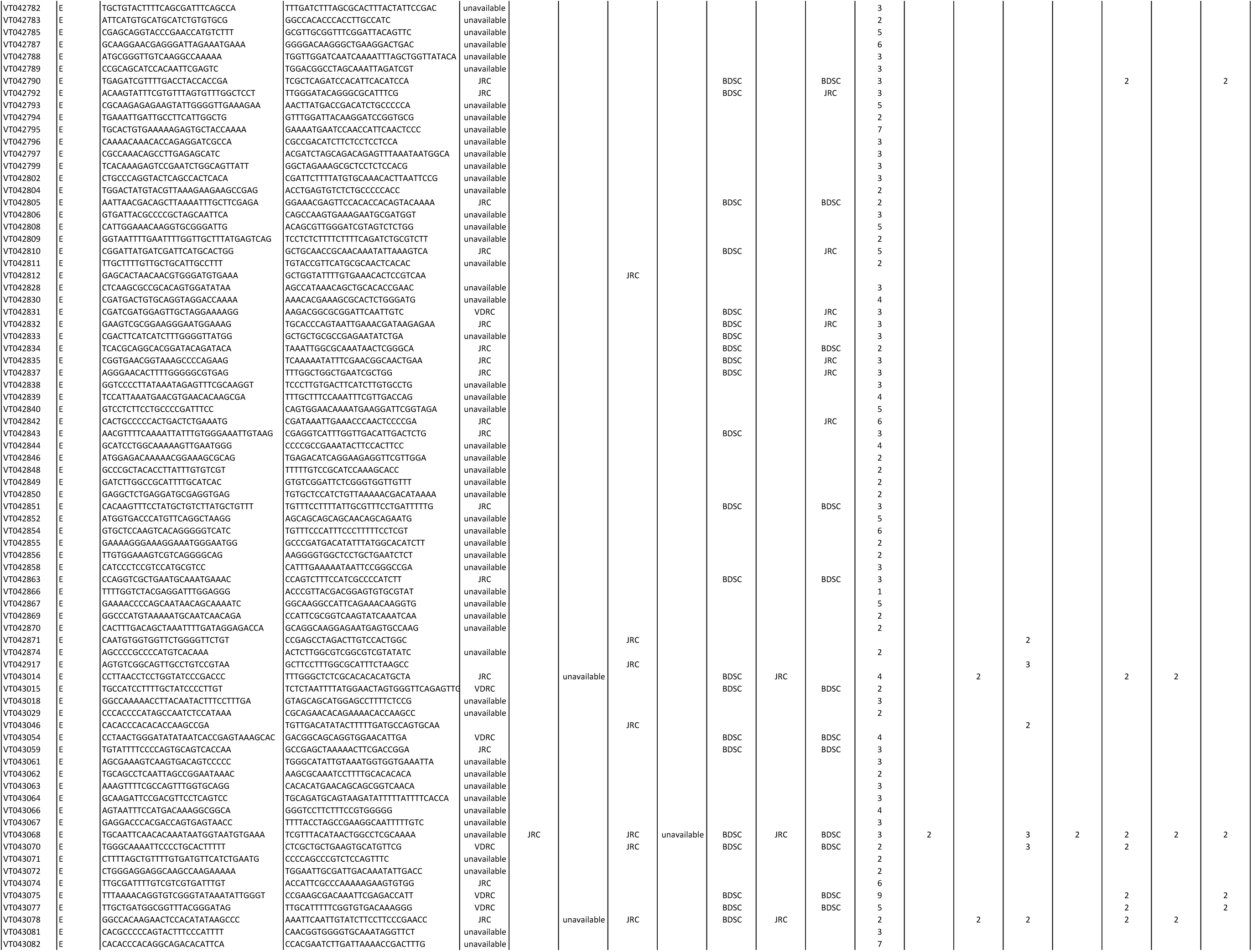
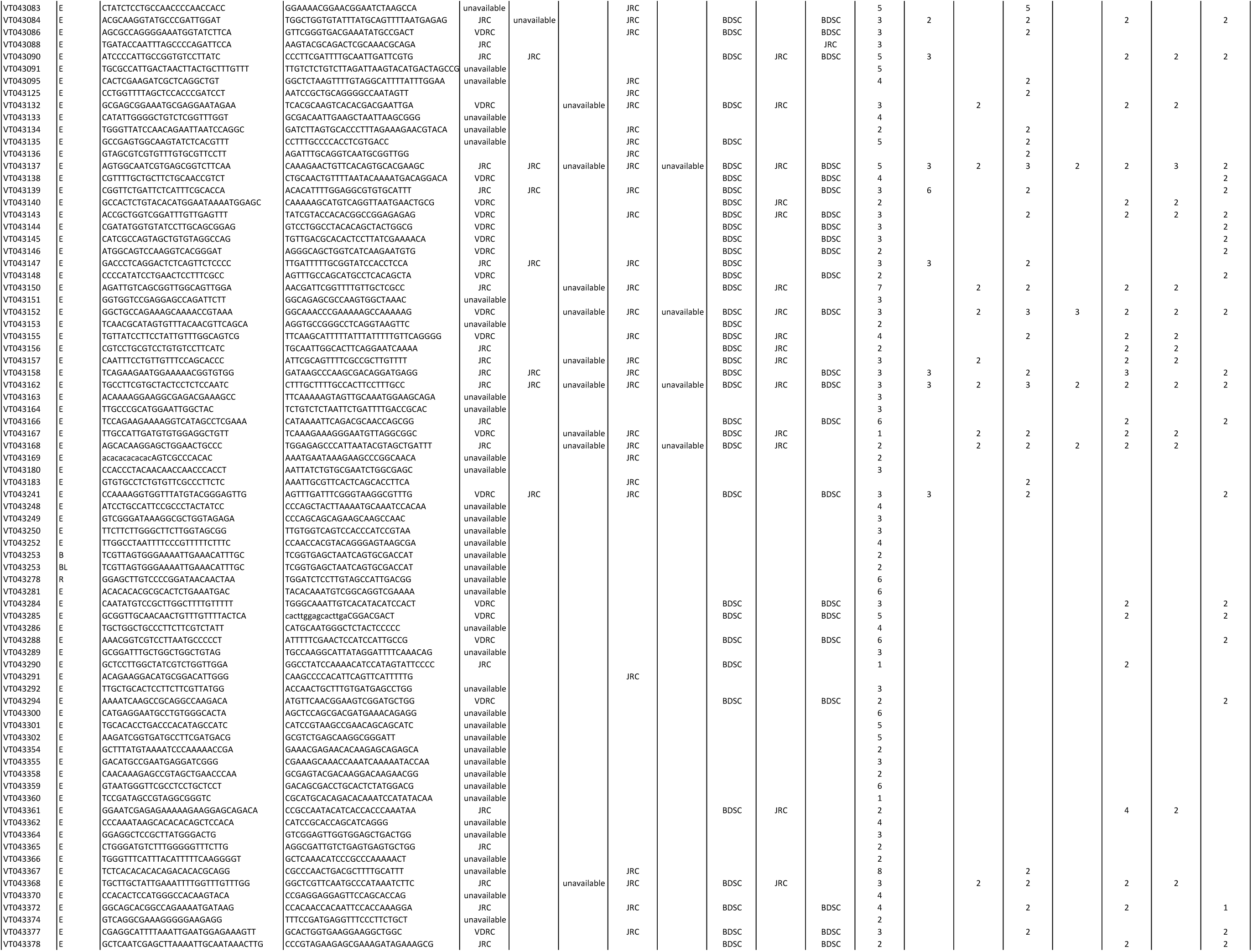
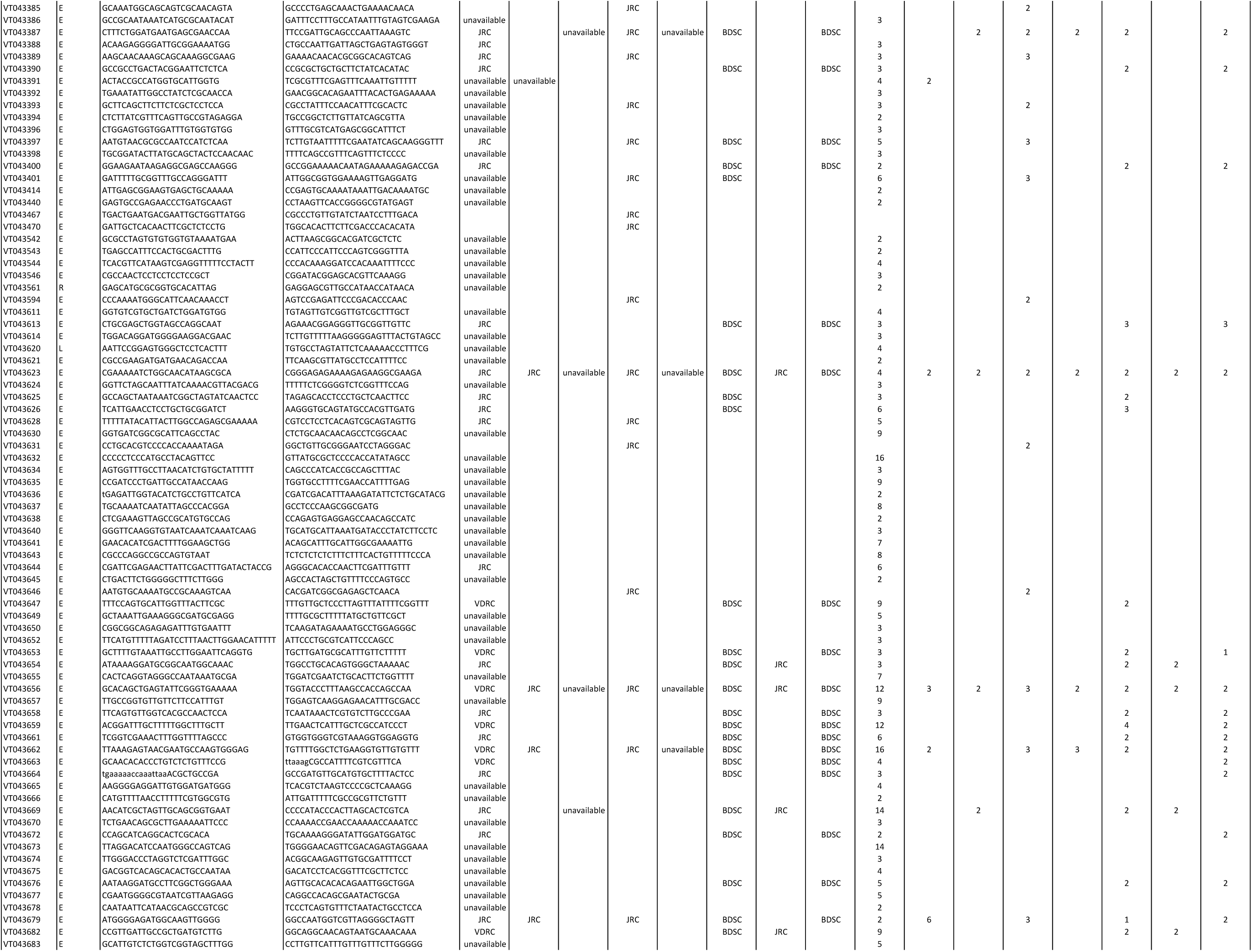
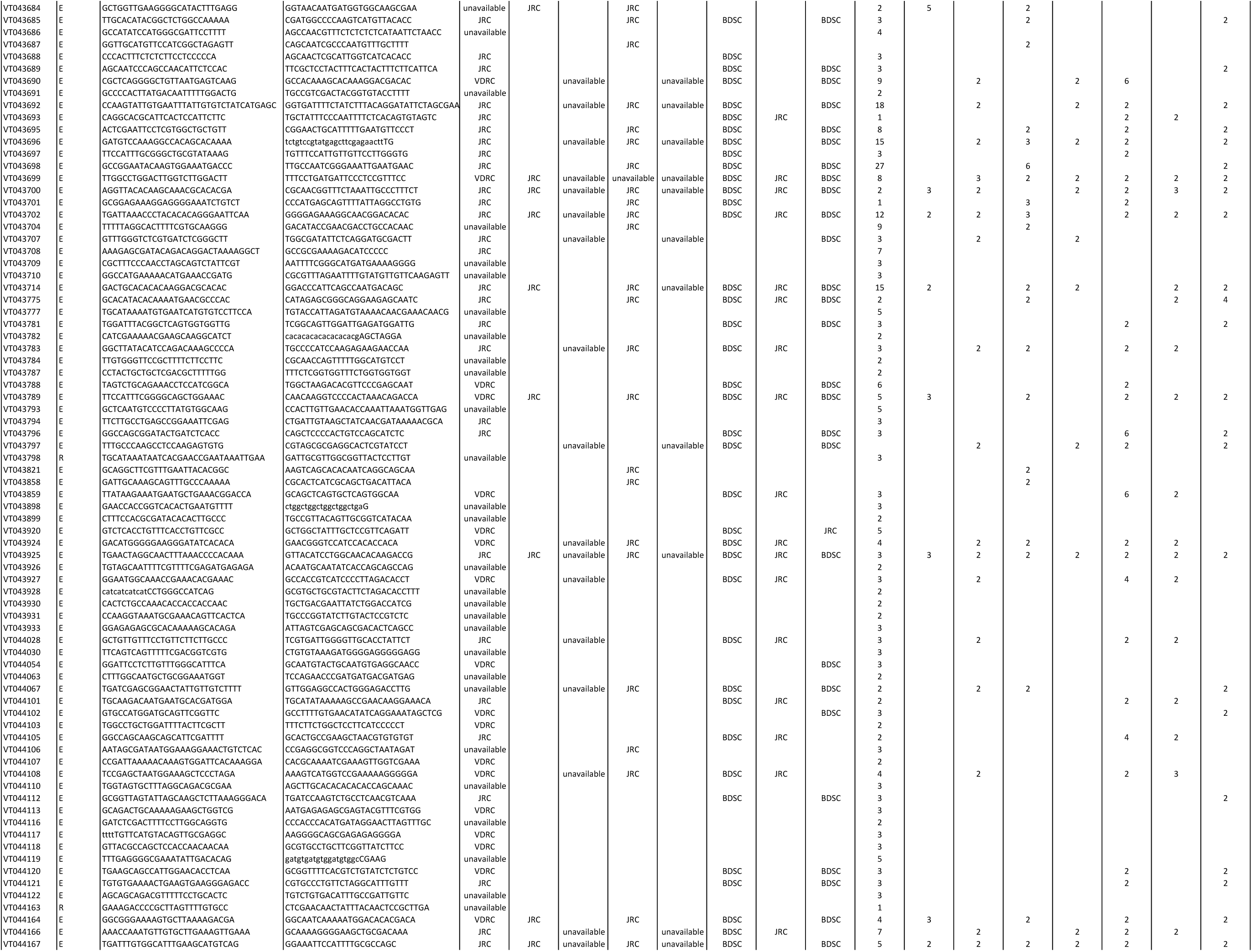
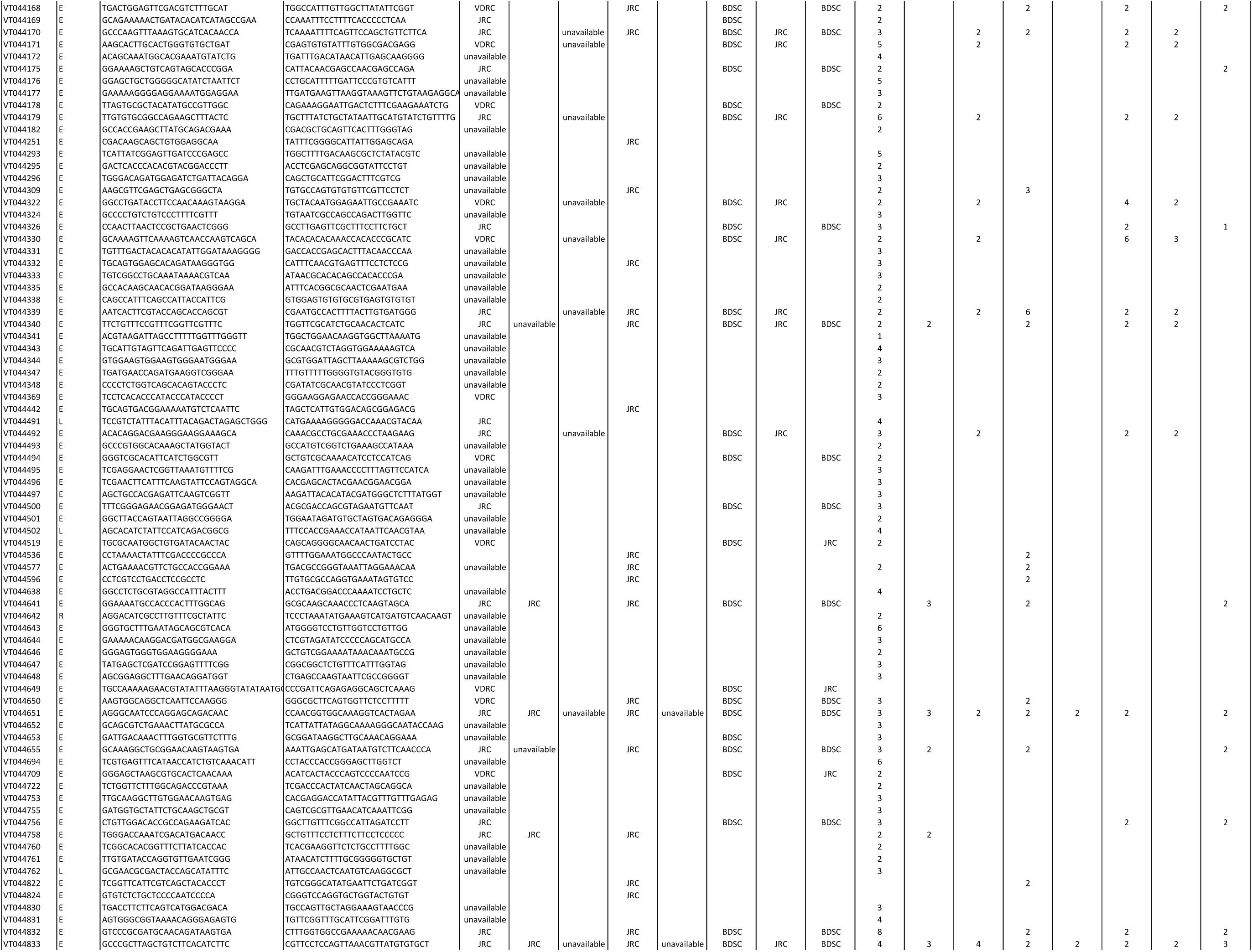
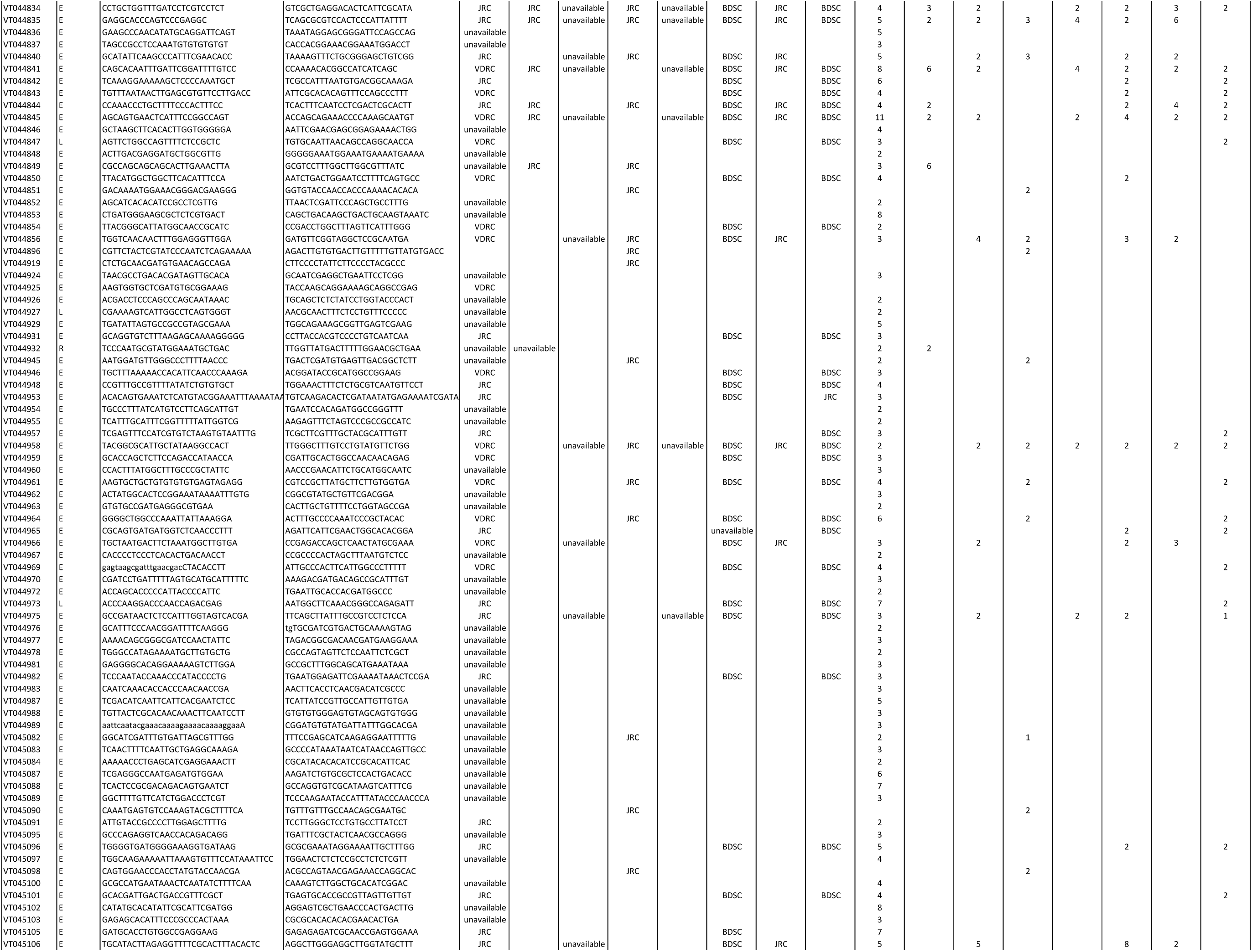
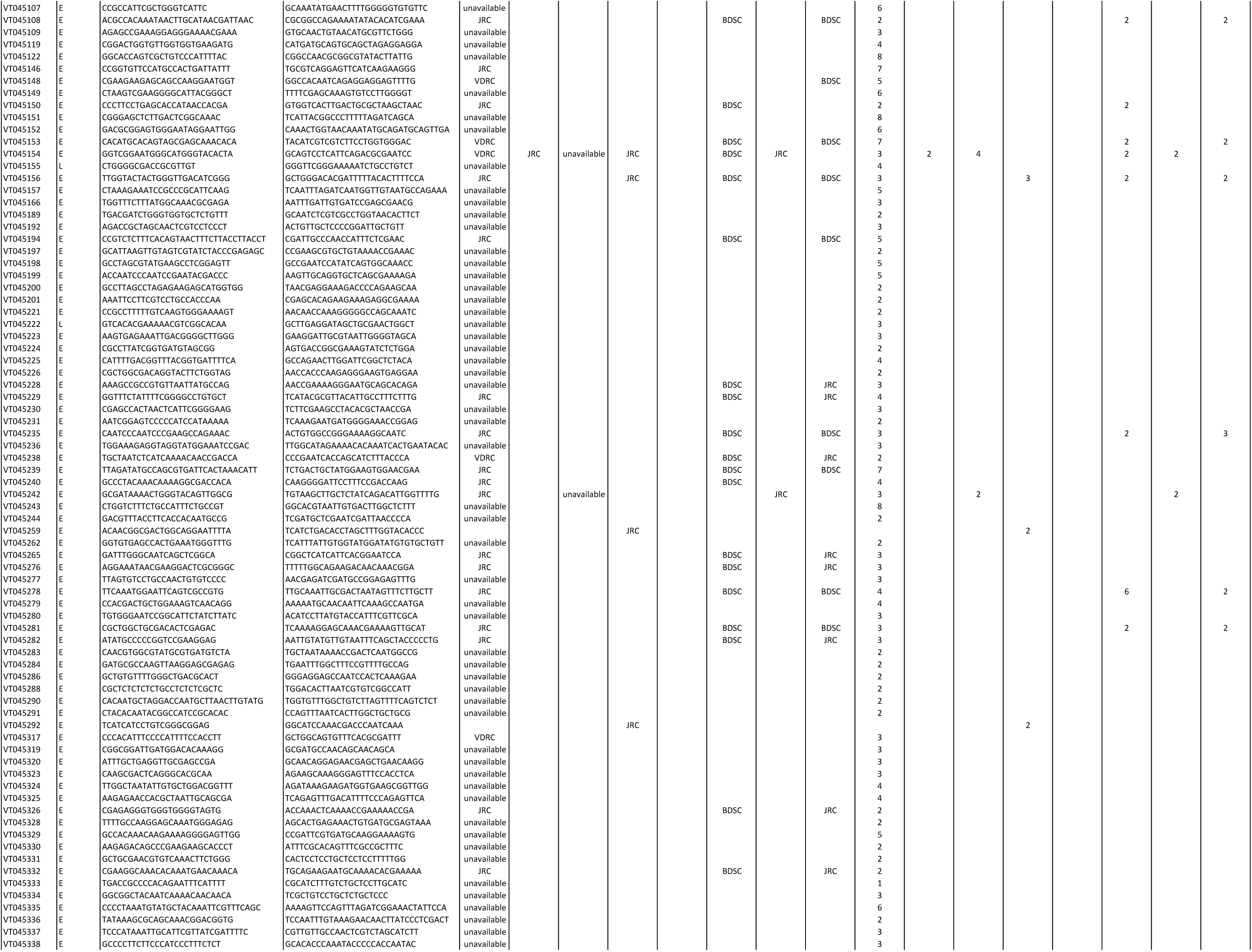
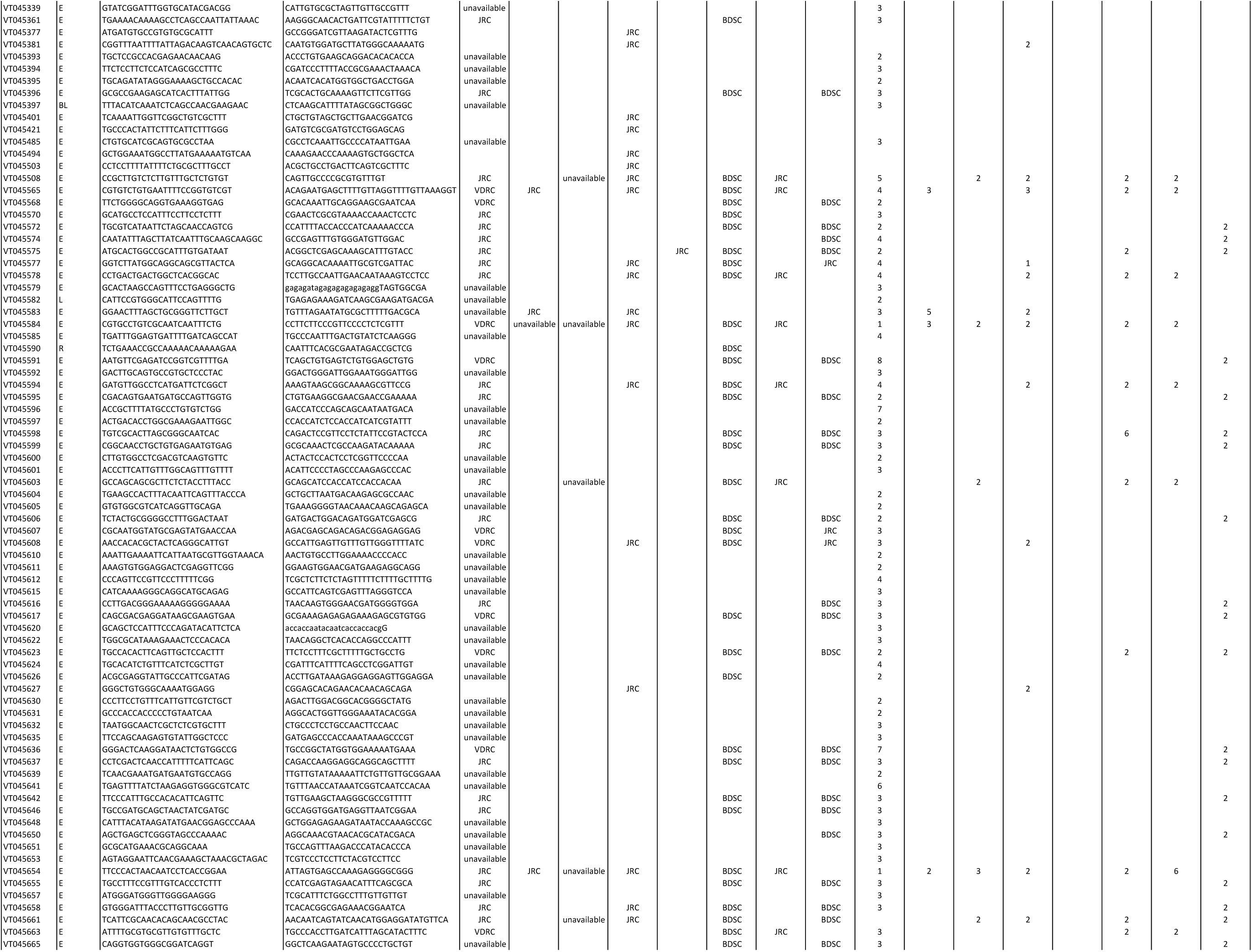
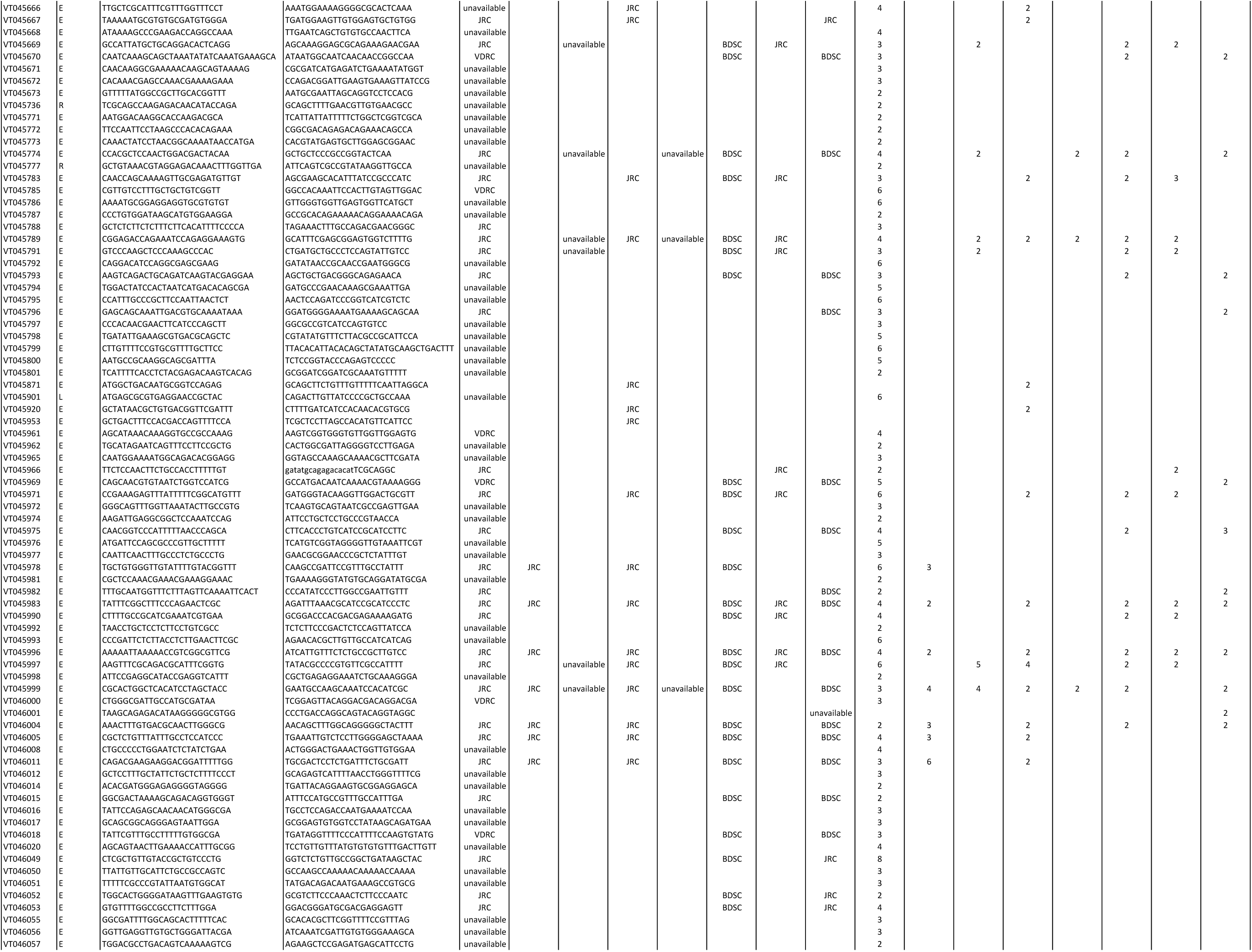
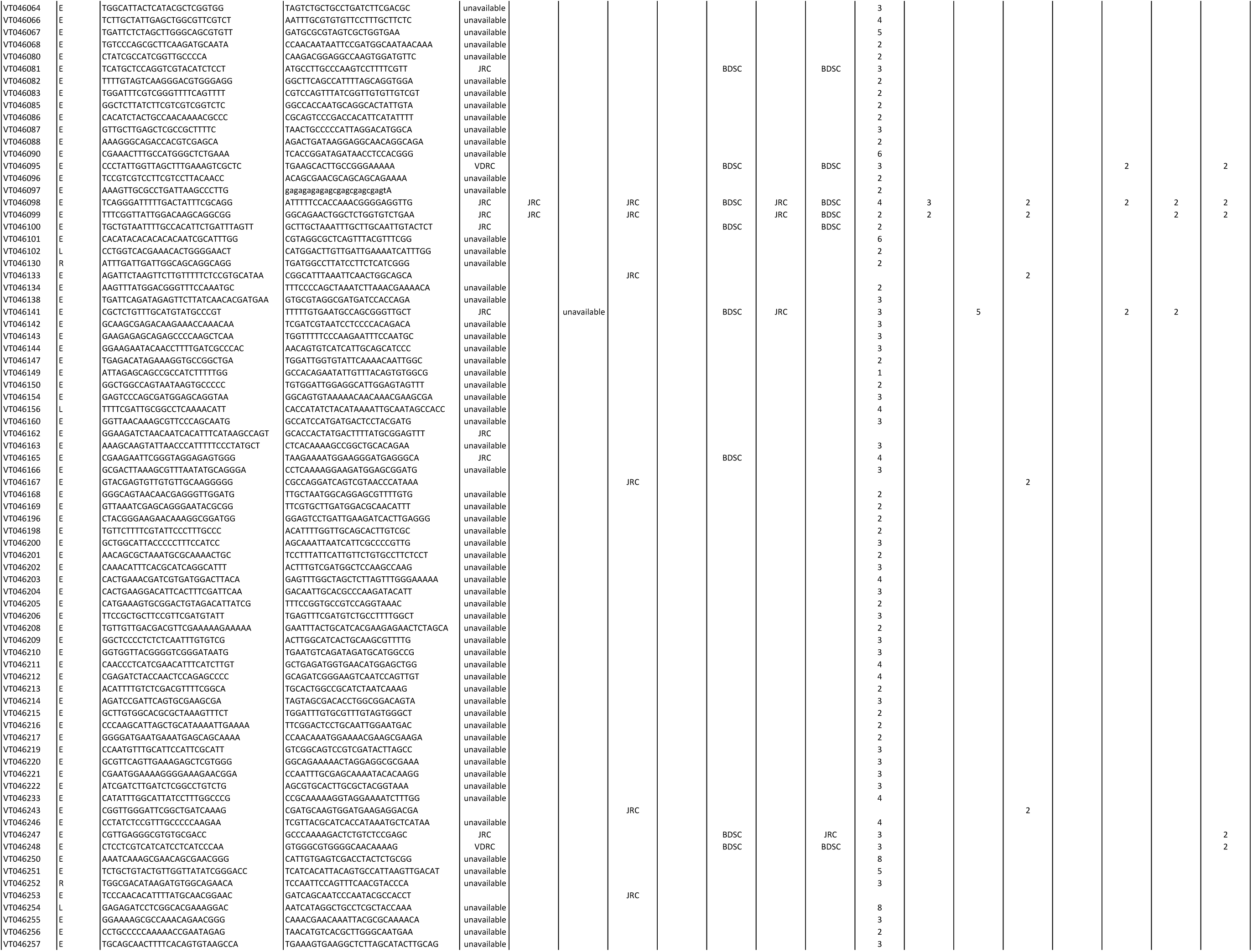
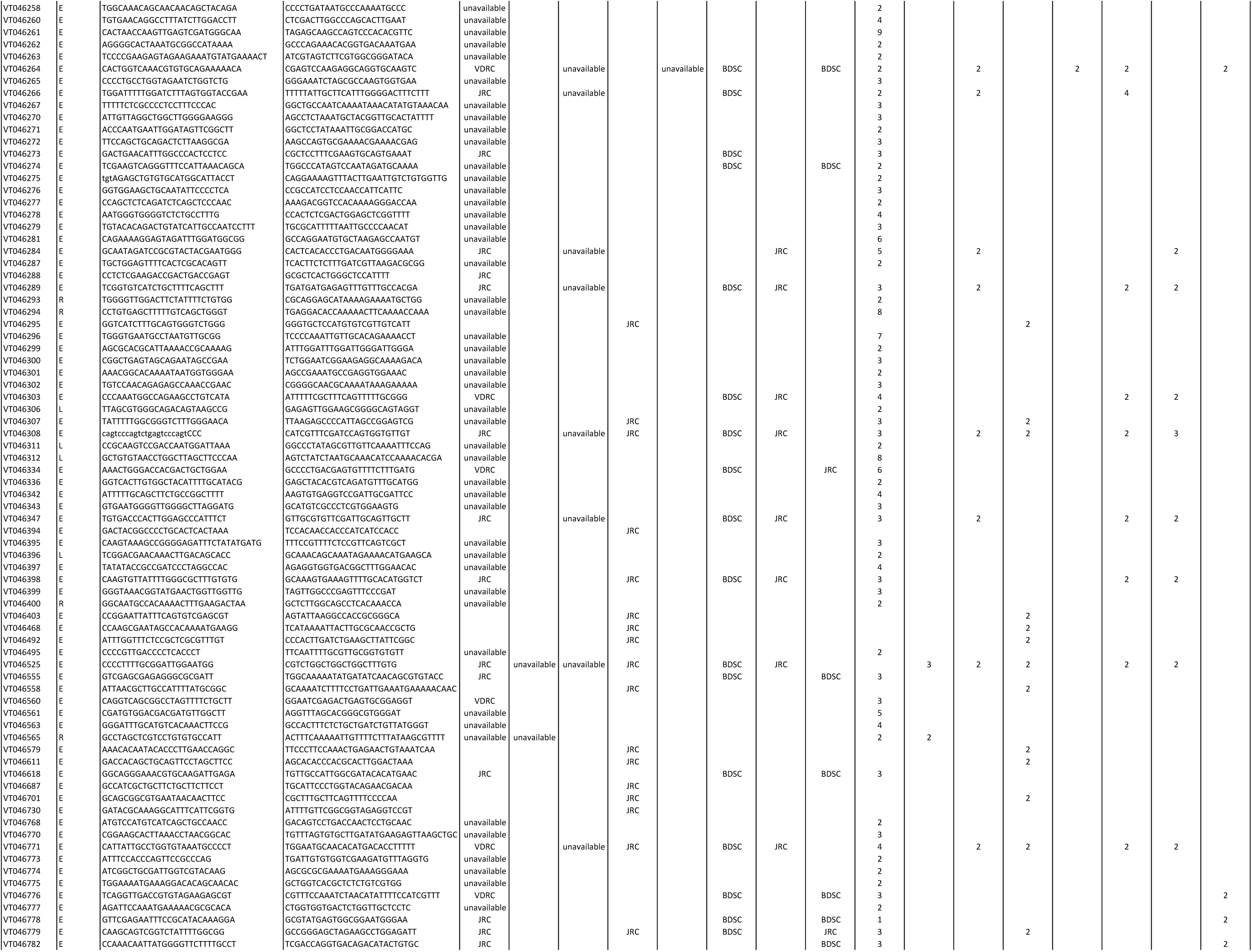
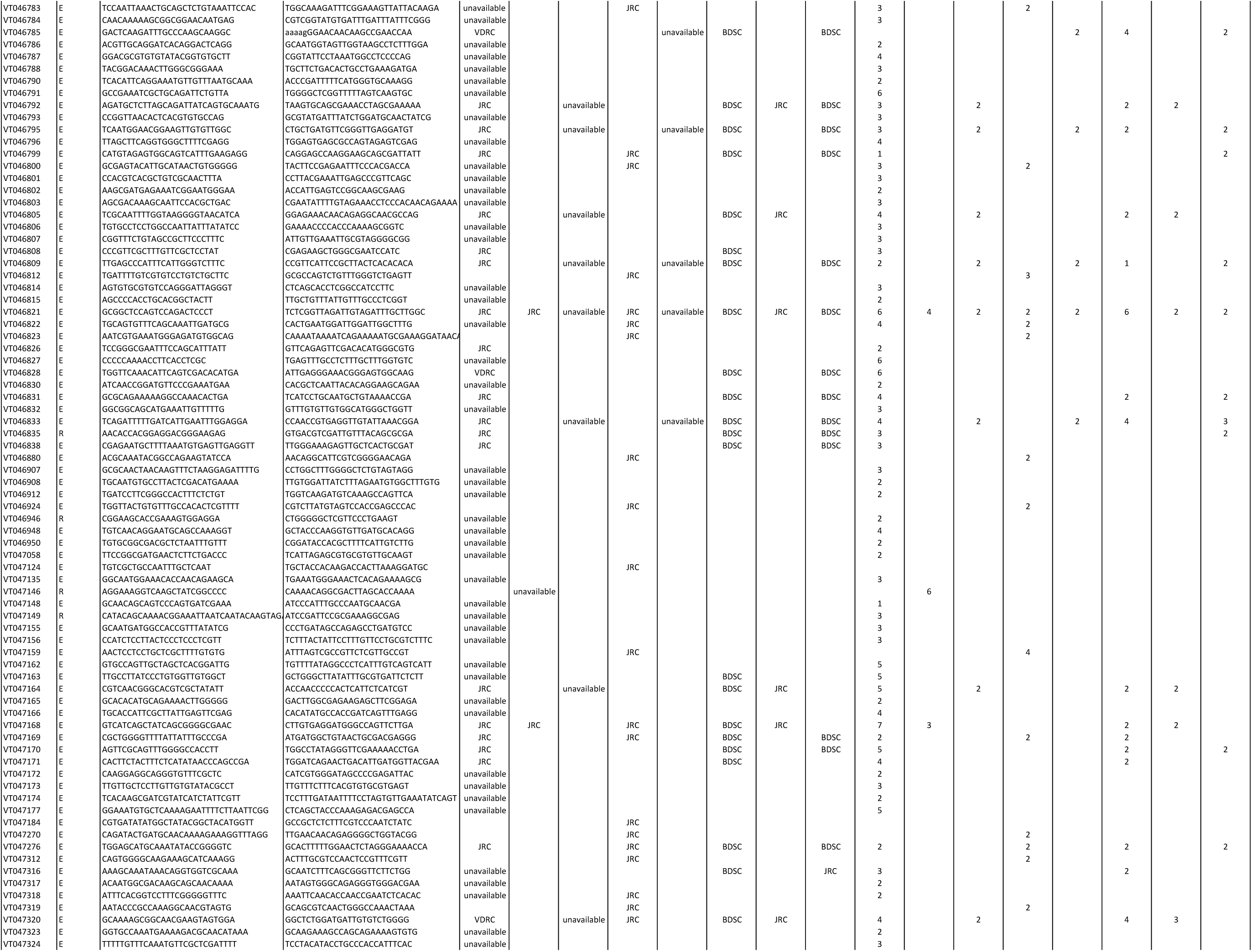
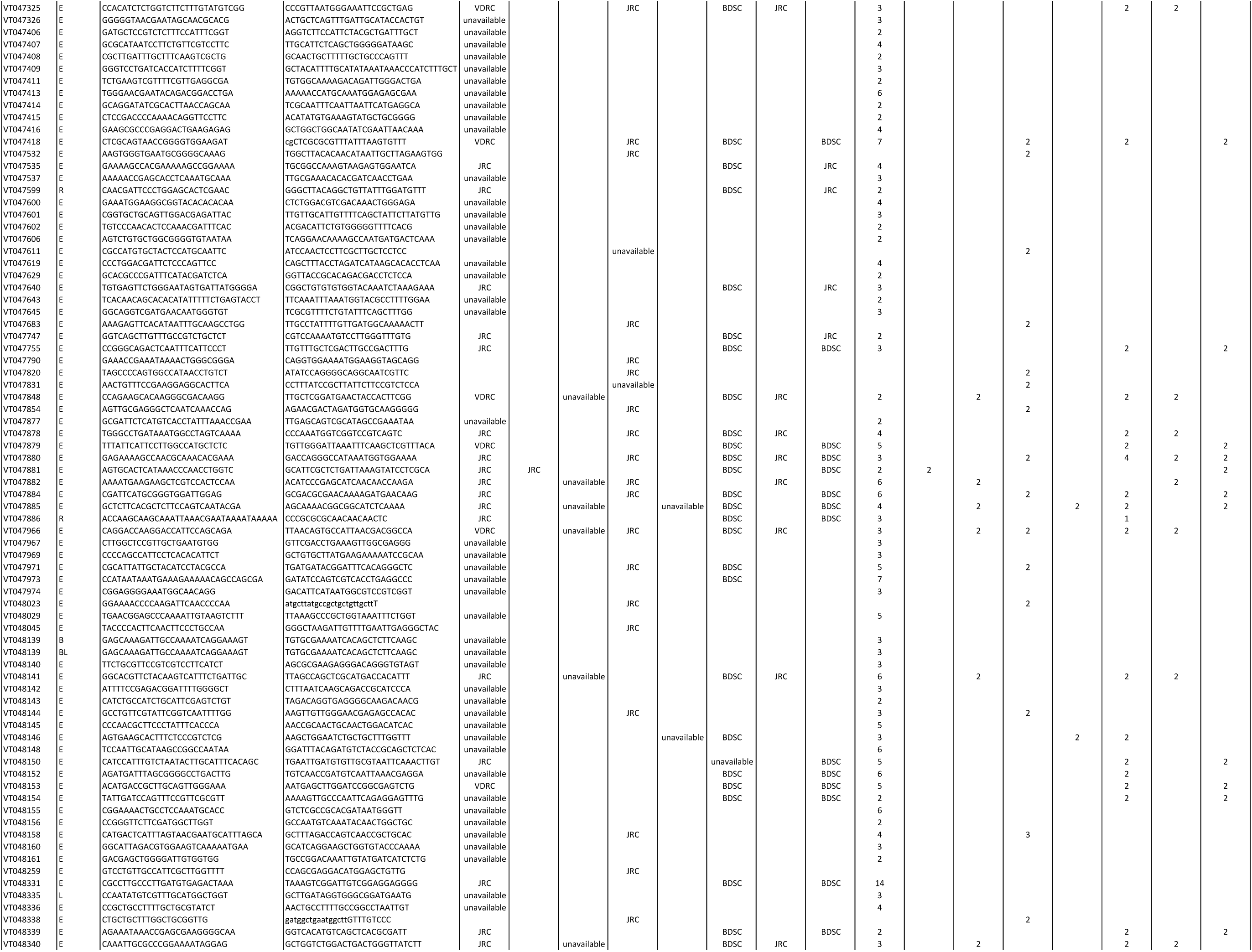
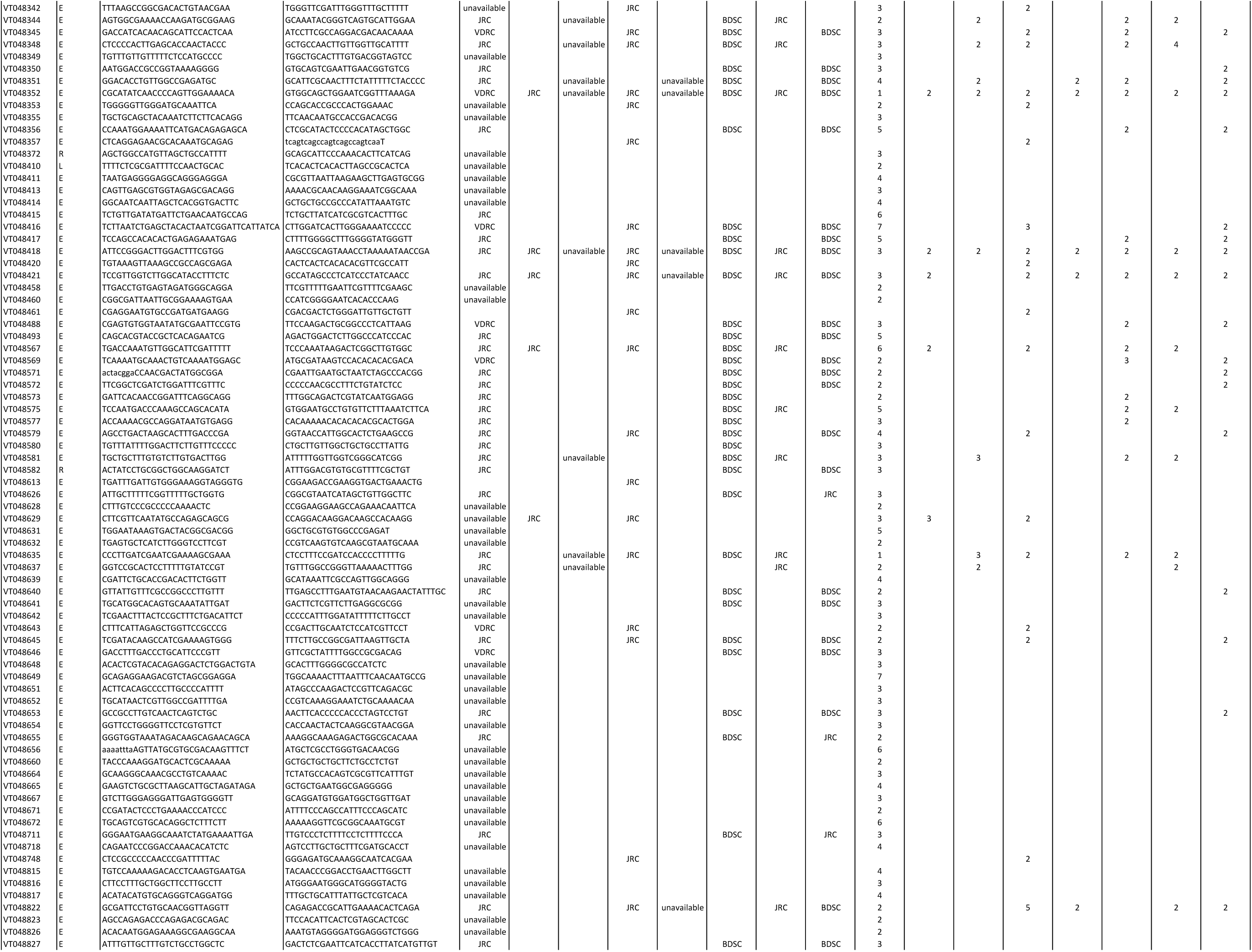
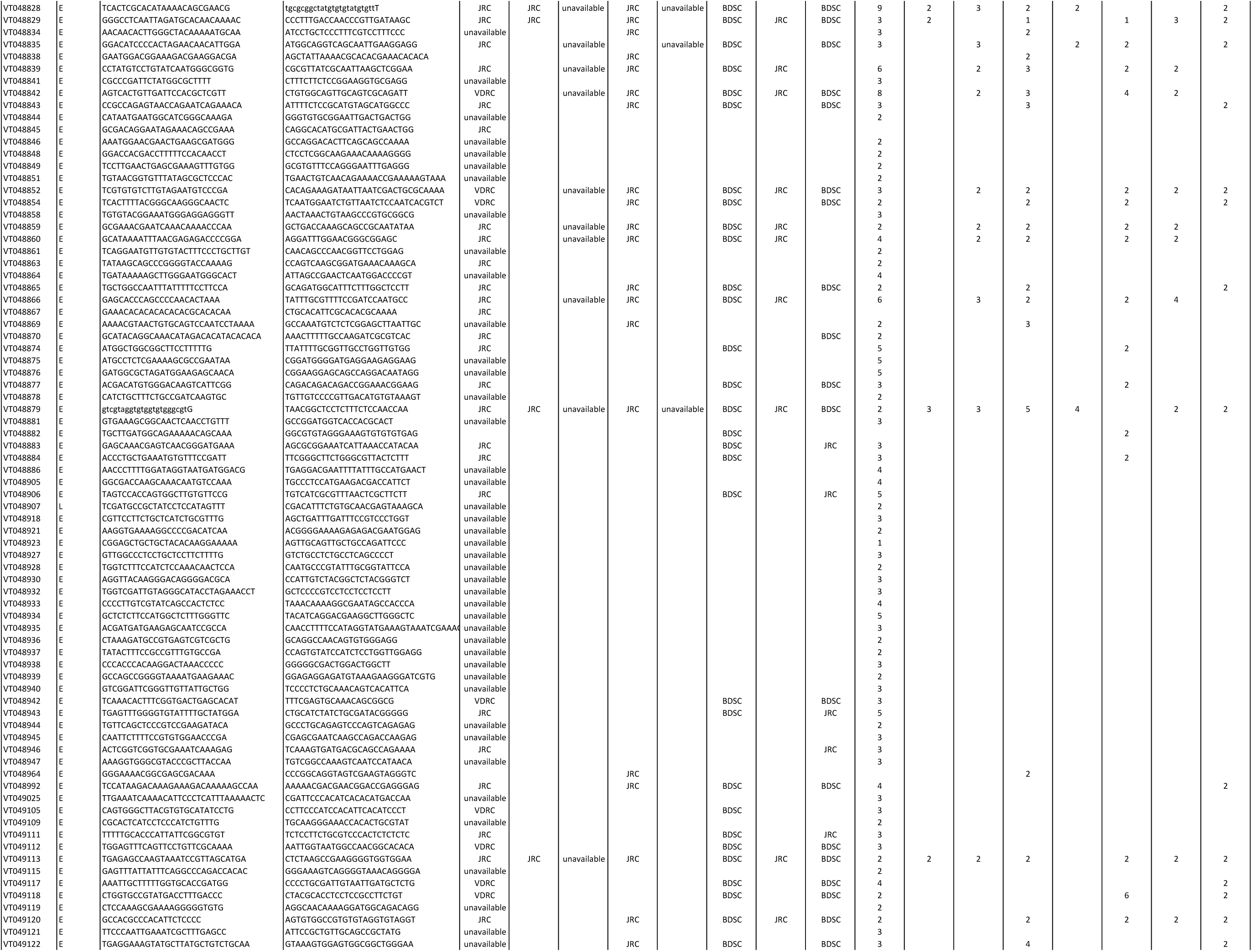
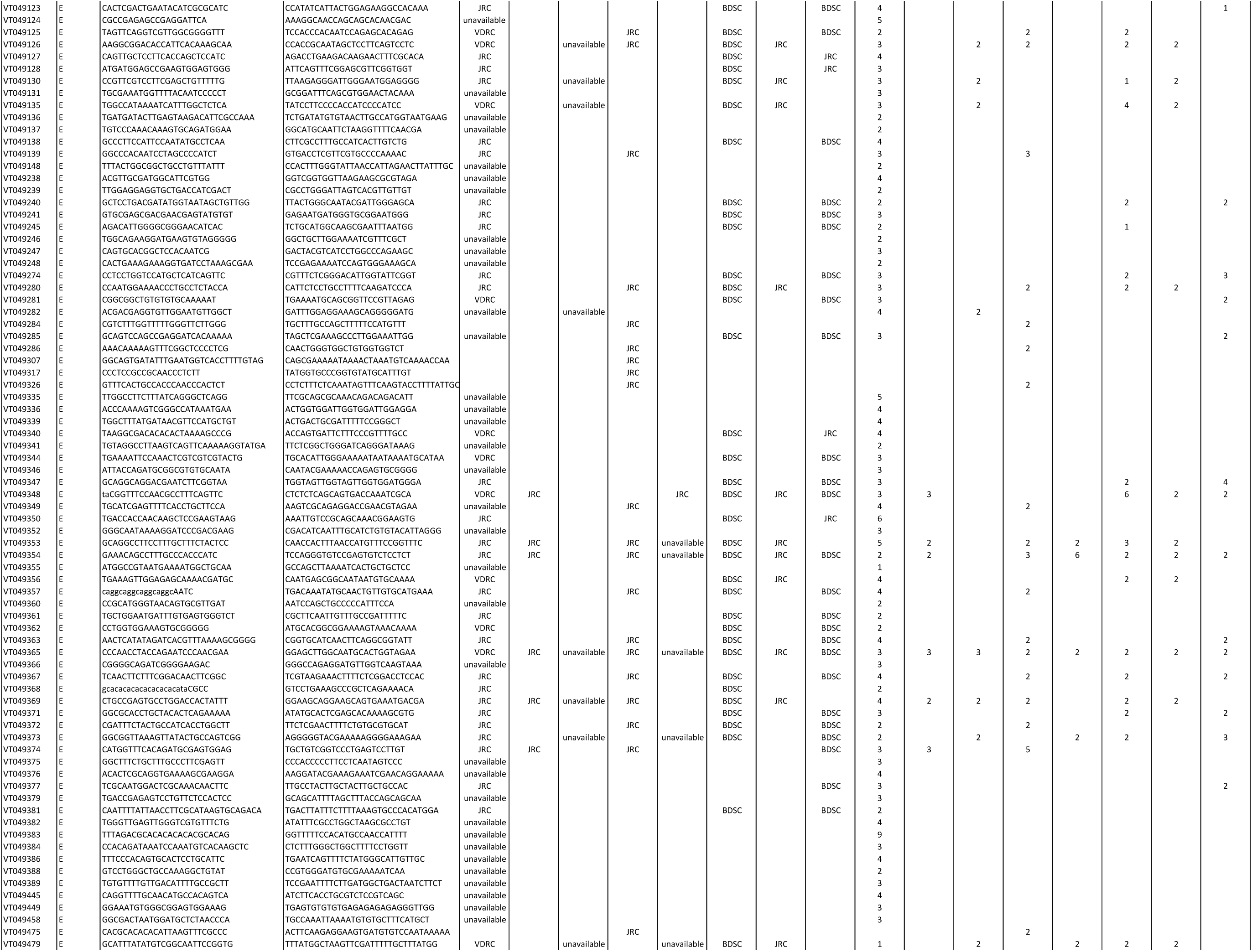
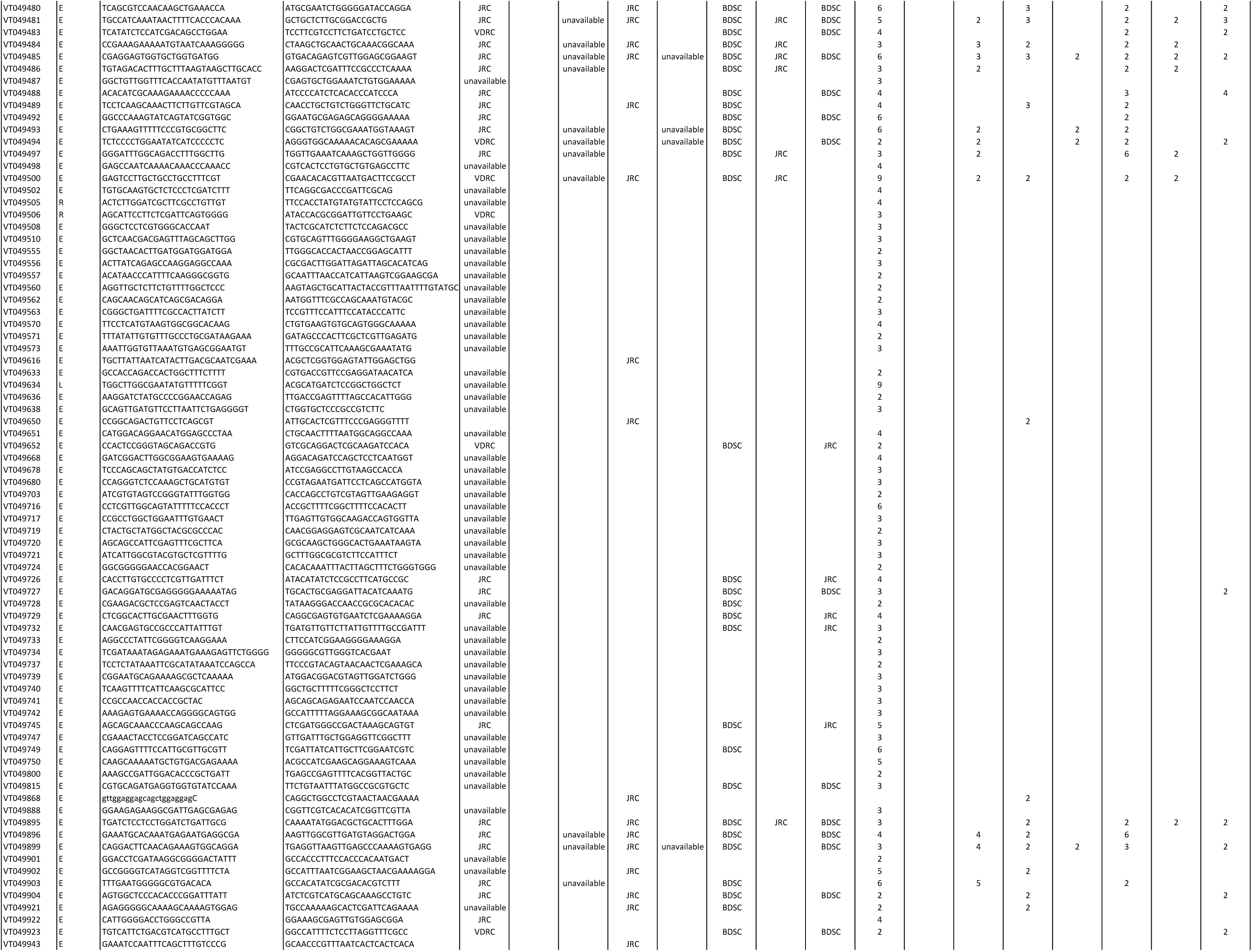
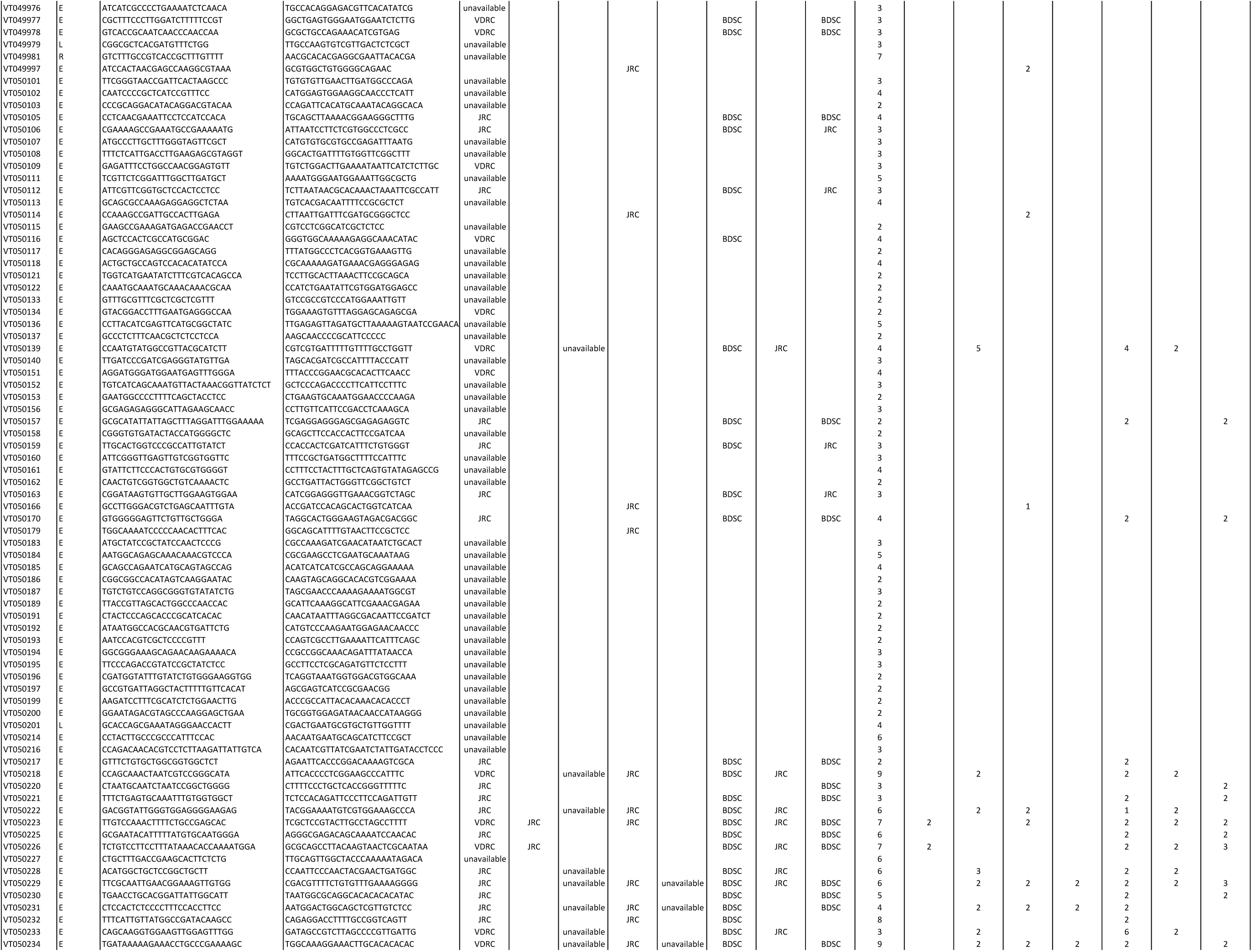
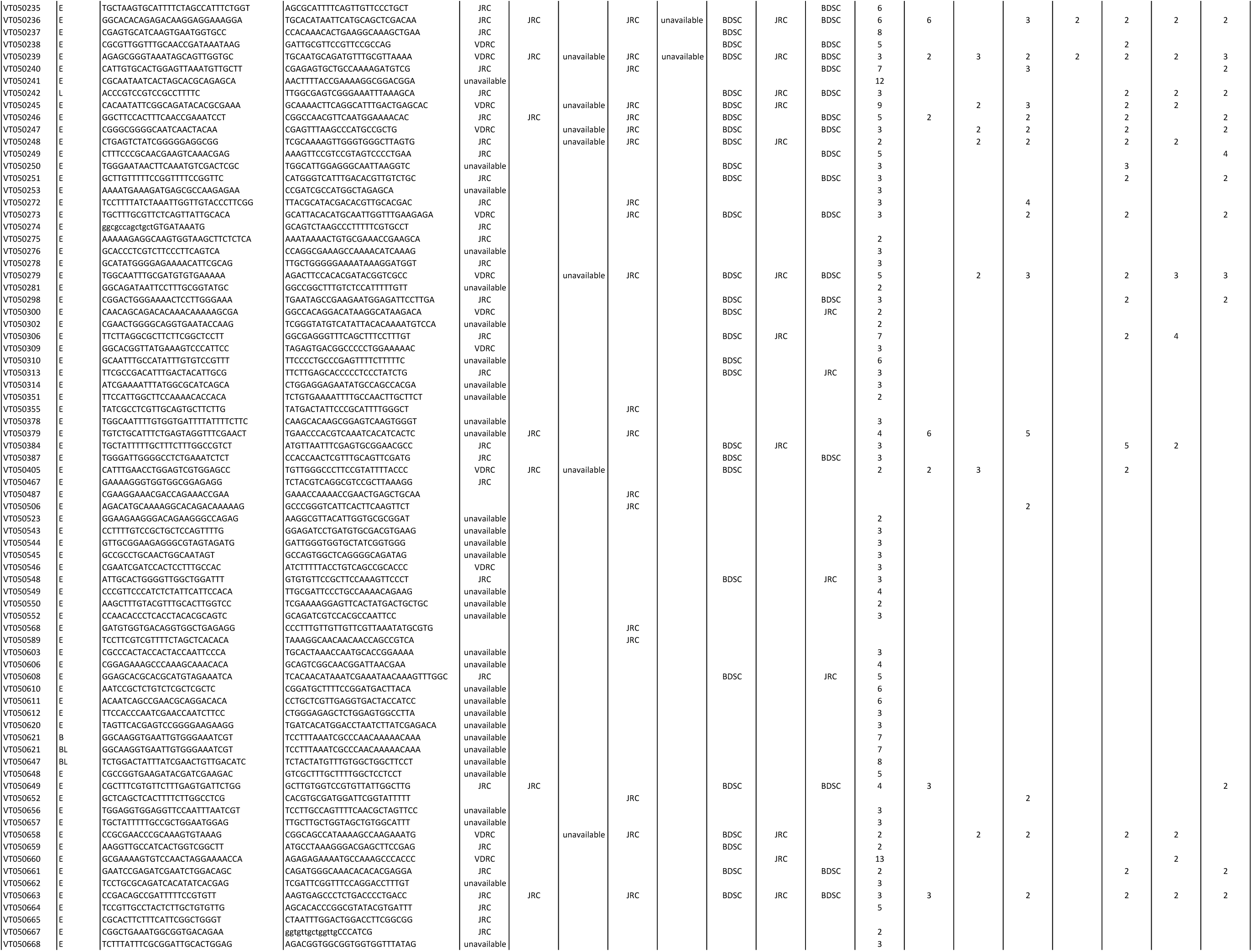
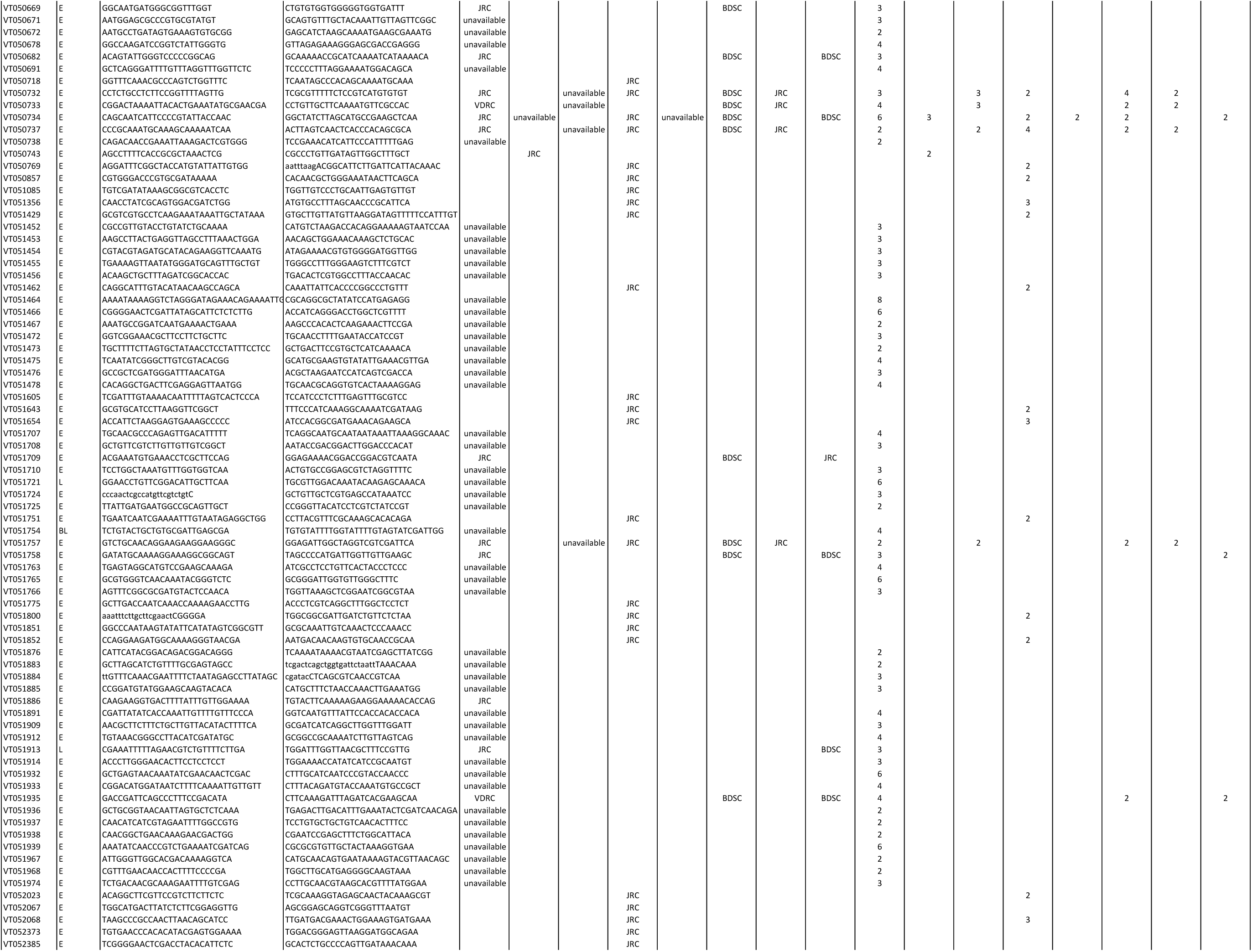
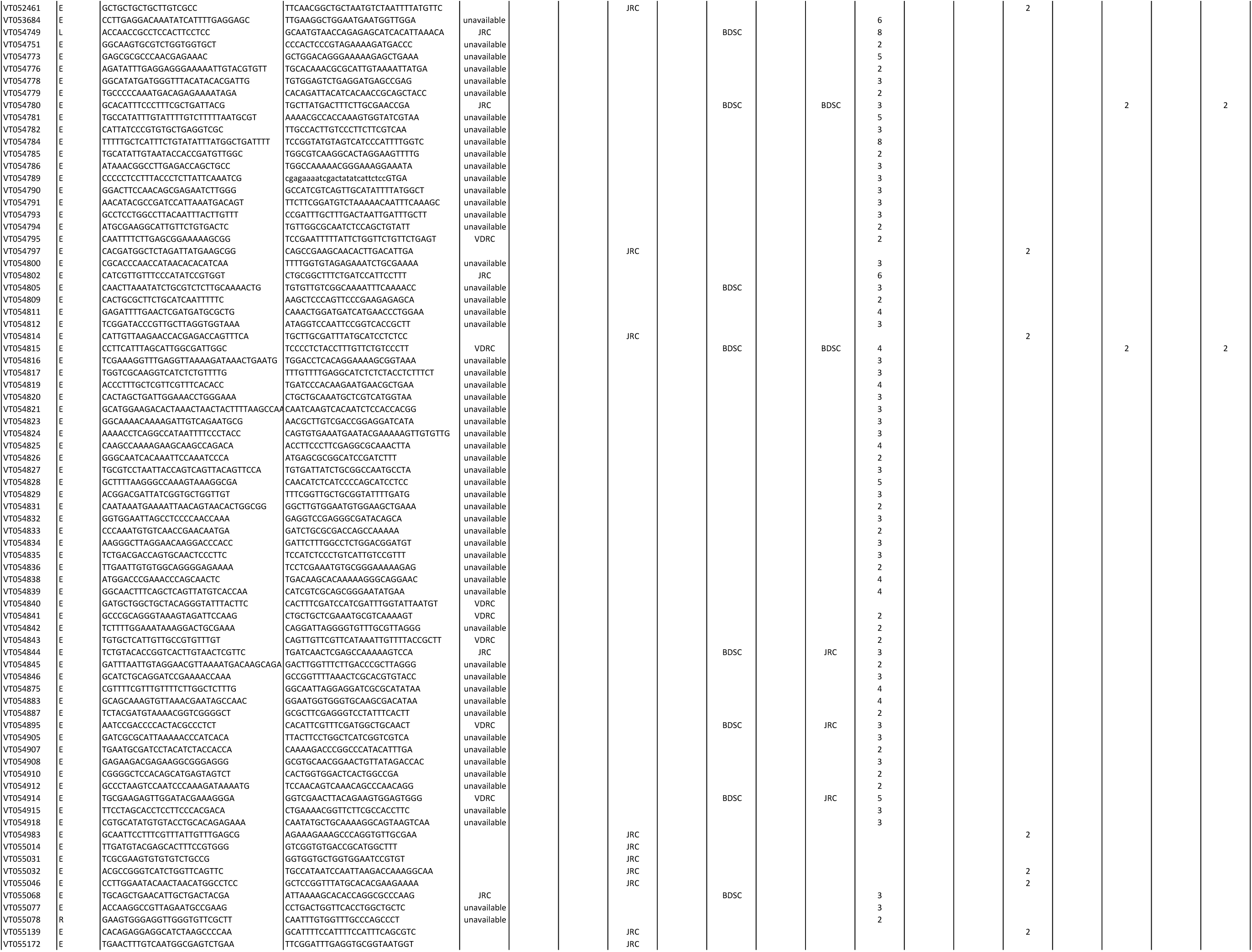
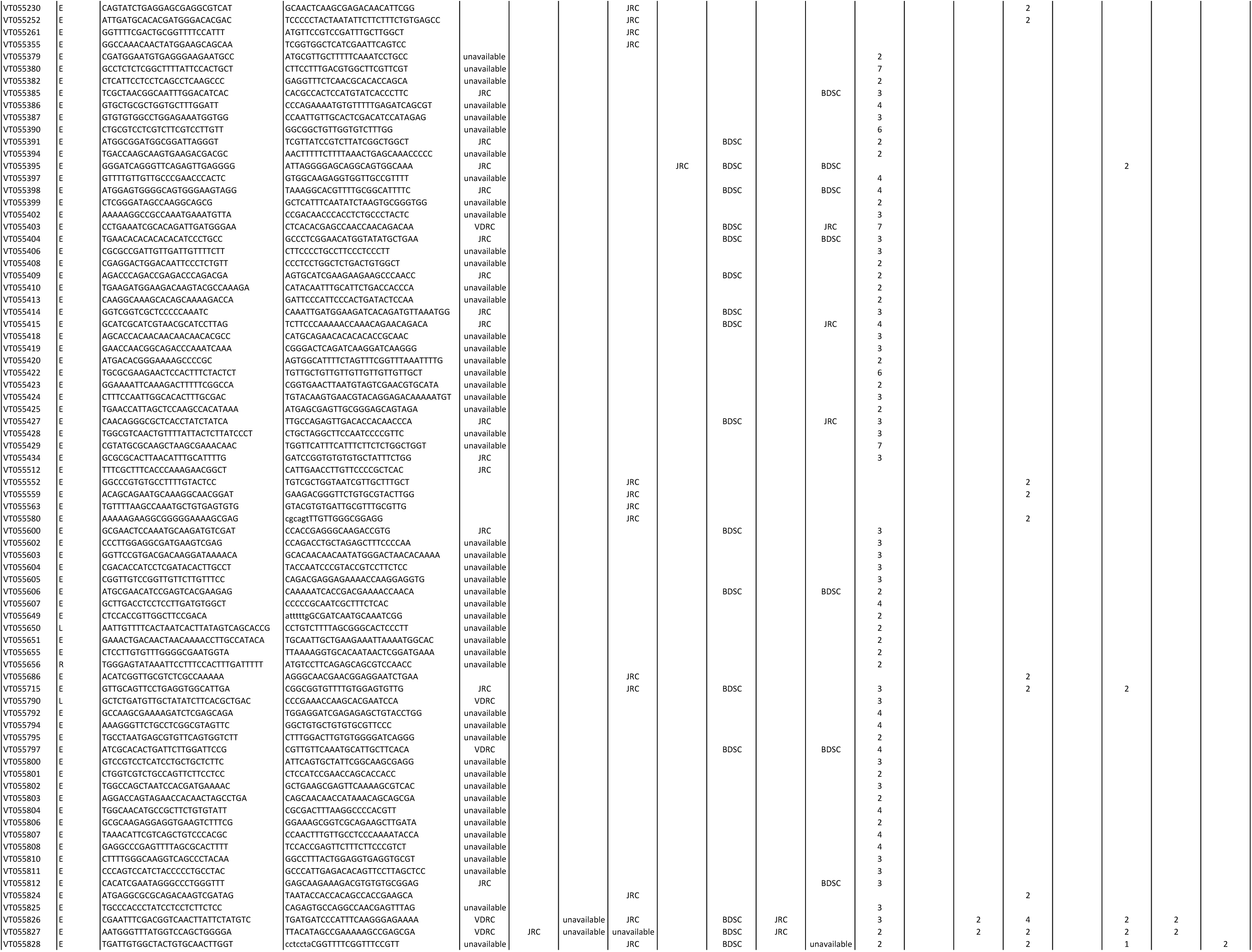
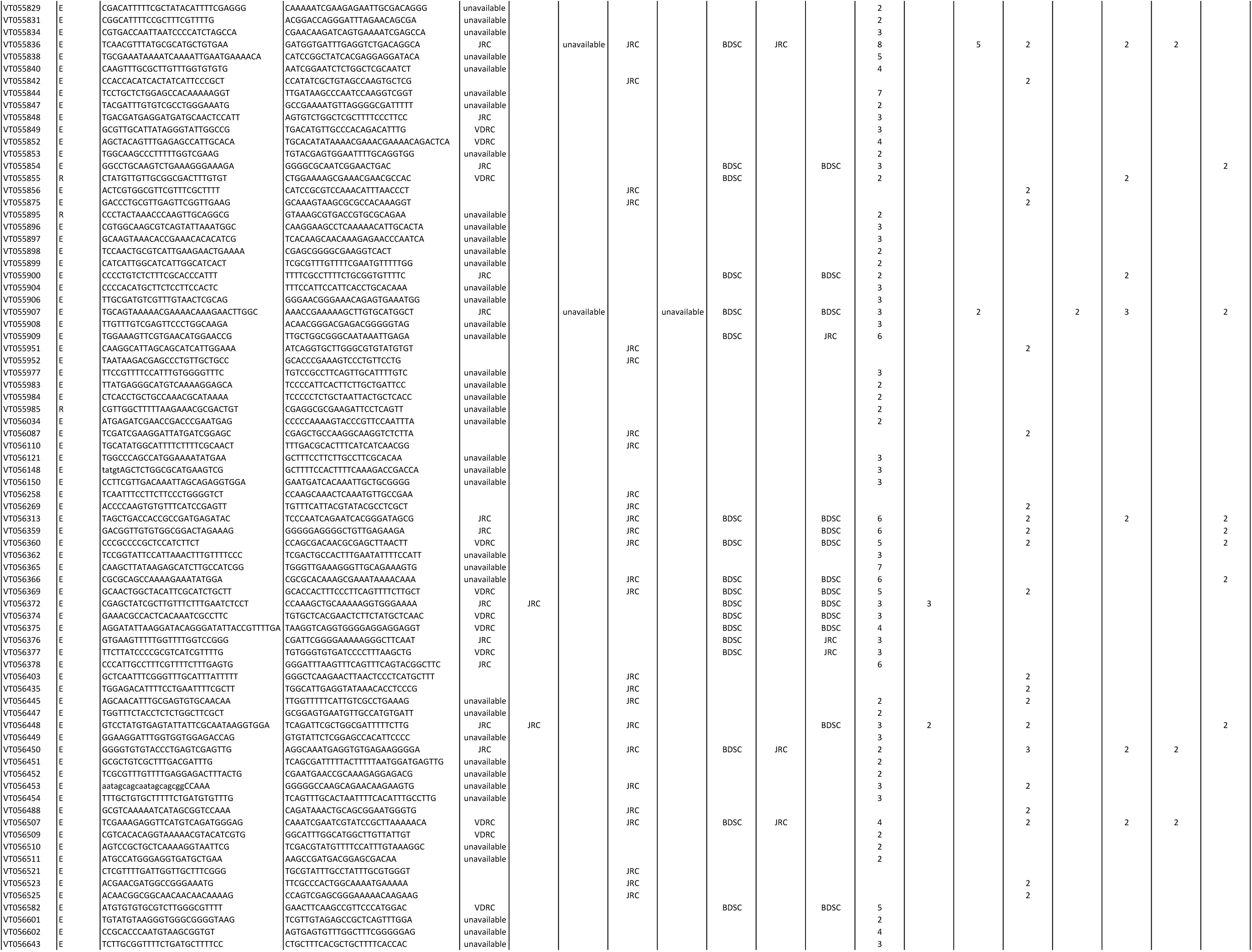
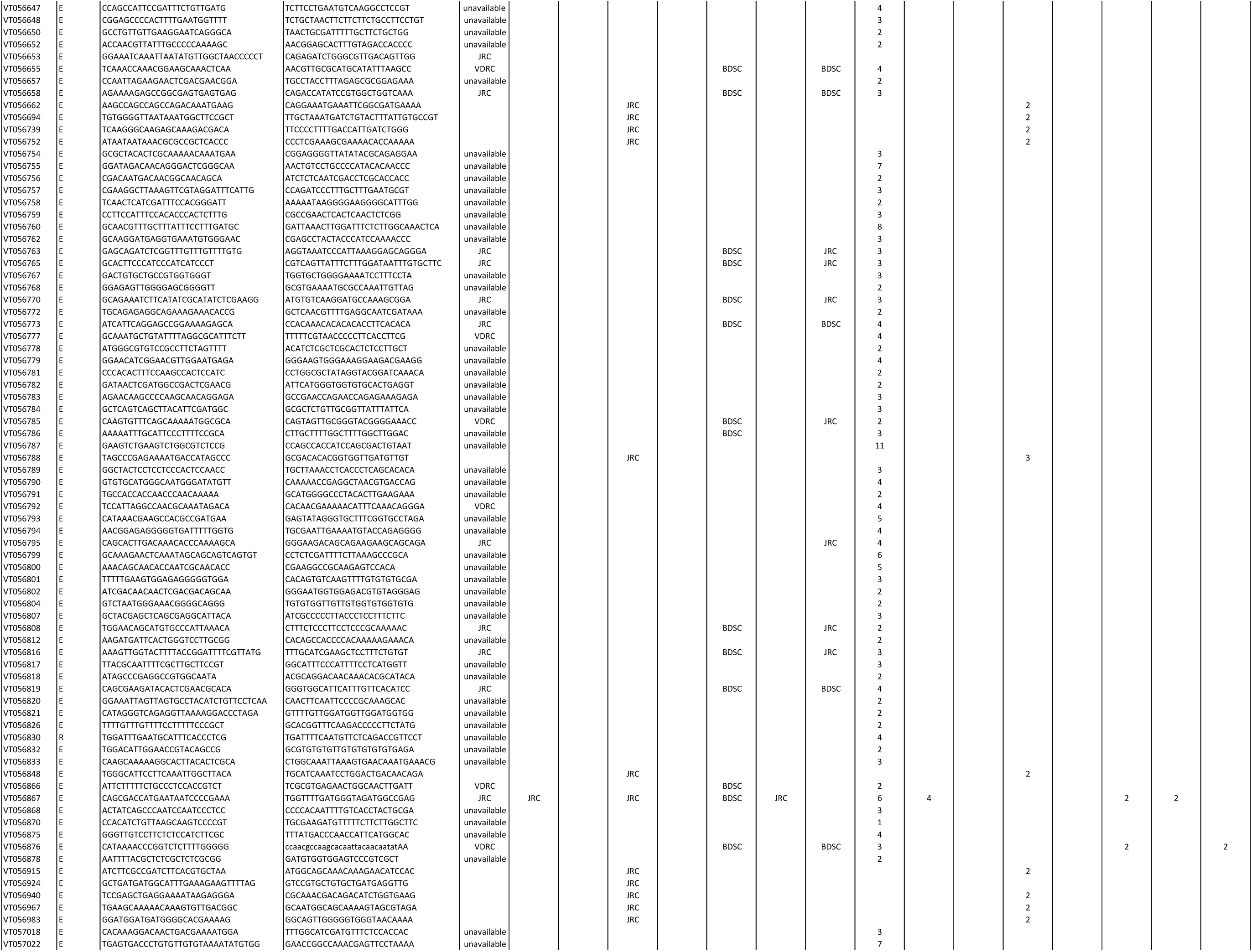
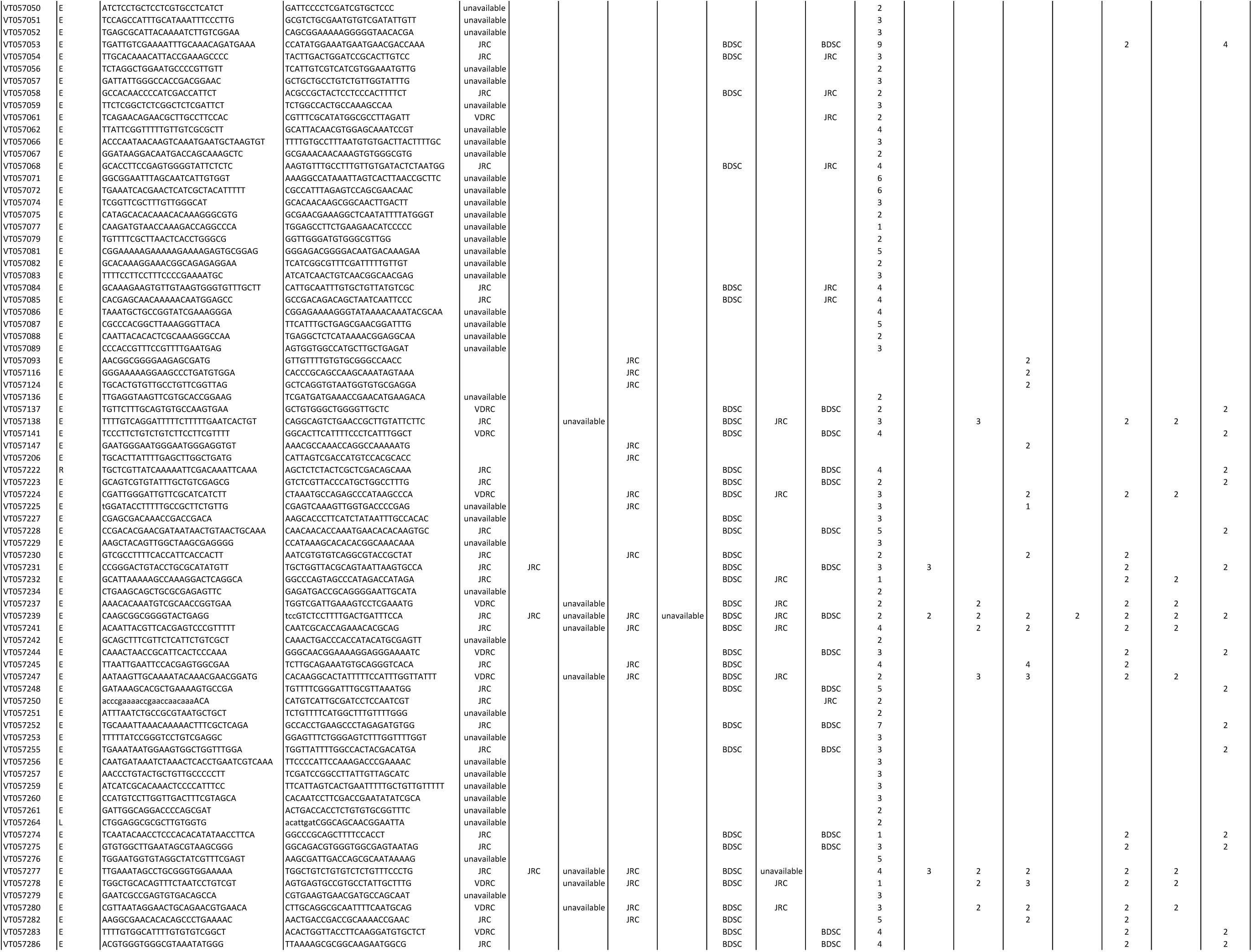
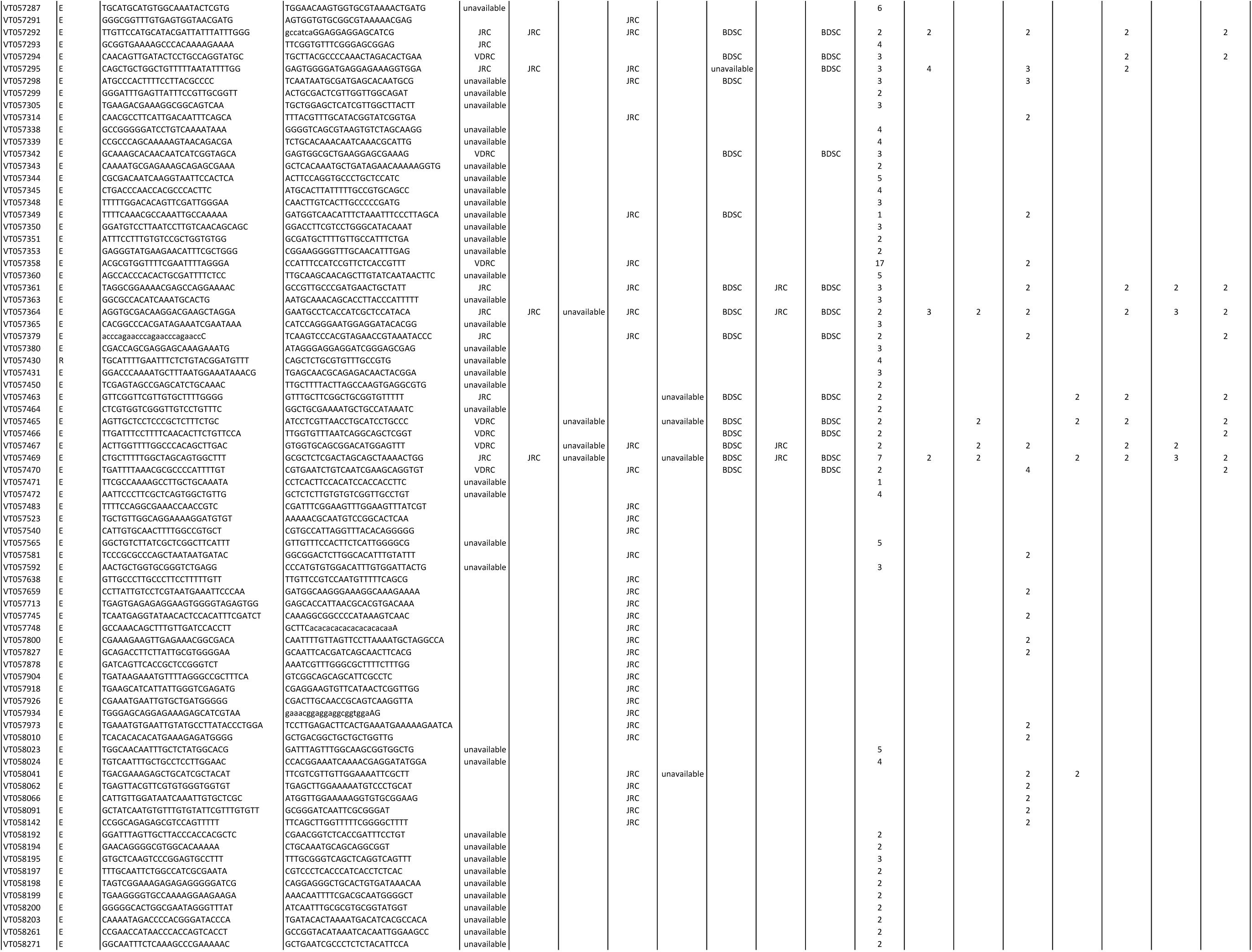
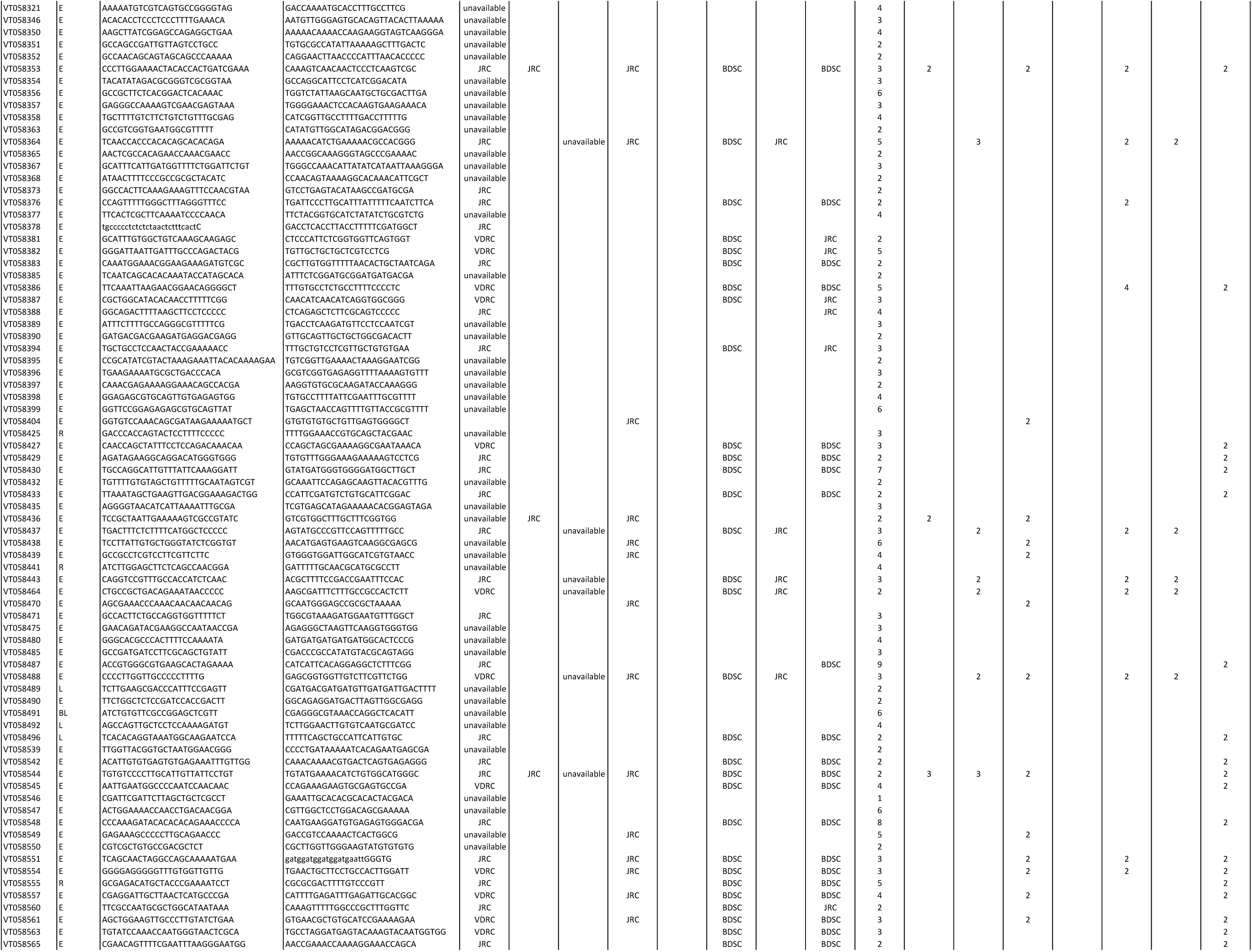
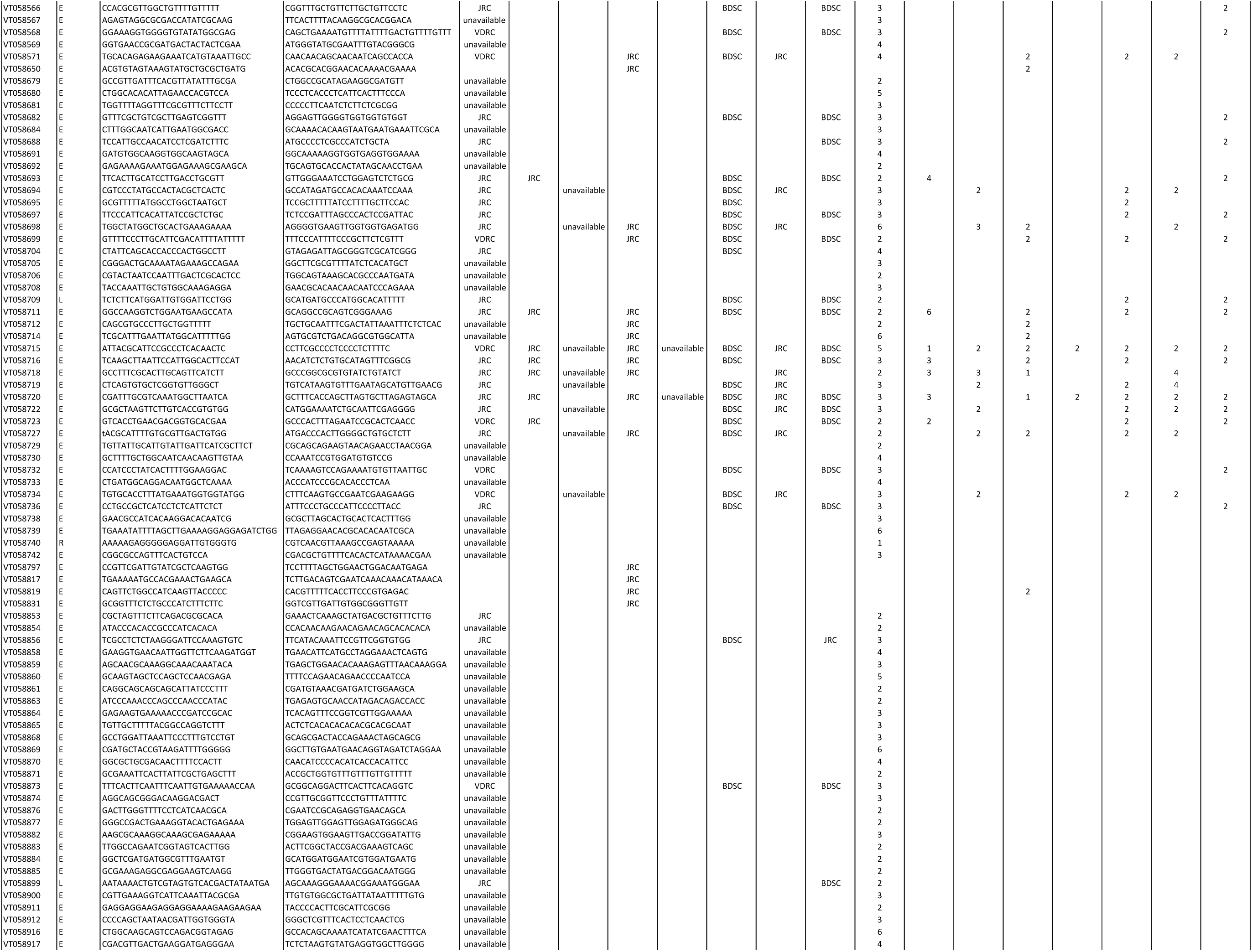
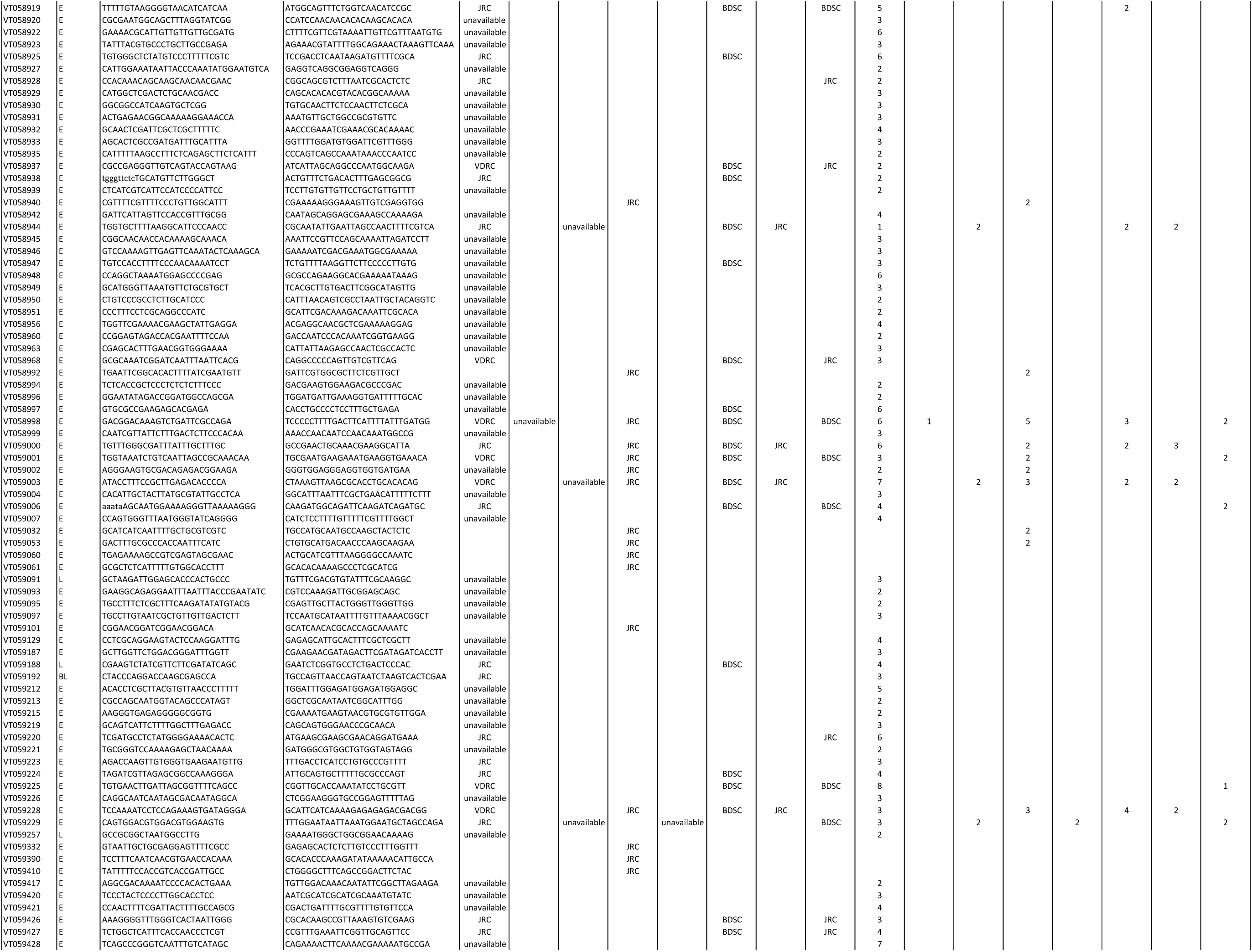
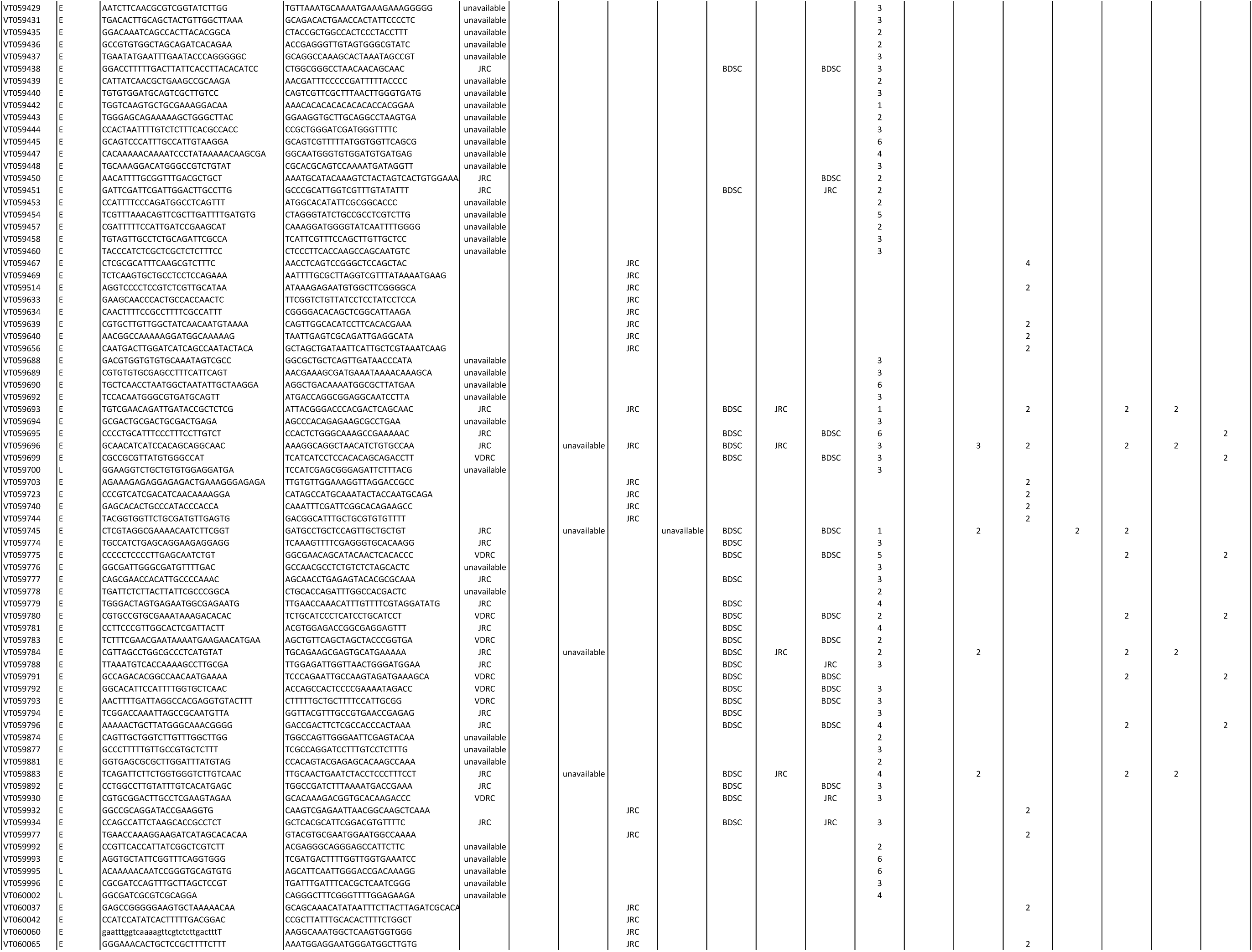
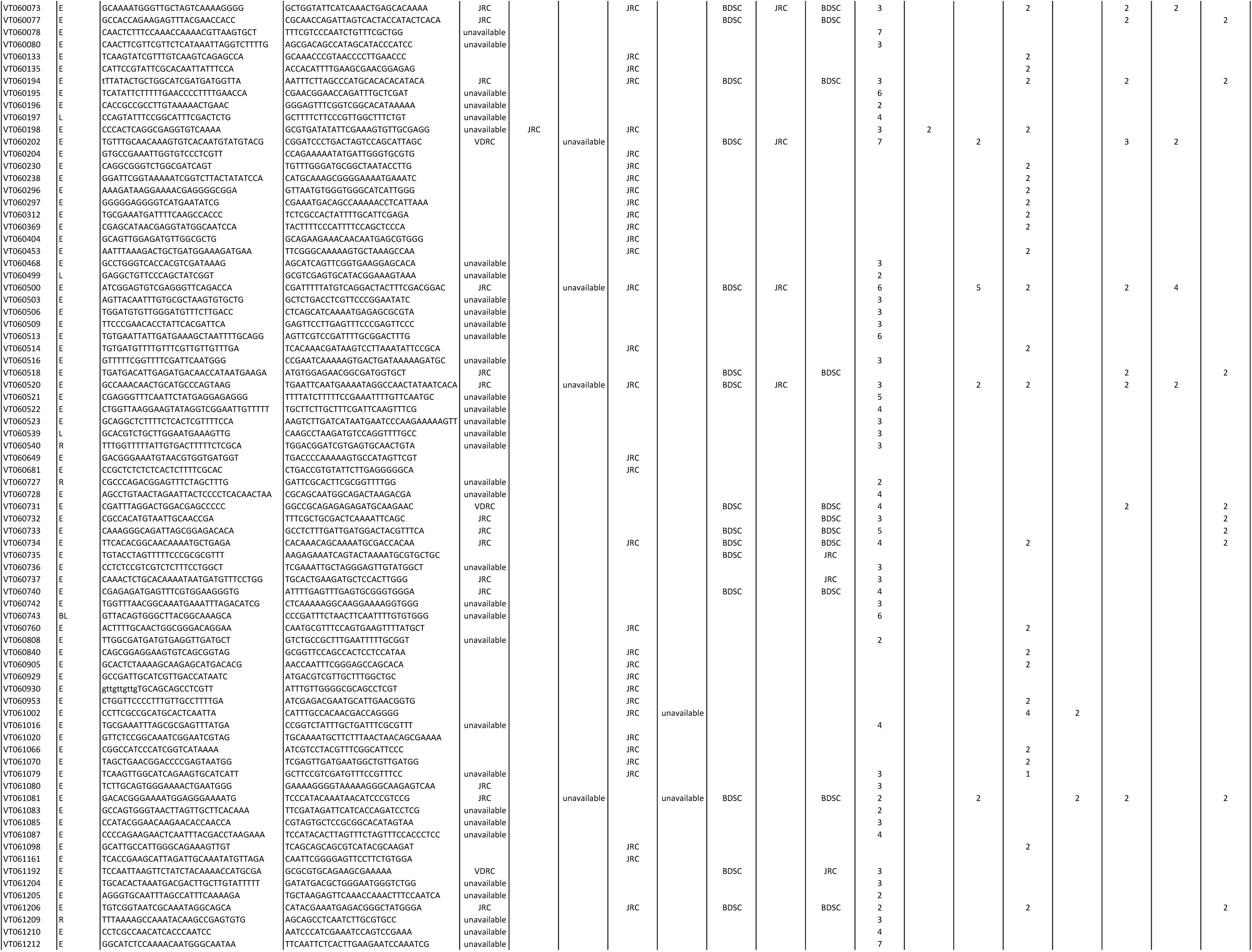
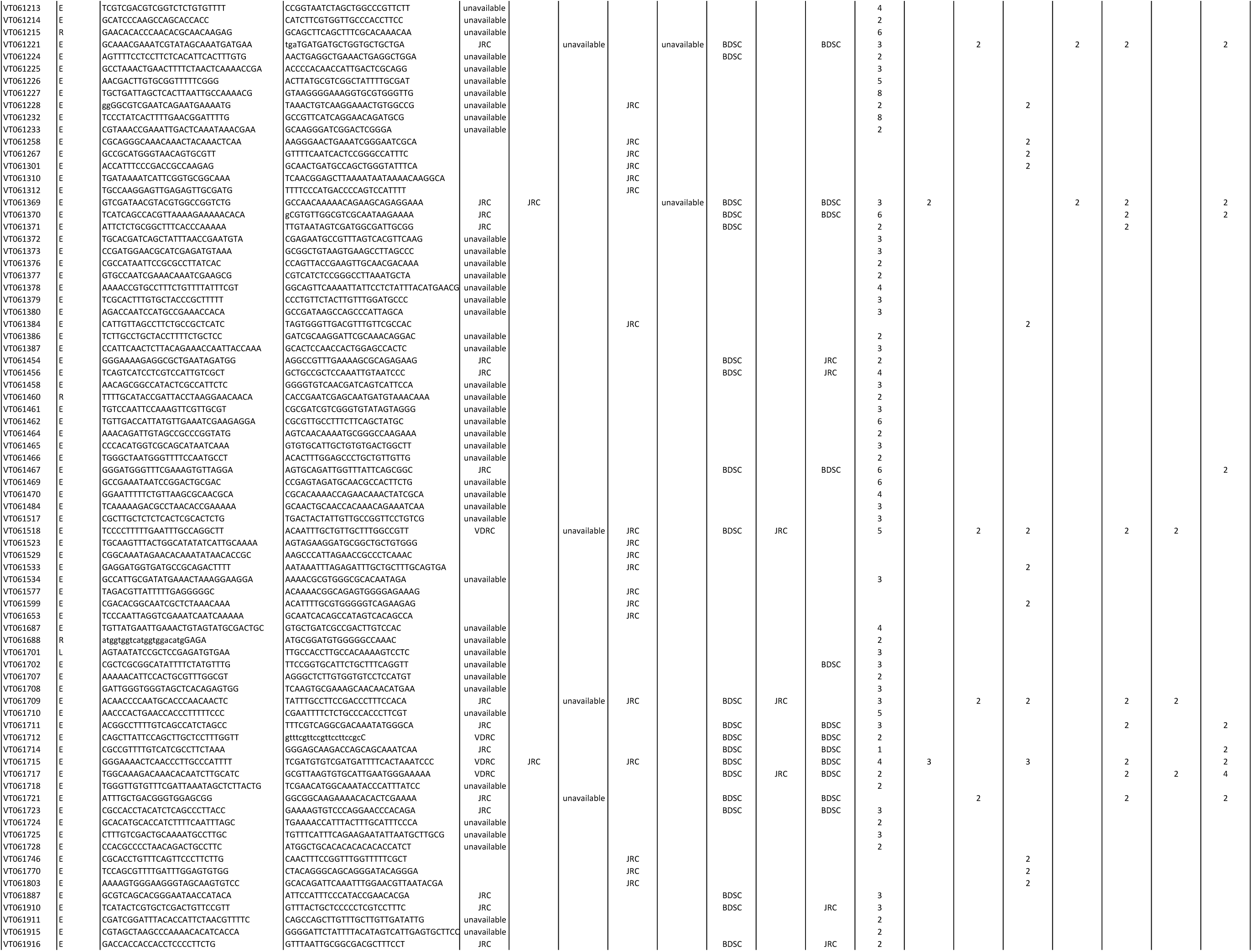
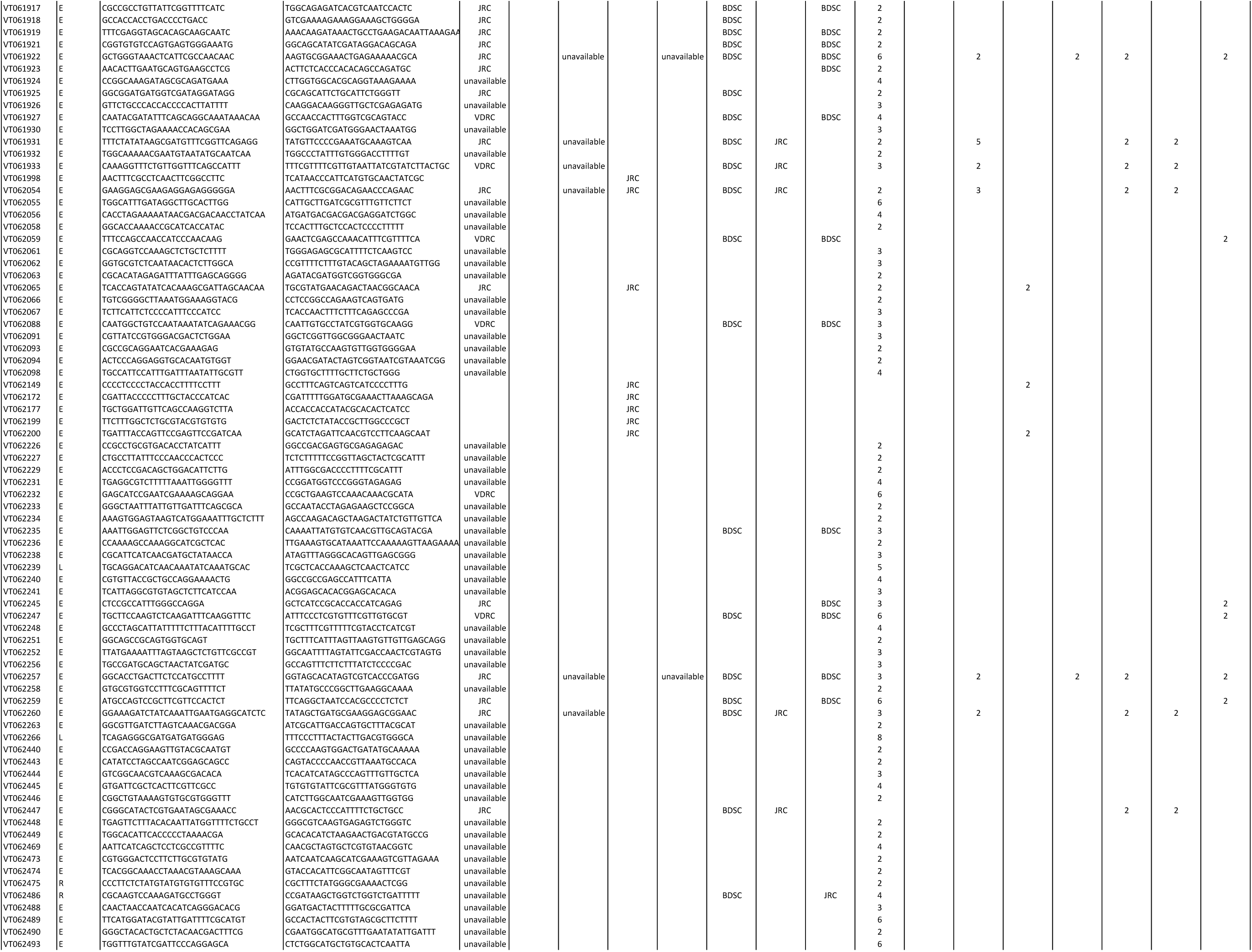
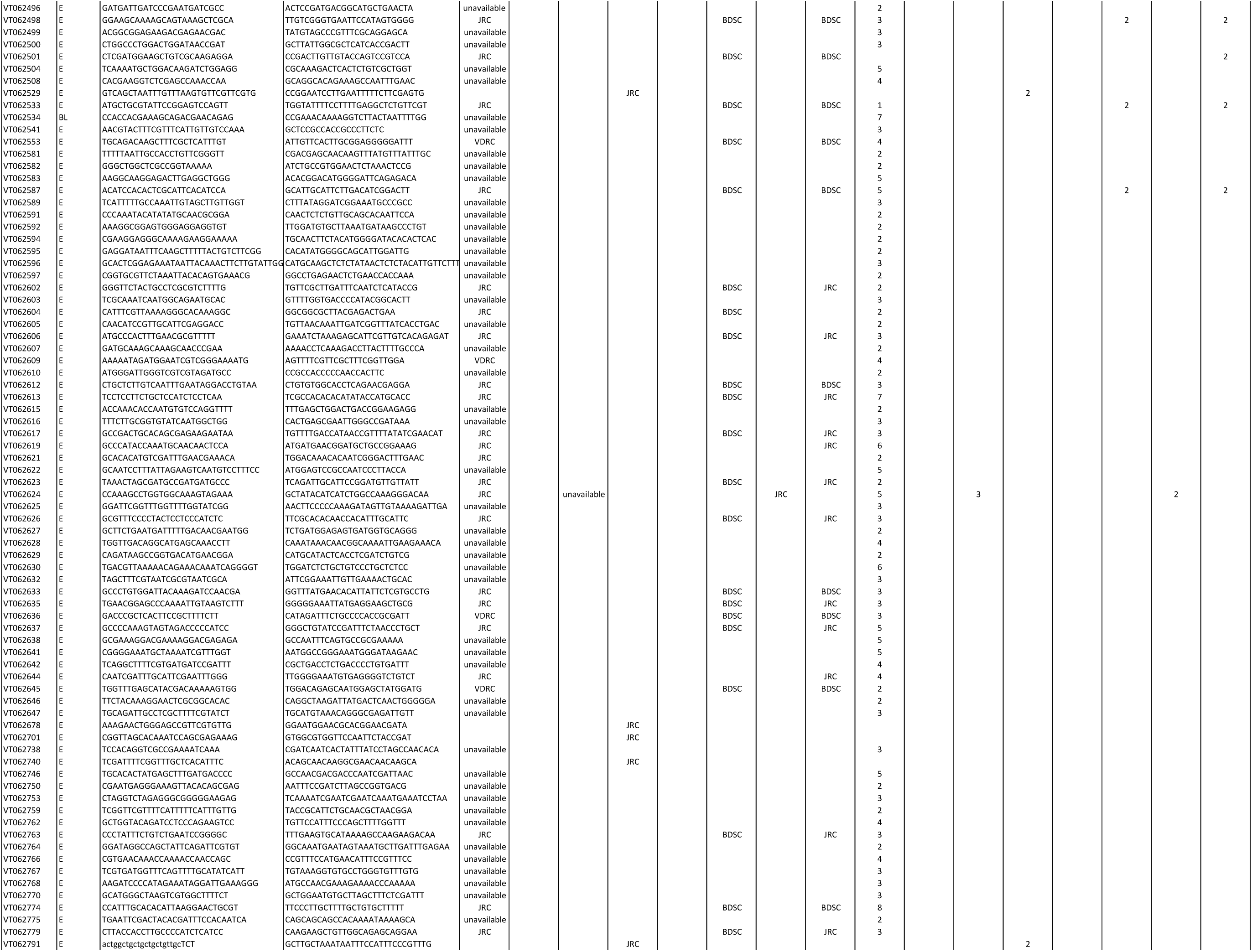
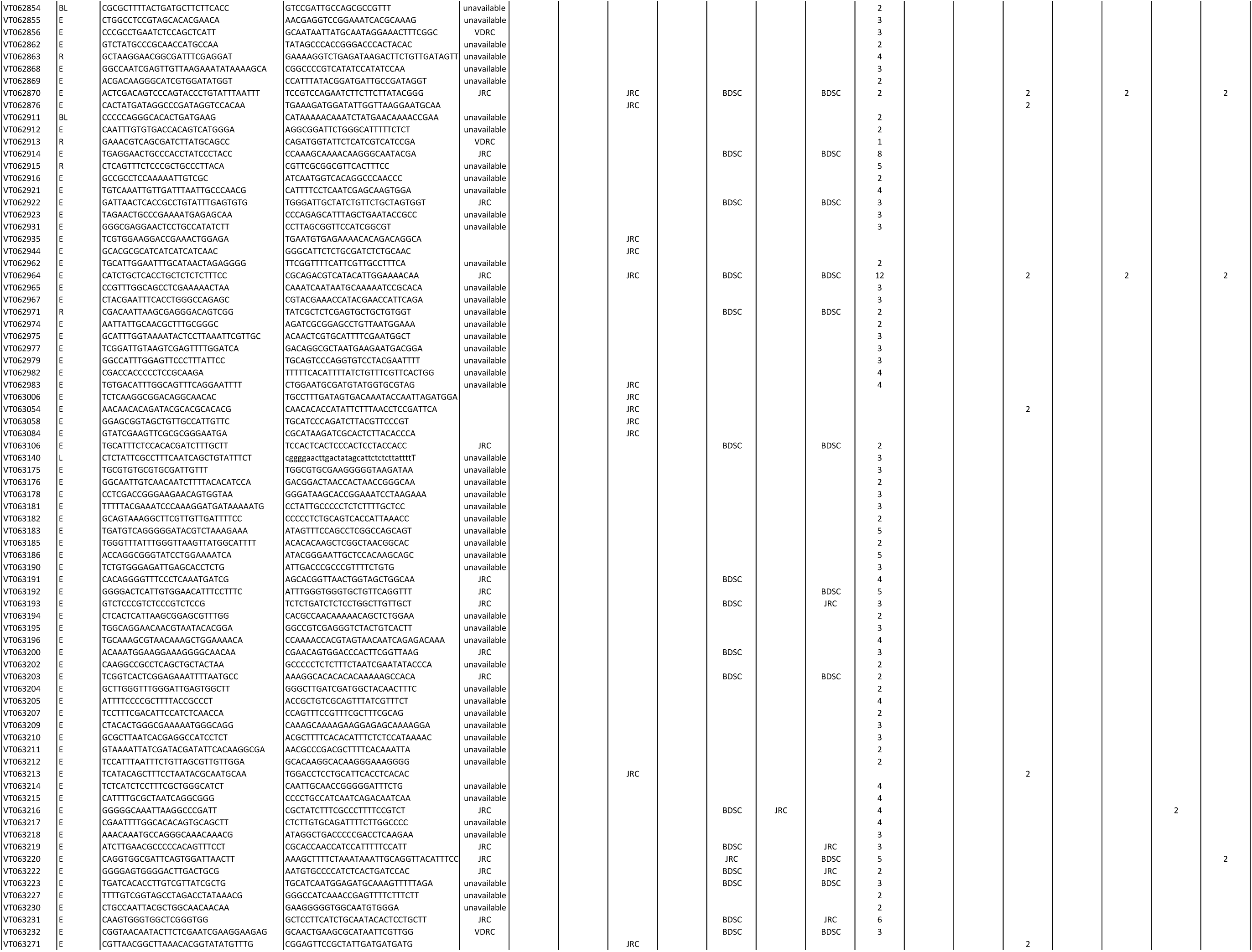
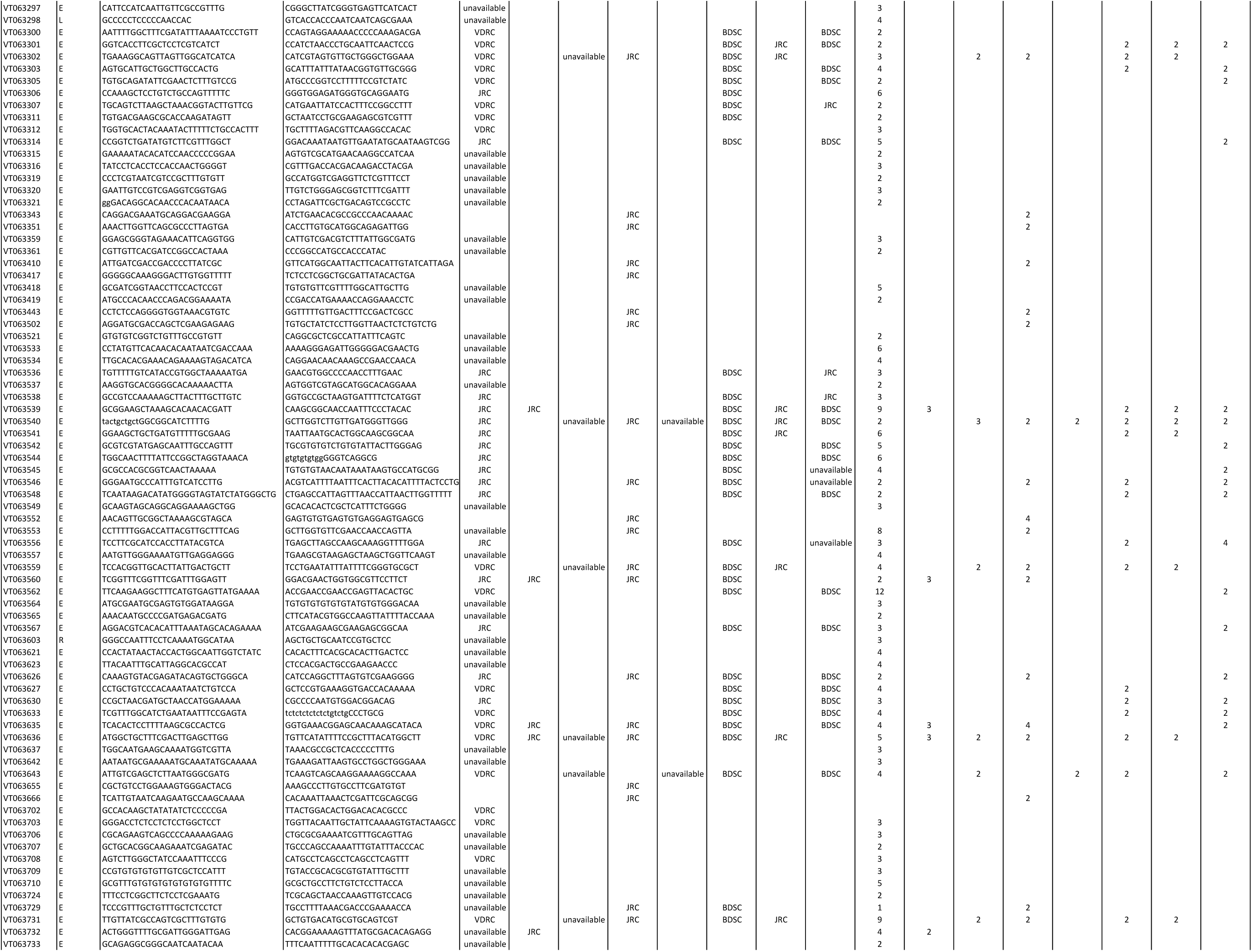
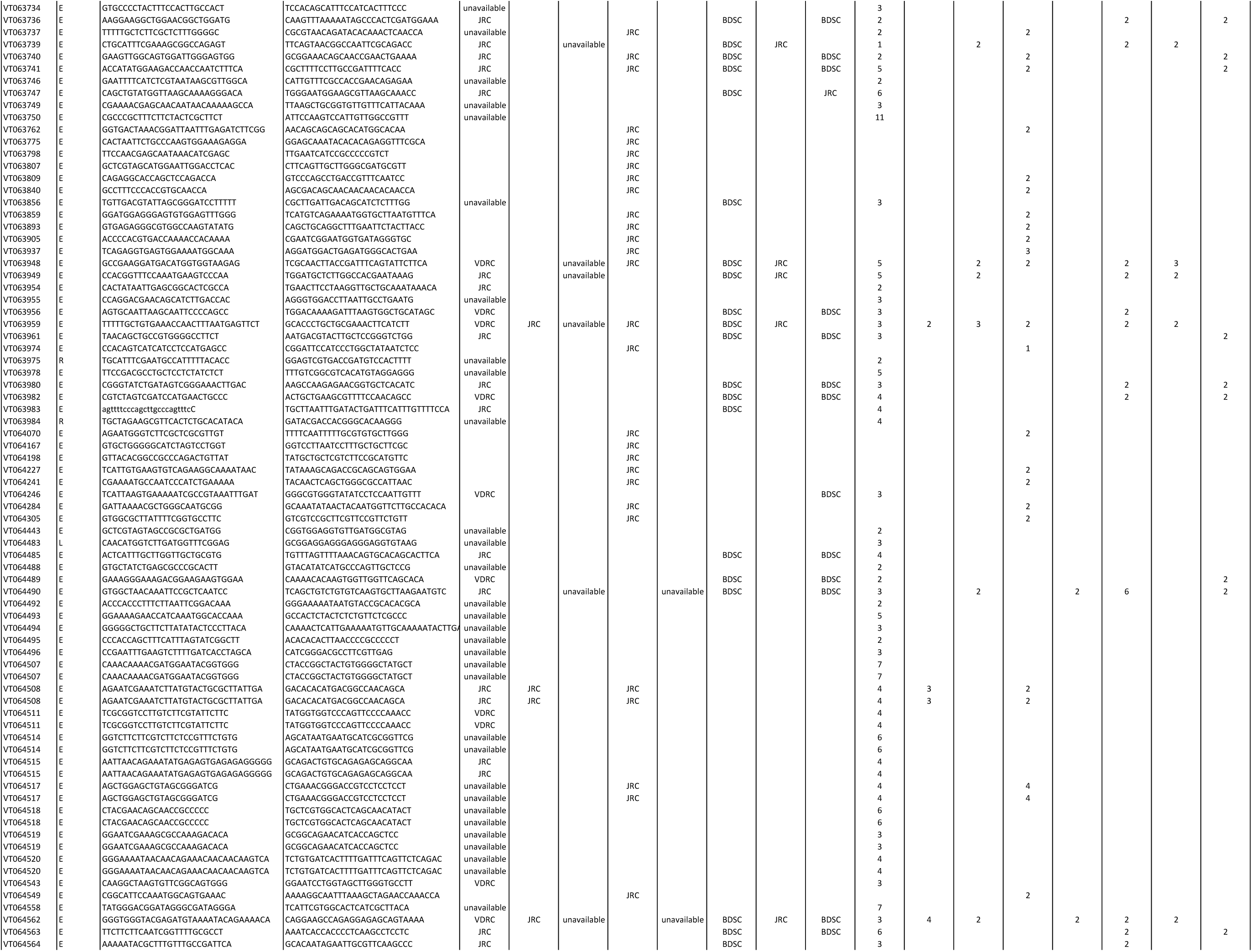
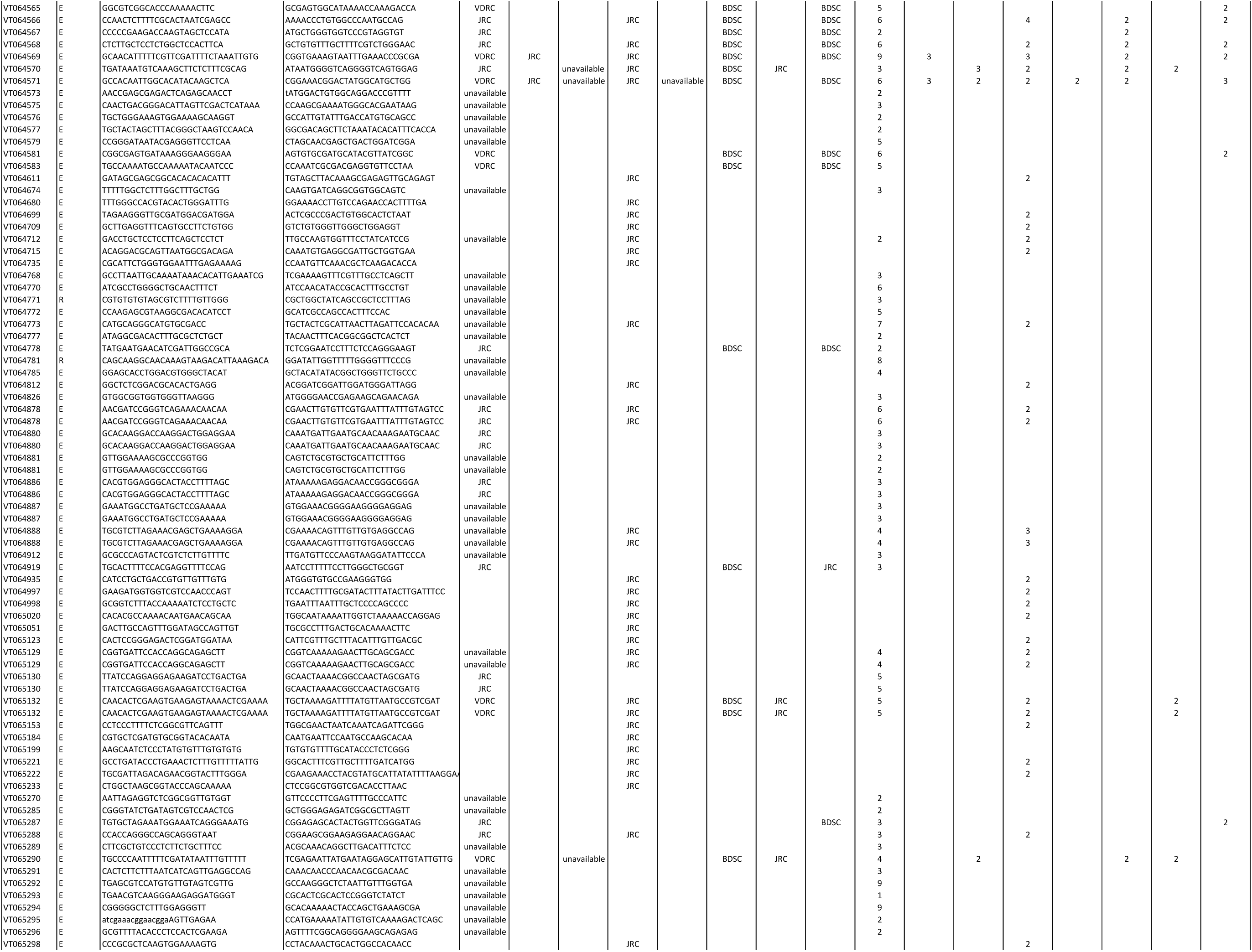
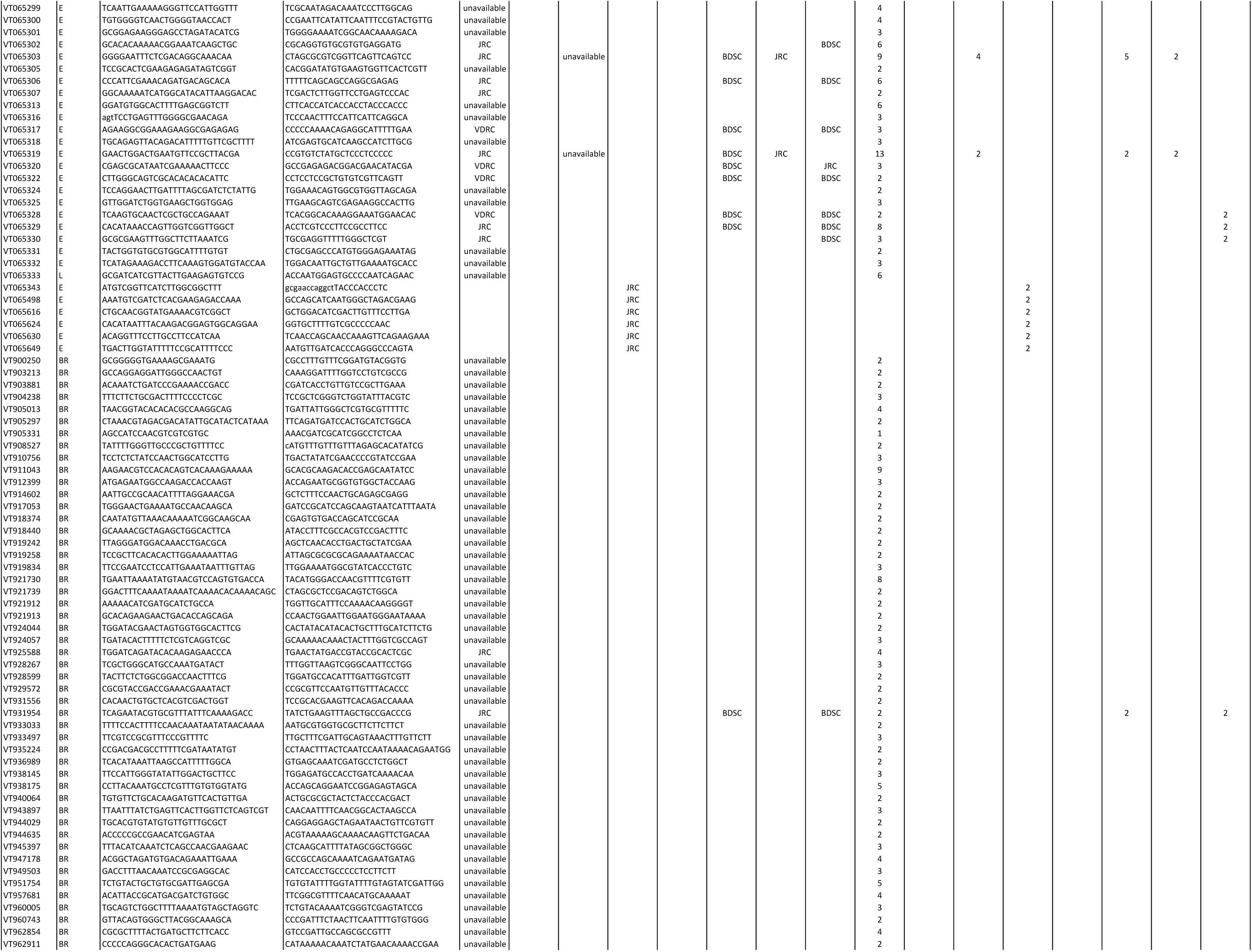
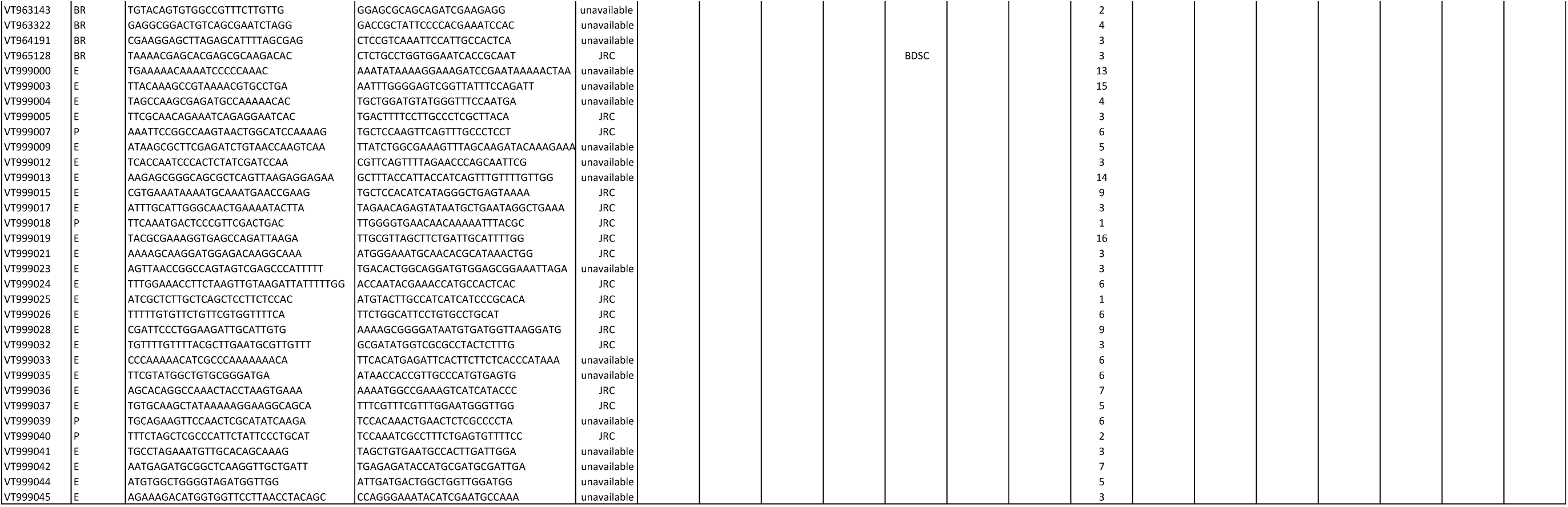
VT lines and images. CRM (cis-regulatory module) type: E, enhancer; L, promoter facing leftward, R, promoter facing rightward. Stock availability: BDSC, Bloomington Drosophila Stock Center; VDRC, Vienna Drosophila Resource Center; JRC, Janelia Research Campus; unavailable, stock no longer publically maintained. Availability according to respective stocklists on Oct 1, 2017.

## Acknowledgements

We thank the following for their contributions to the generation of the VT lines and/or images: Salil Bidaye, Luisa Deszcz, Balazs Erdi, Michaela Fellner, Angela Graf, Attila György, László Hunor Hajdu, Martin Kinberg, Christopher Masser, Magdalena Mosiolek, Thomas Peterbauer, Elena Popowich, Anton Russell, Alex Stark, Johanna Trieb, Sandor Urmosi-Incze, Stefanie Wandl, Yang Wu, Amina Zankel, and Svetlana Zorinyants. We also thank Katja Bühler and colleagues at VRVis for the BrainGazer software package used to manage the imaging pipeline and Karen Hibbard, Evgeny Kvon, Todd Laverty, Lisa Meadows and Cahir O’Kane for comments on the manuscript. Research at the IMP is supported by Boehringer Ingelheim GmbH.

## Methods

### VT line generation

The non-coding, non-repetitve regions of the *Drosophila* genome were tiled into ∼2kb fragments (Kvon et al., 2014) and individual tiles ranked for cloning according to various criteria considered predictive of a neuronal enhancer or promoter, such as proximity to predicted or known neuronal genes and the presence of conserved cis-regulatory motifs. Selected tiles were amplified by PCR from *w*^1118^ genomic DNA and cloned into the entry and transformation vectors as previously described (Pfeiffer et al., 2008; 2010). Germline transformation of *attP2* or *attP40* hosts (Groth et al., 2004) was performed either in house or by BestGene Inc. (www.thebestgene.com).

### Immunohistochemistry and confocal imaging

Flies were dissected, stained and registered to the T1 template as described in (Yu et al., 2010). In brief, brains were dissected in phosphate buffered saline (PBS), fixed in 4 % paraformaldehyde in PBS with 0.1 % TritionX100 and subsequently blocked in 10 % normal goat serum (Gibco Life Technologies). Brains were incubated in primary and secondary antibodies for 36 hours at 4°C and washed in PBS with 0.3% TritionX100 for 48 hours at 4°C. Fly brains were mounted in Vectashield (Vector Laboratories). The following antibodies were used: rabbit polyclonal anti-GFP (1:5000, TP401, Torrey Pines) and mouse monoclonal anti-Bruchpilot (1:20, nc82, Developmental Studies Hybridoma Bank) primary, and Alexa Fluor 488 and 568 conjugated secondary antibodies (1:500 to 1:1000, Invitrogen Life Technologies). Images were acquired with Zeiss LSM700 confocal microscopes equipped with 25x/0.8 plan multi-immersion objectives in a single-track mode with 0.5x0.5x1 μm voxel size. Images were aligned by nonrigid registration using the nc82 staining pattern as reference channel.

